# Live-Cell Covalent Profiling Reveals Principles of RNA-Small Molecule Recognition across the Human Transcriptome

**DOI:** 10.1101/2025.11.05.686775

**Authors:** Yuquan Tong, Amirhossein Taghavi, Xiaoxuan Su, Xueyi Yang, Warren Rouse, Sandra Kovachka, Jessica L. Childs-Disney, Ryuichi Sekioka, Rey Zavala, Yanjun Li, Walter N. Moss, Matthew D. Disney

## Abstract

RNA folds are abundant in mammalian cells yet poorly characterized as small-molecule targets. We present a scalable, *unbiased* live-cell pipeline that maps where small molecules bind RNA across the human transcriptome and convert those binders into selective degraders. A 200-member fragment library bearing diazirine/alkyne handles yielded 23 RNA-binding candidates. Chem-CLIP-Map-Seq in MDA-MB-231 cells identified 723 RNA targets and their binding sites, revealing a strong bias toward 5′ and 3′ untranslated regions (UTRs) in mRNAs and enrichment at thermodynamically stable structures, with limited binding to non-coding RNAs. Expression level and local stability contributed to engagement. An integrated machine-learning model trained on multiple fingerprints distinguished binders from non-binders, and highlighted chemotypes and physicochemical features that favor RNA recognition. Four fragments were converted to RiboTACs; despite broad binding, cleavage was highly selective, with X1-RiboTAC degrading *MPP7* and *SSC4D* mRNAs in an RNase L-dependent manner and reducing their protein levels. A competitive profiling workflow quantified in-cell target occupancy and guided optimization of the RNA-binding module to reprogram selectivity: an X1 derivative produced an *MPP7*-selective RiboTAC that lowered MPP7 mRNA levels and suppressed breast-cancer cell migration, while sparing *SSC4D* transcripts. This end-to-end framework, including transcriptome-wide mapping, data-driven rules, and tunable degradation, establishes practical principles for ligandable RNA sites in cells and enables rational design of RNA-targeted small molecules and degraders.

**TEASER:** Live-cell mapping reveals ligandable RNA sites and guides design of selective RNA degraders.

**TOC graphic:** 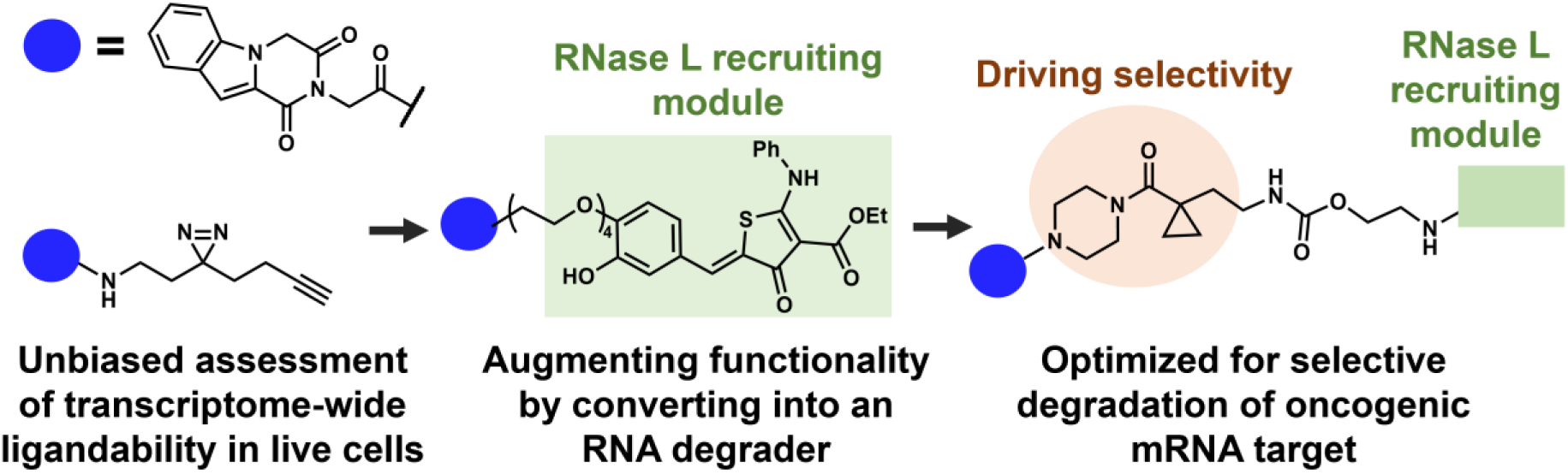

## INTRODUCTION

The biological functions of RNAs are dependent on the structures they adopt, contributing to both homeostasis and disease.^1–3^ RNA function can be inhibited by small molecules that bind to functional structures in all kingdoms of life,^4–6^ including inhibiting the conformational switching of riboswitches in bacteria,^7, 8^ impeding the self-splicing of a group II intron in fungi,^9^ and disrupting the biogenesis of human micro (mi)RNAs.^10–12^ Most bioactive RNA-targeted small molecules have been identified from target-centric approaches, while the ligandability of the entire human transcriptome remains largely unexplored. Unbiased, target-agnostic profiling that defines the molecular fingerprints of RNA-binding small molecules across the transcriptome could expand the druggability of the human genome by identifying ligandable RNA structures. That is, such data could provide a foundation for understanding the molecular recognition between RNA and small molecules in native cellular environments and thereby enable the rational design and optimization of RNA-binding small molecules in an unbiased, data-driven manner. Notably, similar efforts have been ongoing for protein targets, where the proteome has been probed with low molecular weight small molecules that cross-link to their targets (dubbed fully functionalized fragments or FFFs) and therefore probe small molecule-protein interactions.^13–15^ These data, subjected to machine learning, could accelerate future discovery of both RNA- and protein-targeted ligands (**Figure 1**).

**Figure 1.**
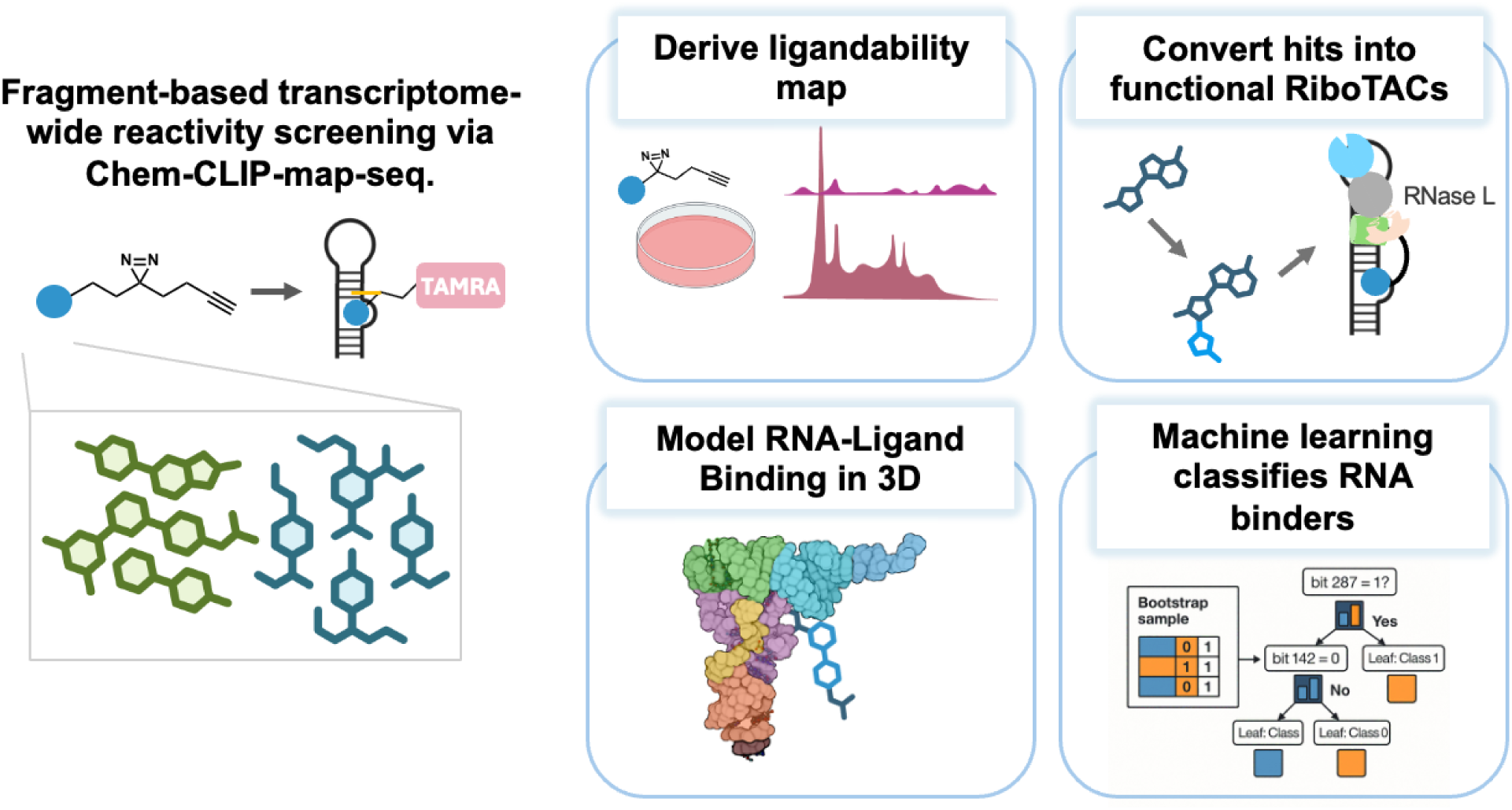
Integrated pipeline for discovering, modeling, and functionalizing RNA-binding small molecules. A chemically diverse set of diazirine/alkyne-tagged fragments is profiled against cellular RNAs by Chem-CLIP (UV-activated crosslinking) and click enrichment, generating ligandability maps across the transcriptome (top left). From these maps, hits are converted into RiboTACs by appending an RNase-L–recruiting module so that targeted RNAs are cleaved in cells (top right). In parallel, RNA–ligand binding is modeled in 3D by secondary-structure–guided RNA modeling and docking to propose binding sites and poses (bottom left). Features from active and inactive fragments are then used in a machine-learning classifier to distinguish RNA binders and to prioritize chemotypes for optimization (bottom right). TAMRA denotes a fluorescent tag used in in-gel readouts. The schematic summarizes the end-to-end workflow applied in MDA-MB-231 cells to map ligandable RNA sites, propose binding modes, and convert binders into functional degraders.

As a parallel to the fragment-based profiling assays developed for protein targets, a platform named Chem-CLIP-Map-Seq (<u>Chem</u>ical <u>C</u>ross-<u>L</u>inking and Isolation by <u>P</u>ull-down to <u>Map</u> Small Molecule-RNA Binding Sites by <u>Seq</u>uencing) has been developed to define the RNA targets bound by small molecules and their respective binding sites therein in live cells.^16–19^ In this platform, each RNA-binding small molecule is appended with a diazirine group, which forms a covalent bond with bound RNA targets upon UV irradiation, and an alkyne handle that enables pull-down and enrichment of bound targets via copper catalyzed azide-alkyne cycloaddition (CuAAC) “click” chemistry^20^ to azide functionalized beads. The binding site of small molecules within their RNA targets are identified from subsequent RNA-seq analysis as “RT-stops”,^17–19, 21^ where reverse transcription is halted by the cross-linked small molecule. In this study, a streamlined pipeline was developed to profile 23 RNA-binding small molecules in live cells, particularly the triple negative breast cancer (TNBC) cell line MDA-MB-231, identifying a total of 723 unique RNA targets bound by small molecules as well as their respective binding sites.

As previous studies have shown that not all small molecule binding events elicit a biological response, that is, they are biologically inert,^22^ the functionality of the identified RNA binders was augmented by tethering them to an RNase L-recruiting module to create heterobifunctional molecules that induce cleavage of the RNA targets.^23^ Transcriptome-wide profiling of these ribonuclease targeting chimeras (RiboTACs) was completed in two CRISPR-modified MDA-MB-231 cell lines – one where RNase L was knocked down (ablating or reducing cleavage of *bona fide* targets) and the other as a control cell line that expresses RNase L. These studies identified two RNA targets cleaved in an RNase L-dependent manner by the same molecule, **X1-RiboTAC**, MAGUK P55 Scaffold Protein 7 (*MPP7*) and Scavenger Receptor Cysteine Rich Family Member with 4 Domains (*SSC4D*). Optimizing the structure of the **X1** RNA-binding module reprogrammed its selectivity to cleave the oncogenic target *MPP7* with no detectable effect on the non-oncogenic target *SSC4D*.

Collectively, these studies lay a foundation to identify the RNA targets of small molecules in a massively parallel and target-agnostic fashion (**Figure 1**). Broadly, these rich datasets also contribute to deciphering the codes in RNA structures that can be bound by small molecules, required for the design and optimization of RNA-targeted precision medicines.

## RESULTS

### Profiling an RNA-focused compound library for binding human RNAs *in vitro*

A chemically diverse library of 200 low molecular weight (373 ± 27 Da), compounds was designed based on similarity to known RNA-binding ligands from the Inforna^24^ and R-BIND databases,^25^ as visualized by the Uniform Manifold Approximation and Projection for Dimension Reduction (UMAP)^26^ analysis (**Figure S1A**; see Supplemental Dataset for structures). Each small molecule contains a diazirine cross-linking module that is photoactivated by UV light to form a covalent bond with bound RNA targets and an alkyne handle that can react with either azide-functionalized fluorescent dyes for imaging or azide-functionalized agarose beads for pull-down and enrichment of the bound RNAs (**Figure 2A**, left panel).

**Figure 2.**
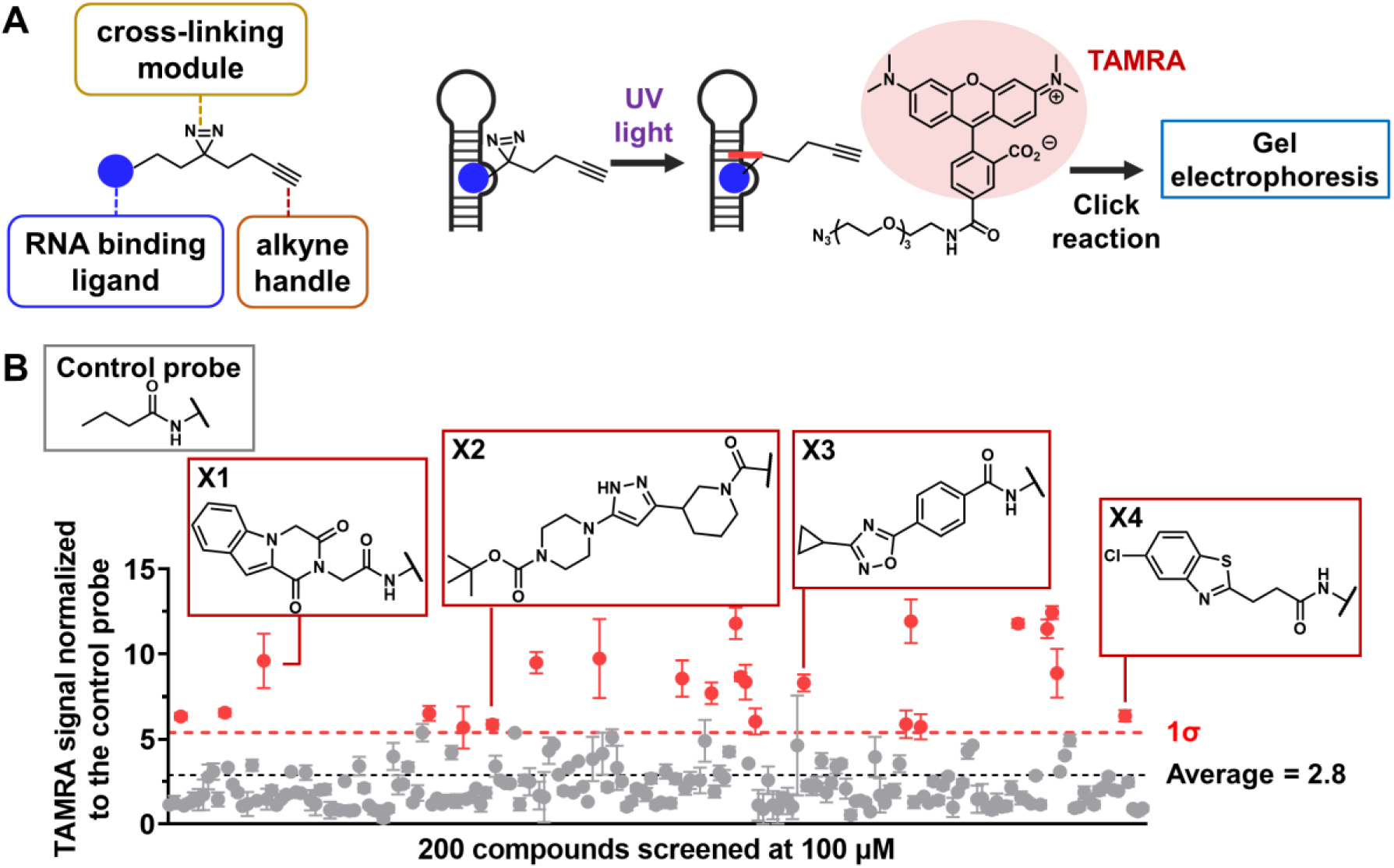
Transcriptome-wide mapping of small molecule-RNA interactions. (A) The general structure of functionalized small molecules containing a diazirine cross-linking module and an alkyne handle and schematic of *in vitro* screening to identify small molecules that preferentially interact with total RNA isolated from MDA-MB-231 cells. After UV irradiation to form covalent bonds between the small molecules and bound RNA, a fluorescent dye was conjugated to the alkyne handle by using click chemistry. The subsequent gel electrophoresis and fluorescence imaging enabled quantification of percent reaction between the small molecules and RNA. (B) Amongst 200 small molecules screened in this assay (*n* = 2 independent replicates), 23 were identified as hits with fluorescence signals that are 1σ above the average, after normalizing to the background signal from the control probe lacking the RNA-binding module. All data are reported as mean ± S.D.

To assess the general reactivity of these 200 small molecules, total RNA was isolated from MDA-MB-231 TNBC cells (10 µg RNA per sample) and incubated individually with each small molecule probe (100 µM) at room temperature, followed by UV light irradiation to induce formation of covalent bonds between the small molecule and bound RNA (**Figure 2A**, right panel). To visualize and quantify the reactivity of each compound, an azide-functionalized TAMRA (5-carboxytetramethylrhodamine) dye was conjugated to the alkyne handle of the RNA-small molecule complex via a click reaction. The TAMRA-labeled RNA-small molecule complexes were then analyzed by gel electrophoresis and fluorescence imaging (**Figure 2A**, right panel). A control probe lacking the RNA-binding module (“**Control probe**”) was included to measure background reactivity of the diazirine, and the TAMRA signals from each FFF-treated RNA sample were normalized to that of the control probe (**Figures 2B & S1B**). Fragments with signals 1σ (1σ = 2.5) higher than the mean (2.8 ± 2.4-fold as compared to **Control probe**) were deemed as hits, affording 23 small molecules that bind cellular RNAs (**Figures 2B & S2**). Comparison of the physicochemical properties of the 23 identified hits with the remaining 177 fragments (non-hits) in the library revealed that the hits contained a greater number of aromatic rings (1.8 ± 0.4 vs. 1.4 ± 0.6; *p* < 0.01) and aromatic nitrogen atoms (1.8 ± 1.0 vs. 1.2 ± 1.3; *p* < 0.05). (**Figure S1C**). This is perhaps not surprising as small molecule-RNA interactions are often governed by aromatic stacking and hydrogen bonding.^27^ Interestingly, there is no significant difference between the hit and non-hit small molecules in other properties including cLogP (partition coefficient), cLogS (aqueous solubility), the numbers of hydrogen bond donors or acceptors, total surface area (SA), or total polar surface area (TPSA) (**Figure S1C**).

### Feature Analysis of RNA-Binding Fragments Using a Random Forest Model

To assess whether RNA-binding fragments share common structural features, substructure analysis was performed using DataWarrior.^28^ No statistically significant scaffolds were identified, likely due to the limited number and structural diversity of binders (**Figure S3**). Consequently, a machine learning approach was employed to capture molecular features associated with RNA binding (**Figure 3**).

**Figure 3.**
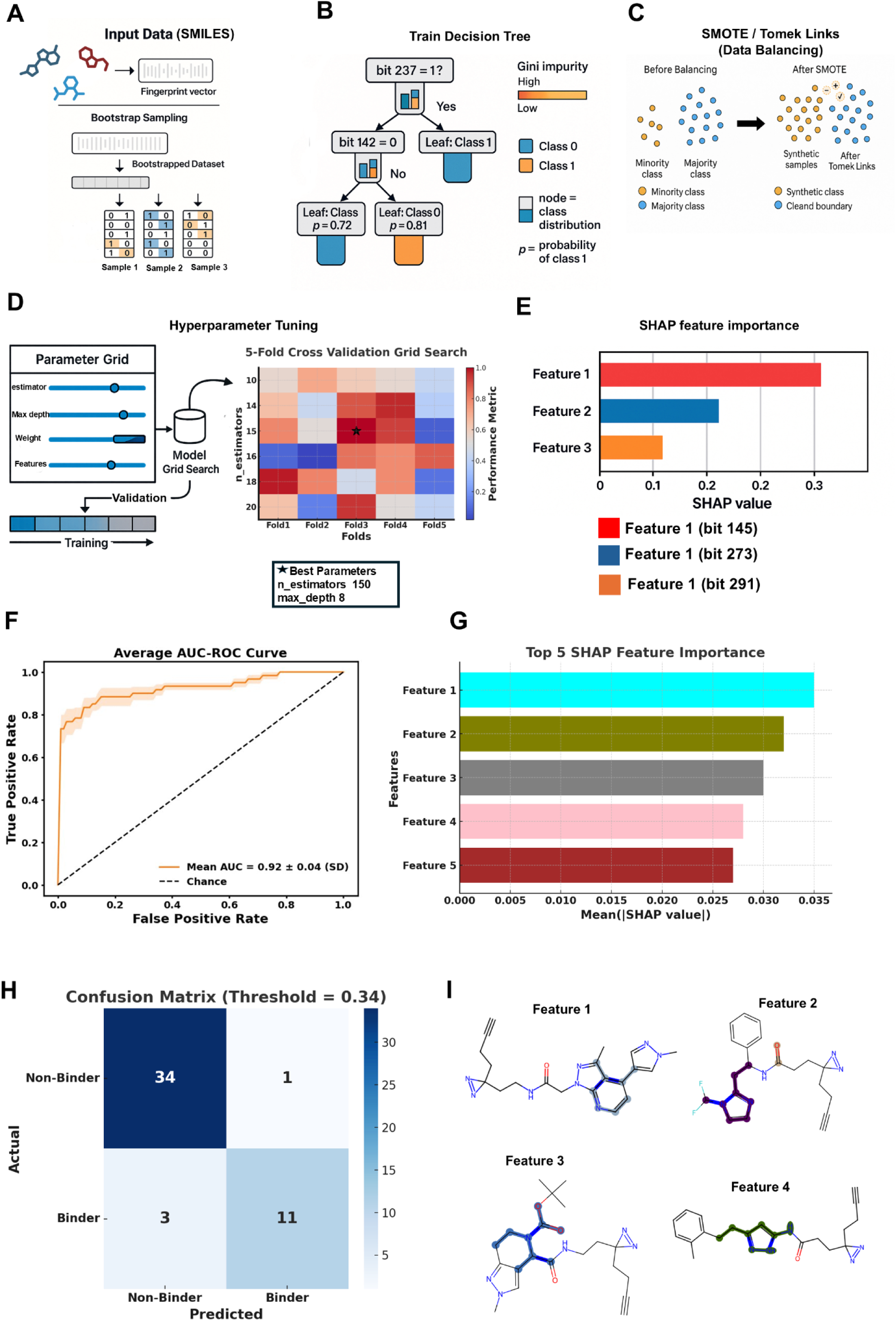
Workflow for building a binary Random Forest model from SMILES. (A) SMILES are converted to fingerprint vectors (Morgan/RDKit/AtomPair) and labeled as binder (Class 1) or non-binder (Class 0). Bootstrap sampling ensures trees in the forest see varied subsets of data. (B) Each decision tree splits on fingerprint bits to reduce impurity, with leaf nodes outputting class probabilities. Predictions are averaged across trees. (C) Class imbalance is addressed with SMOTE oversampling and Tomek-link cleaning applied within training folds. (D) Hyperparameters are optimized by stratified k-fold CV; the heat map summarizes mean AUROC/F1, with the star marking the best setting. (E) SHAP values identify influential fingerprint features, which can be mapped back to substructures. (F) ROC curve shows strong model performance (mean AUROC = 0.92 ± 0.04). (H) Confusion matrix (threshold = 0.34) shows 45/49 samples correctly classified with high specificity and balanced sensitivity. (I) Top SHAP features were traced to SMILES fragments, highlighting chemically intuitive motifs that drive binder prediction.

Molecular structures were represented using three complementary chemical fingerprints (Morgan^29^, RDKit^30^, and atom-pair^31^), which together encode both local and global aspects of chemical architecture (**Figure 3A**). These fingerprints were used as input to a random forest classifier trained to distinguish binders from non-binders (**Figure 3B**). Data imbalance between the two classes was corrected using synthetic oversampling (SMOTE)^32^ and Tomek Link^33^ (**Figure 3C**) filtering to improve classification accuracy (details provided in the Supporting Information).

The model was optimized through cross-validation and hyperparameter tuning (**Figure 3D, E**), yielding strong predictive performance (mean ROC-AUC = 0.92 ± 0.04) (**Figure 3F**). This strong separation was further supported by the confusion matrix, which showed that the majority of non-binders (34/35) and binders (11/14) were correctly classified (**Figure 3H**). This result indicated that the combined fingerprint approach was capable of reliably differentiating binders from non-binders based solely on molecular features. Interpretation of the model using SHAP (**Figure 3E, G**) (Shapley Additive exPlanations)^34, 35^ analysis revealed that aromatic and heteroatom-rich substructures contributed most strongly to RNA binding, suggesting that planarity, polarizability, and hydrogen-bonding potential are key determinants of recognition (**Figure 3A-I**).

Further analysis of statistically significant molecular descriptors^36^ showed that RNA-binding fragments exhibit distinct electrostatic surface distributions and three-dimensional shapes compared to non-binders (**Figure S4, Table S2**). Specifically, descriptors reflecting partial charge separation, dipole moment, and surface curvature were enriched among binders, consistent with preferential interactions with the negatively charged RNA backbone and structured grooves (details provided in the Supporting Information).

Together, these analyses indicate that RNA-binding fragments are not defined by a single scaffold but by a combination of physicochemical properties that favor electrostatic complementarity, stacking interactions, and defined molecular geometry. The machine learning framework thus establishes a predictive basis for identifying and optimizing small molecules with enhanced RNA affinity and selectivity. Detailed methodological information, hyperparameter settings, and descriptor statistics are provided in the Supporting Information.

### Analyzing ligandability patterns across the transcriptome

The binding landscape for each of the 23 fragments was profiled across the entire human transcriptome in live MDA-MB-231 cells by using Chem-CLIP-Map-Seq pipeline (**Figure 4A**).^16–18^ In brief, MDA-MB-231 cells were incubated with each small molecule or control diazirine probe (**Control probe**; 20 µM, 16 h), followed by irradiation with UV light, harvesting of total RNA, and pull-down of bound targets with azide-functionalized agarose beads. RNA-seq analysis was then used to compare: (i) the enrichment of each transcript in the pulled-down fraction, as compared to the input RNA, which was not subjected to the pull-down steps; and this enrichment as compared to the enrichment observed for the **Control probe**. In these analyses, transcripts, whether coding or noncoding (nc), were deemed a target of the small molecule or the **Control probe** if a minimum threshold of enrichment of 1.5-fold and a minimum read count of 5 were obtained, as previously implemented (p < 0.01; **Figure 4B**).^18^ All transcripts that met these minimum threshold criteria for **Control probe** were excluded from further analysis. This comprehensive and unbiased nature of this rich dataset enables patterns of the recognition of cellular RNAs to be elucidated, as described below.

**Figure 4.**
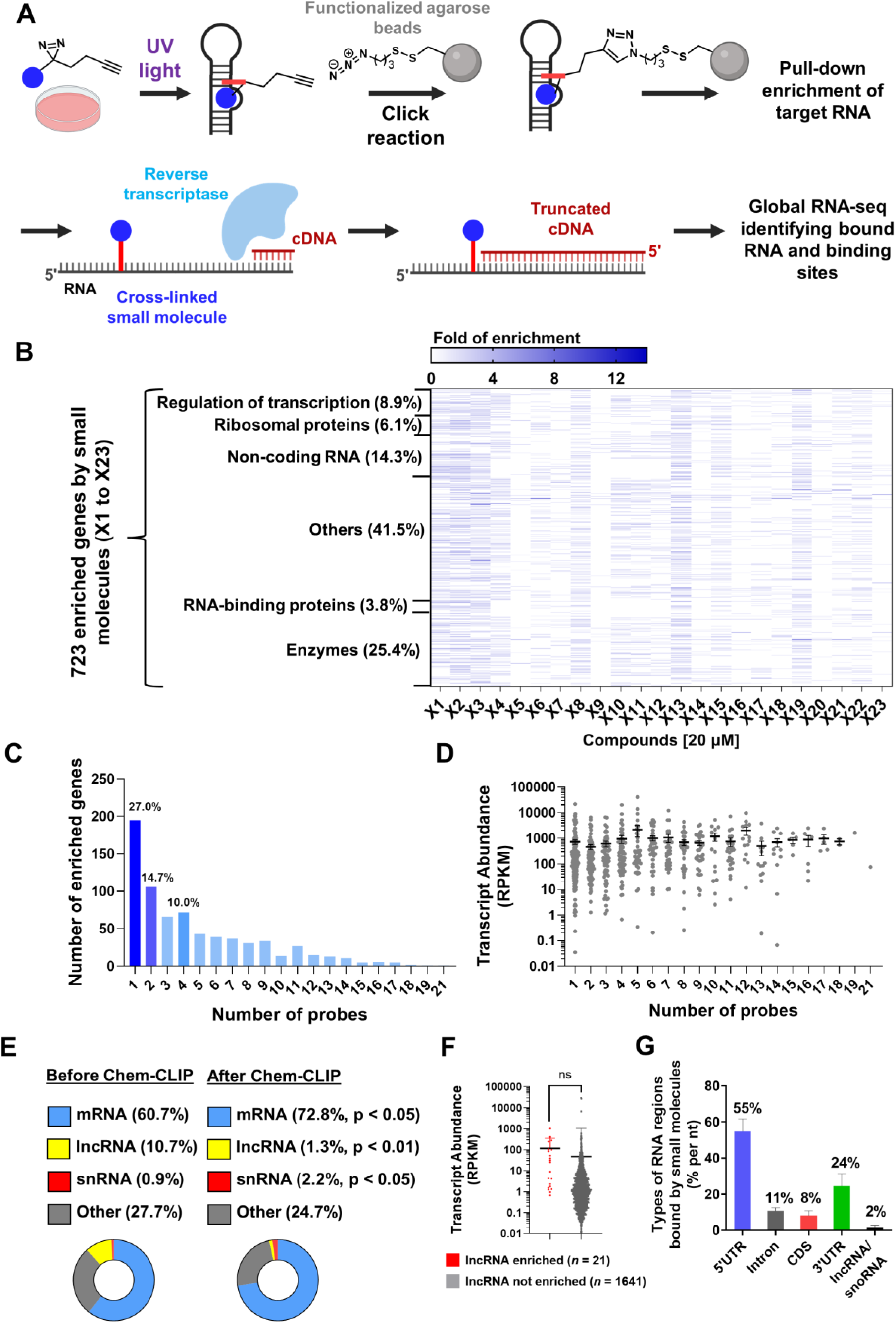
Ligandability analysis across the human transcriptome in live cells. (A) Schematic of mapping binding sites of small molecules on target RNA in live cells. The 23 hit small molecules were treated in live MDA-MB-231 cells (20 µM, 16 h) and their bound RNA targets were enriched by pull-down with azide-functionalized beads. The binding sites of small molecules were mapped by using global RNA-seq to identify the “RT-stop” sites during the reverse transcription. (B) Heat map summary of 723 RNA targets bound by the 23 hit small molecules (**X1** to **X23**) in MDA-MB-231 cells by using Chem-CLIP-Map-Seq. The types of proteins encoded by these RNAs are grouped as shown on the y-axis. (C) Distribution of the number of genes enriched by each profiled probe. (D) Transcript abundance for genes with different enrichment specificities, as indicated by the number of probes that simultaneously enrich the same target. (E) The enrichment (mRNA and snRNA) and depletion (lncRNA) of different RNA classes during the Chem-CLIP process. The abundance of each RNA class is quantified by using the number of reads mapped from global RNA-seq data. The p values are calculated by using the Chi squared tests. (F) Transcript abundances for lncRNAs that were or were not enriched by a fragment in these studies. No significant difference in transcript abundance between bound and unbound lncRNAs was observed. (G) Types of RNA regions bound by **X1** to **X23** for all 723 targets. % by nt indicates distribution of enriched regions after normalizing to the average length of each type calculated from all known human transcripts.All data are reported as mean ± S.D.

Amongst >20,000 genes detected by RNA-seq, a total of 723 unique RNA targets, including both coding and ncRNAs, were identified to be enriched by at least one of the 23 small molecule probes (p < 0.01; **Figure 4B & Supplemental file 1**). Notably, distinctive patterns of molecular fingerprints by each small molecule are observed, supporting that changes in chemical structures alter the molecular recognition between small molecules and RNA targets (**Figure 4B**). The number of RNA targets bound by each small molecule ranges from 20 – 326, while the fold enrichment ranged from 1.5 – 14.0 (**Figure 4B & Supplemental file 1**), suggesting differences in specificity and affinity (target occupancy), respectively. Particularly, **X9** and **X5** bound the fewest transcripts, 20 and 23, respectively, while **X3** and **X4** bound the most, 326 and 308, respectively. The highest fold enrichment was observed for **X3** and its engaged target Major Histocompatibility Complex, Class I, A (*HLA-A;* 14.0-fold). For the 723 unique RNA targets, 27.0% were enriched by only one probe, 14.7% were enriched by two probes, and the remaining were enriched by three or more probes (up to 21) (**Figure 4C**). One target, TMEM123, was bound by 21 probes, indicating the most promiscuous and non-specific binding events with small molecules (**Supplemental file 1**). No clear correlation between transcript abundance and the target enrichment specificity, as indicated by the number of probes that simultaneously enrich the same target, was observed (**Figure 4D**).

Next, the types of RNA enriched by each of the 23 small molecules were investigated. Messenger RNAs (mRNA) were significantly enriched by small molecules, where they comprised 60.7% of total reads in the input RNA as compared to 72.8% of total reads in the enriched fractions (p < 0.05 by Chi squared test; **Figure 4E**), suggesting that mRNA is more favorably bound by small molecules than other RNA classes. In contrast, long noncoding RNA (lncRNA) were generally not bound by the fragments, dropping from 10.7% of total reads in total RNA harvested from MDA-MB-231 cells to 1.4% of total reads of bound transcripts (p < 0.01 by Chi squared test; **Figure 4E**). To understand this observation for lncRNAs, we first looked at their expression levels in the RNA-seq data. Among 6546 known human lncRNAs, 1662 were detected; on average those that were enriched (n = 21; 1.3%) were expressed at higher levels than those that were not, although this difference is not statistically significant (115.4 ± 51.5 RPKM vs. 42.3 ± 24.9 RPKM; p = 0.24; **Figure 4F**). The reduced read counts in the enriched fractions were not due to fewer FFFs binding lncRNAs as 22 of 23 fragments enriched at least one lncRNA, where 15/21 lncRNAs (71%) were enriched by at least 2 probes. Collectively, these data suggest that lncRNAs in general may not readily interact with small molecules, they may form dynamic structures that are therefore not ligandable, or their interactions have rapid kinetics.

To investigate the regions, whether the 5’ untranslated region (UTR), coding sequence (CDS), or 3’ UTR, within mRNAs that are preferentially bound by small molecules, their binding sites were mapped by identifying truncations of the cDNA; that is, the covalent cross-link formed between an RNA target and the fragment impedes reverse transcriptase and causes an “RT stop”, or truncation in the cDNA.^17, 18, 21^ The number of identified presumptive “RT-stop” sites (peak start for positive-strand RNA and peak end for negative-strand RNA) for each fragment ranged from 20 to 326, consistent with the number of RNA targets bound by each small molecule. Notably, the lengths of the 5’ UTRs, CDS, and 3’ UTRs differ and to control for this intrinsic bias, the average length of each element calculated from the NCBI database^37^ was applied as a normalizing factor, affording the percentage of mapped binding events per nucleotide (**Figures 4G & S5**). Interestingly, the vast majority of binding sites (79%) were mapped to the UTRs, with 55% and 24% mapped to the 5’ UTR and 3’ UTR respectively, which aligns with previous studies demonstrating higher levels of structured RNA folds within UTRs than introns or coding regions (generally thought to be dynamic or unstructured).^2, 3^ The presence of many regulatory elements located in UTRs of mRNA,^38^ for example iron responsive elements ^39–41^ and internal ribosomal entry sites,^42^ is well documented, and bioactive small molecules that target these functional elements and modulate biology have been reported.^43–45^ These results suggest that structures present in the UTRs of mRNAs should be prioritized for small molecule discovery efforts.

As aforementioned, some transcripts are bound by more than one fragment. For example, *HSPD1* and *hnRNP H1* were bound by 17 fragments while *TRFC* and *hnRNP U* were bound by 15 fragments. Thus, we determined whether fragment binding sites clustered in loci in the 25 transcripts that bound the greatest number of fragments (range: 6 – 17 fragments; **Figure S6**). The transcripts were binned into two groups: (i) those that tightly clustered in a particular locus as defined by two or more fragments that have partially or fully overlapped binding regions; and ^46^ those that are moderately clustered where at least one fragment had a unique binding region (**Figure S6A**). For *HSPD1*, 16 of 17 binding fragments cluster in a region comprising nt 197,486,820-197,487,010 (**Figure S6B**). These data suggest that these transcripts form robust, ligandable structure and that this methodology could be used to map the RNA structurome. In contrast, the least clustered transcript among the top 25 transcripts showed only moderate clustering (i.e. TFRC, 9 out of the 15 bound fragments showing clustered binding regions). Perhaps unsurprisingly, transcripts that bound fewer fragments were more tightly clustered than those bound more (p < 0.05; **Figure S6C**). No significant differences in fold enrichment or transcript abundance for tightly vs. moderately clustered transcripts were observed (**Figure S6D-E**).

Our previous transcriptome-wide profiling studies suggested that small molecules preferentially bind regions of RNA that form thermodynamically stable structures.^19^ To gain insight into the generalizability of this observation as well as the secondary structure of the small molecule binding site, a thermodynamic analysis of each binding site identified by our cellular Chem-CLIP-Map-Seq pipeline was completed. A computational pipeline, ScanFold 2.0,^47, 48^ was used to model the structure of the binding sites. ScanFold identifies unusually thermodynamically stable structures by comparing the minimum free energy (MFE) of the native RNA sequence to the minimum free energy of a library of randomized sequences. The MFEs for functional RNA structures are usually lower than a random sequence,^49–51^ and this can be attributed to the evolution and composition requirements for functional RNAs.^51^ The difference in stability between the native and random sequences is quantified by the statistical parameter, z-score. A negative z-score indicates that the native sequence is more thermodynamically stable than the average MFE of all shuffled sequences, suggesting that the native sequence may be conserved and functional. Notably, amongst the 723 RNA targets bound by small molecules in this study, 415 (63%) have negative z-scores in or nearby their respective small molecule binding sites (average z-score of –2.99 ± 0.02; average ΔG°_37_ = −24.67 ± 0.27 kcal/mol); that is, they are unusually thermodynamically stable, suggesting that these regions may be evolutionarily conserved with functional structures (**Figure S7**).

### Converting RNA binders into RiboTACs targeting cancer-dependent mRNAs

Our previous studies have shown that conversion of an RNA binder into a RiboTAC improves selectivity, a composite of the RNA binder’s selectivity, the inherent specificity of RNase L for unpaired pyrimidines, the juxtaposition of the binding site and an RNase L-sensitive site, and how closely the linker mirrors this distance.^19, 22, 52–56^ Similar phenomena have been observed for protein-targeting small molecules where the selectivity of promiscuous protein binders has been improved by conversion to a proteolysis-targeting chimera (PROTAC) and where the selectivity of PROTACs was enhanced by optimizing linker length and attachment point.^57, 58^ We therefore converted the four most promiscuous FFFs, **X1 – X4** into their respective RiboTAC by appending each with a small molecule that locally activates endogenous RNase L in cells thus inducing cleavage of the bound RNA targets via induced proximity (**Figures 5A & S8**).

**Figure 5.**
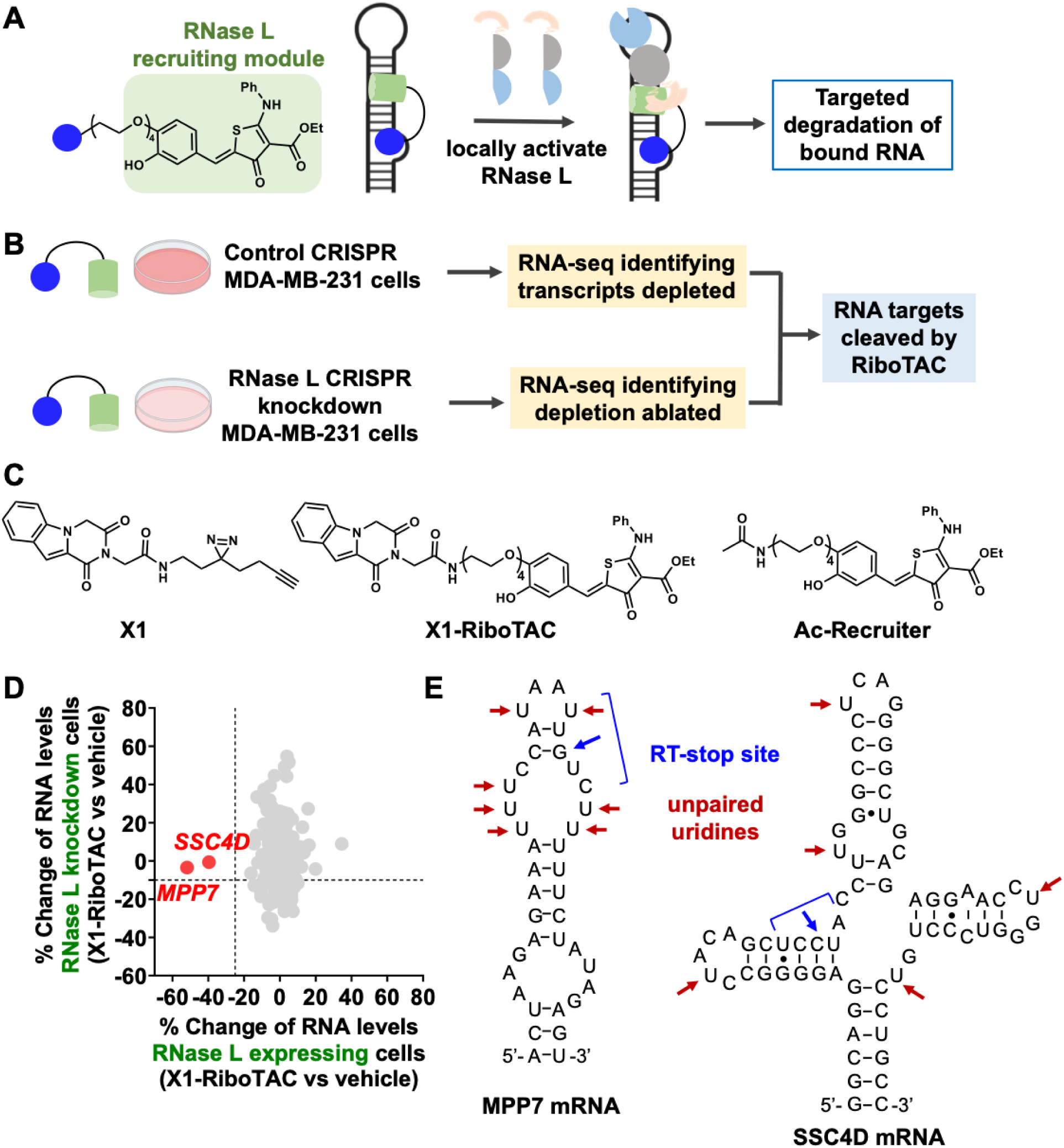
Conversion of RNA-binding small molecules in RiboTAC degraders. (A) General structure of ribonuclease targeting chimeras (RiboTACs) and a schematic of RiboTAC locally activating RNase L to cleave the RNA targets in cells. (B) Two CRISPR modified MDA-MB-231 cell lines were used to assess RiboTACs, one with a control gRNA (express RNase L) and the other with RNase L knockdown by ∼60%.^77^ Treatment of RiboTACs in these two cell lines followed by global RNA-seq to identify the *bona fide* targets cleaved by RiboTACs in an RNase L dependent manner. (C) Structures of **X1**, **X1-RiboTAC**, and a control compound lacking the RNA-binding module (**Ac-Recruiter**). (D) Transcriptome-wide profiling identified two targets (*MPP7, SSC4D*) that were cleaved (>25% reduction) by **X1-RiboTAC** in the control cell line expressing RNase L, with no effect (<10% change) in the RNase L knockdown cell line (*n* = 3 biological replicates). (E) Predicted RNA structures of *MPP7* and *SSC4D* near the **X1** binding sites. The blue bracket indicates the range of mapped RT-stop sites induced by cross-linking of **X1**, and the blue arrow indicates the RT-stop site with the highest statistical confidence. The red arrows indicate unpaired uridines, which are preferred cleavage sites by RNase L.

To identify the targets cleaved by the four RiboTACs in cells, two CRISPR-modified MDA-MB-231 cell lines were employed: one in which RNase L expression was knocked down and the other as a control cell line that expresses RNase L to a similar level as wild type (WT) MDA-MB-231 cells.^19, 55^ Our previous studies showed that these two CRISPR-modified cells lines and WT MDA-MB-231 have very similar transcriptome profiles and thus can be compared to each other.^19^ The two CRISPR cell lines were each treated with a RiboTAC of interest in parallel (**X1-** to **X4-RiboTAC**; 10 µM, 48 h), followed by global RNA-seq analysis to assess their effects on the entire transcriptome (**Figure 5B**). A *bona fide* RNA target cleaved by a RiboTAC has reduced abundance in the control CRISPR cell line expressing RNase L, but with no change in abundance in the CRISPR RNase L knockdown cell line. That is, an RNA target affected in both cell lines upon RiboTAC treatment (RNase L-independent) is likely due to perturbation of a downstream biological pathway rather than RiboTAC-induced cleavage.

Interestingly, amongst a total of 473 RNA targets bound by **X1** – **X4**, only two targets were cleaved in an RNase L dependent manner (>25% decrease in control CRISPR cells and <10% change in RNase L knockdown cells), *MPP7* and *SSC4D*, both by **X1-RiboTAC** (**Figures 5C-D & S9**). **X1** bound 282 transcripts, of which the binding sites within 102 transcripts could be identified from Chem-CLIP-Map-Seq data (**Figure S6**). That is, an RT stop site marked by a drop in RNA-seq reads^59^. *MPP7* is a scaffold protein that promotes the migration of breast cancer cells^60, 61^ while *SSC4D* has no known associated oncogenicity. These results underscore the selectivity of RiboTACs across the transcriptome, as only a small fraction (2/473; 0.42%) of bound RNA targets was cleaved by RiboTACs.

The MPP7 transcript levels were reduced by 52 ± 3% (p = 0.05) while *SSC4D* mRNA levels were reduced by 40 ± 2% (p = 0.02). *MPP7* and *SSC4D* are less abundant than the other **X1**-bound transcripts (on average 91.3 ± 7.8 RPKM), while their transcript abundances are similar to each other (0.35 RPKM for *MPP7*, 0.10 RPKM for *SSC4D*). Both *MPP7* and *SSC4D* showed comparable enrichment by **X1** (2.45-fold for *MPP7*, 2.05-fold for SSC4D) to each other as well as the average enrichment of other **X1**-bound transcripts (2.50-fold) (**Supplemental file 1**). The **X1** binding site for both transcripts was mapped to the 3’ UTR (the only **X1**-binding site for both transcripts in RNA-seq data; *MPP7* and *SSC4D* were not bound by the other 22 probes). Meanwhile, **X1** preferentially bound 5’ UTR regions for other engaged transcripts (53.9%) compared to regions including 3’ UTR (26.4%), intron (10.5%), CDS (8.2%), or ncRNAs (1.1%) (**Figure S5**).

The structure formed by the *SSC4D* binding site is unusually thermodynamically stable, with a z-score of −3.08 (ΔG°_37_ = −31.3kcal/mol), comprising a three-way junction, in which two of the hairpins have an internal loop embedded in the stem. The cross-linked nt is harbored at the end of one of the helices nearby the junction (nt C22; **Figure 5E**). Although there is an unusually thermodynamically stable site nearby the *MPP7* binding site (z-score = −3.37; ΔG°_37_ = −21.2 kcal/mol), the binding site itself did not appear as a structured region. Folding a 120 nt window by free energy minimization (RNAstructure^62^) where the cross-linked site is centered in the window produces a thermodynamically stable structure with ΔG°_37_ = −23.6 kcal/mol. The region containing the cross-linked nt G23 adopts a 4-nt hairpin with 4×4 nt and 4×3 nt internal loops embedded in the stem (**Figure 5E**), where G23 forms a closing GC base pair of the 4×4 nt internal loop adjacent to the hairpin (**Figure 5E**). Interestingly, both the *MPP7* and *SCC4D* binding sites are nearby pyrimidine-rich, non-canonically paired structures, which are preferred cleavage sites for RNase L; nearby (within 15 nt of sequence space) the **X1**-binding site in *MPP7* and *SCC4D* are eight and three unpaired uridines, respectively, although two additional unpaired uridines are close in 3-dimensional space for *SSC4D*, *vide infra* (**Figure 5E**).

To verify the activity of **X1-RiboTAC** observed from RNA-seq analysis, reduction of *MPP7* and *SSC4D* transcripts was measured in WT MD-MB-231 cells alongside two control compounds, **X1** (lacking the RNase L-recruiting module) and **Ac-Recruiter** (lacking the RNA binding module) (**Figure 6A**). After 48 h treatment, neither **X1** nor **Ac-Recruiter** affected the abundance of *MPP7* and *SSC4D* mRNAs (**Figure 6A**, left panel). In contrast, **X1-RiboTAC** dose-dependently reduced both *MPP7* and *SSC4D* mRNA levels. In line with RNA-seq analysis (**Figure 6A**), treatment with 10 µM of the RiboTAC reduced *MPP7* transcript abundance by 61 ± 3% and *SSC4D* abundance by 47± 2% (p < 0.001 for both transcripts; **Figure 6A**). Similar results were observed in control CRISPR MDA-MB-231 cells expressing RNase L where **X1-RiboTAC** dose-dependently reduced both transcripts, *MPP7* by 61 ± 3% (p < 0.001) and *SSC4D* by 44 ± 5% at the 10 µM dose (p < 0.001) while **X1** and **Ac-Recruiter** had no effect on the abundance of either target (**Figure S10A**; see **Figures S10B-C** for primer validation).

**Figure 6.**
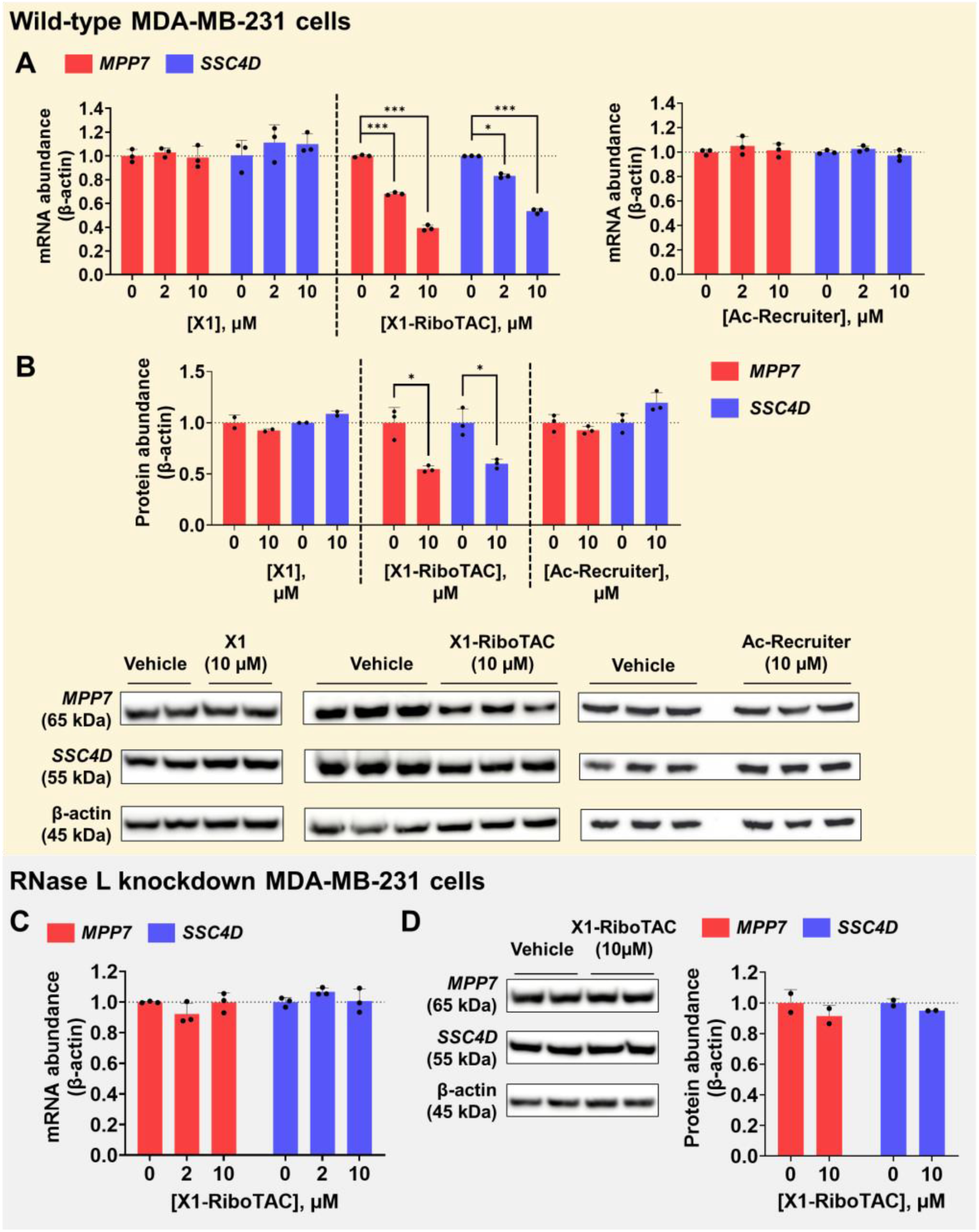
X1-RiboTAC reduces *MPP7* and *SSC4D* mRNA and protein levels in an RNase L dependent manner. (A) Effect of **X1**, **X1-RiboTAC**, and **Ac-Recruiter** on the abundance of *MPP7* and *SSC4D* in WT MDA-MB-231 cells, as measured by RT-qPCR (*n* = 3 biological replicates). (B) Effect of **X1**, **X1-RiboTAC**, and **Ac-Recruiter** on the *MPP7* and SSC4D protein abundance in WT MDA-MB-231 cells, as measured by Western blot (*n* = 2 biological replicates for **X1** and 3 biological replicates for **X1-RiboTAC**). (C) Effect of **X1**, **X1-RiboTAC**, and **Ac-Recruiter** on *MPP7* and *SSC4D* transcript abundance in CRISPR RNase L knockdown MDA-MB-231 cells, as measured by RT-qPCR (*n* = 3 biological replicates). (D) Effect of **X1-RiboTAC** on MPP7 and SSC4D protein levels in CRISPR RNase L knockdown MDA-MB-231 cells, as measured by Western blot (*n* = 2 biological replicates). * p < 0.05; *** p < 0.001, as determined by two-tailed Student’s t-test. All data are reported as the mean ± SD.

Assessment of the effect of **X1-RiboTAC**, **X1**, and **Ac-Recruiter** on MPP7 and SSC4D protein levels in WT MDA-MB-231 cells showed that **X1** (10 µM, 48 h treatment) had no effect on the abundance of either protein, as determined by Western blotting; that is, its binding to these two mRNA transcripts is biologically silent (**Figure 6B**). Similarly, treatment of **Ac-Recruiter** at 10 µM did not decrease MPP7 or SSC4D protein levels upon 48 h incubation, indicating RNase L binding event alone was not sufficient to drive decay of the two targets. In contrast, **X1-RiboTAC** (10 µM, 48 h treatment) reduced *MPP7* (45 ± 3% reduction, p < 0.05) and SSC4D (40 ± 4% reduction, p < 0.05) protein abundance (**Figure 6B**). Thus, the RiboTAC approach can convert inactive RNA binders into bioactive degraders that affect the function of their bound RNA targets.^22^

To further support the RNase L-dependency of **X1-RiboTAC** as indicated by RNA-seq studies (**Figure 5D**), the level of *MPP7* or *SSC4D* transcripts was measured in the CRISPR RNase L knock-down MDA-MB-231 cells upon treatment with **X1-RiboTAC** (2 μM and 10 μM; 48 h treatment). **X1-RiboTAC** had no effect at either dose on the abundance of *MPP7* or *SSC4D* transcript levels, supporting recruitment of RNase L as a mode of action to cleave the target RNAs (**Figure 6C**). As expected, MPP7 and SSC4D protein levels were not affected by **X1-RiboTAC** in this cell line (**Figure 6D**).

### Optimizing the structure of the X1 RNA-binding module to enhance RiboTAC selectivity

In previous studies, it was shown that various factors contribute to RiboTAC selectivity, as described above.^19, 22, 52–56^ Here, it is investigated whether the specificity of **X1-RiboTAC** could be improved for one transcript over the other by studying analogs of the **X1** RNA-binding module, where some analogs grow the fragment. A panel of ten **X1** derivatives (**X1-D1** to **X1-D10**) was selected with chemically diverse moieties appended to the core structure of **X1** (**Figure 7A**). To assess *MPP7* and *SSC4D* mRNA target engagement of these derivatives, a competitive Chem-CLIP (C-ChemCLIP) assay^17, 63^ was employed (**Figure 7B**). In this assay, WT MDA-MB-231 cells were co-treated with equal concentrations (20 µM, 16 h treatment) of the fully functionalized **X1** probe and one of its unreactive derivatives that lacks the diazirine and the alkyne groups. If **X1** and the analog bind to the same binding site in the RNA, the extent of cross-linking of **X1** and hence pull-down of its RNA target will be reduced, here as measured by RT-qPCR.

**Figure 7.**
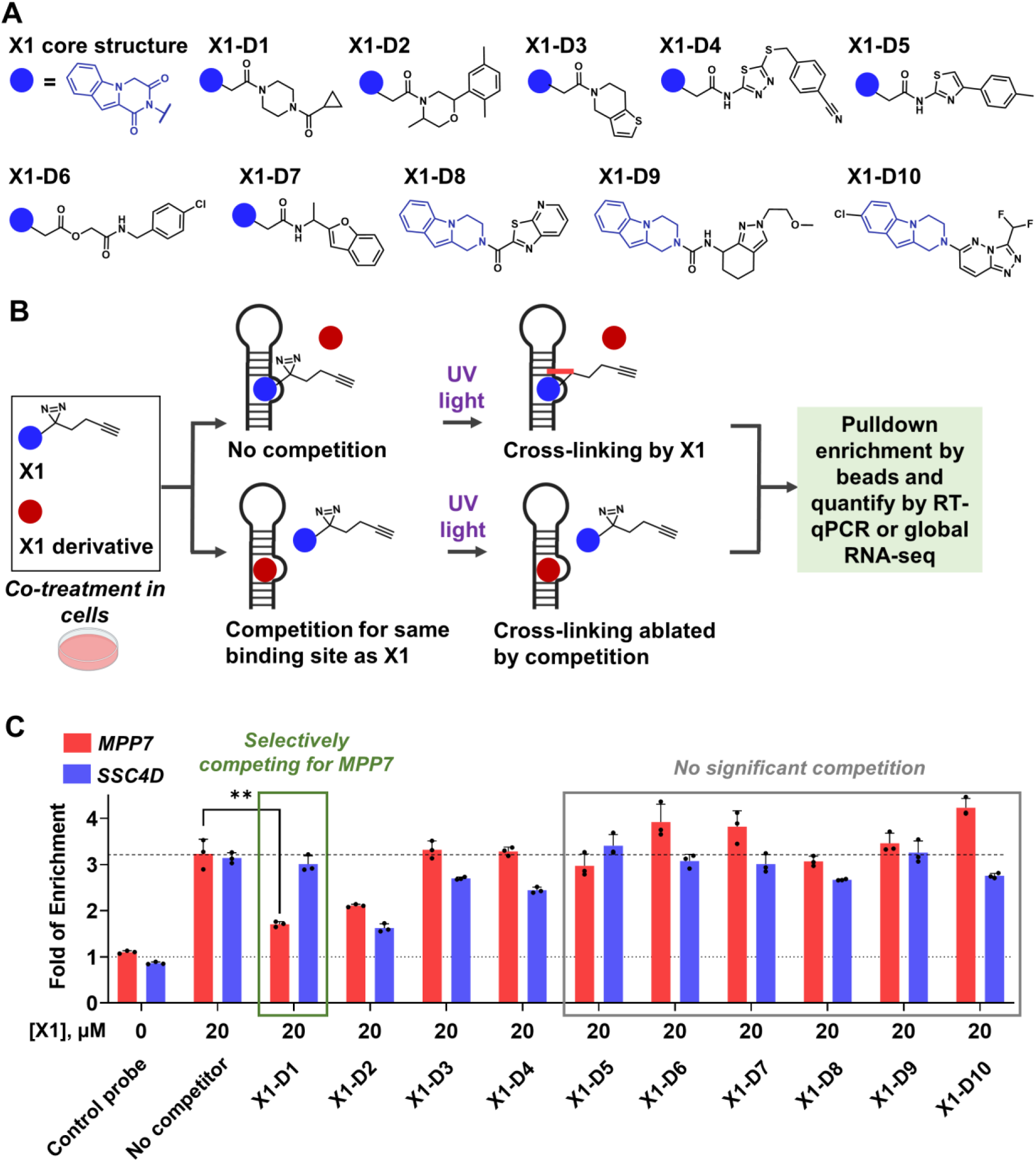
Competitive Chem-CLIP to identify X1 derivatives that selectively bind to *MPP7* over *SSC4D*. (A) Structures of the **X1** derivatives used in this study. (B) Schematic of competitive Chem-CLIP to screen **X1** derivatives. The **X1** probe (20 µM) containing the cross-linking (diazirine) and the pull-down (alkyne) modules was co-treated with **X1** derivates (20 µM) in wild-type MDA-MB-231 cells to assess if the derivatives compete off the enrichment of RNA targets from **X1**. (C) Competitive Chem-CLIP by co-treating **X1** with an equimolar (20 µM) of **X1** derivatives in wild-type MDA-MB-231 cells followed by pull-down enrichment as measured by RT-qPCR (*n* = 3 biological replicates). ** p < 0.01, as determined by two-tailed Student’s t-test. All data are reported as the mean ± SD.

In line with RNA-seq analysis (**Figure S11**) in the absence of competing small molecule, **X1** enriched *MPP7* or *SSC4D* mRNA by 3.23 ± 0.32-fold and 3.14 ± 0.12-fold, respectively (**Figure 7C**). In competition experiments, six of the ten **X1** derivatives (**X1-D5** to **X1-D10**) showed no significant competition with **X1** for engaging either *MPP7* or *SSC4D* mRNA (<25% reduction in the fold enrichment after pull-down) (**Figure 7C)**. These observations suggest that the associated changes in the **X1** structure reduced or ablated affinity for the target, reduced cellular permeability, and/or changed cellular localization of the small molecule. Interestingly, both **X1-D1** and **X1-D2** competed with **X1** for binding. While **X1-D2** competed the enrichment of both targets to similar extent (34 ± 9% reduction in enrichment for *MPP7* and 28 ± 3% reduction in enrichment for *SSC4D*; p < 0.01), **X1-D1** selectively competed the enrichment of *MPP7* (47 ± 2% reduction in enrichment; p < 0.01) with no effect on the enrichment of *SSC4D* (<10% competition, p < 0.05) (**Figure 7C**). These results indicate that the 1-(4-(cyclopropanecarbonyl)piperazin-1-yl)ethan-1-one moiety of **X1-D1** drives target selectivity towards *MPP7* mRNA (**Figure 8A**) and supports that in-cell C-Chem-CLIP can be used to study the relative occupancy of a series of compounds to an RNA target and facilitate identification of higher affinity binders.

**Figure 8.**
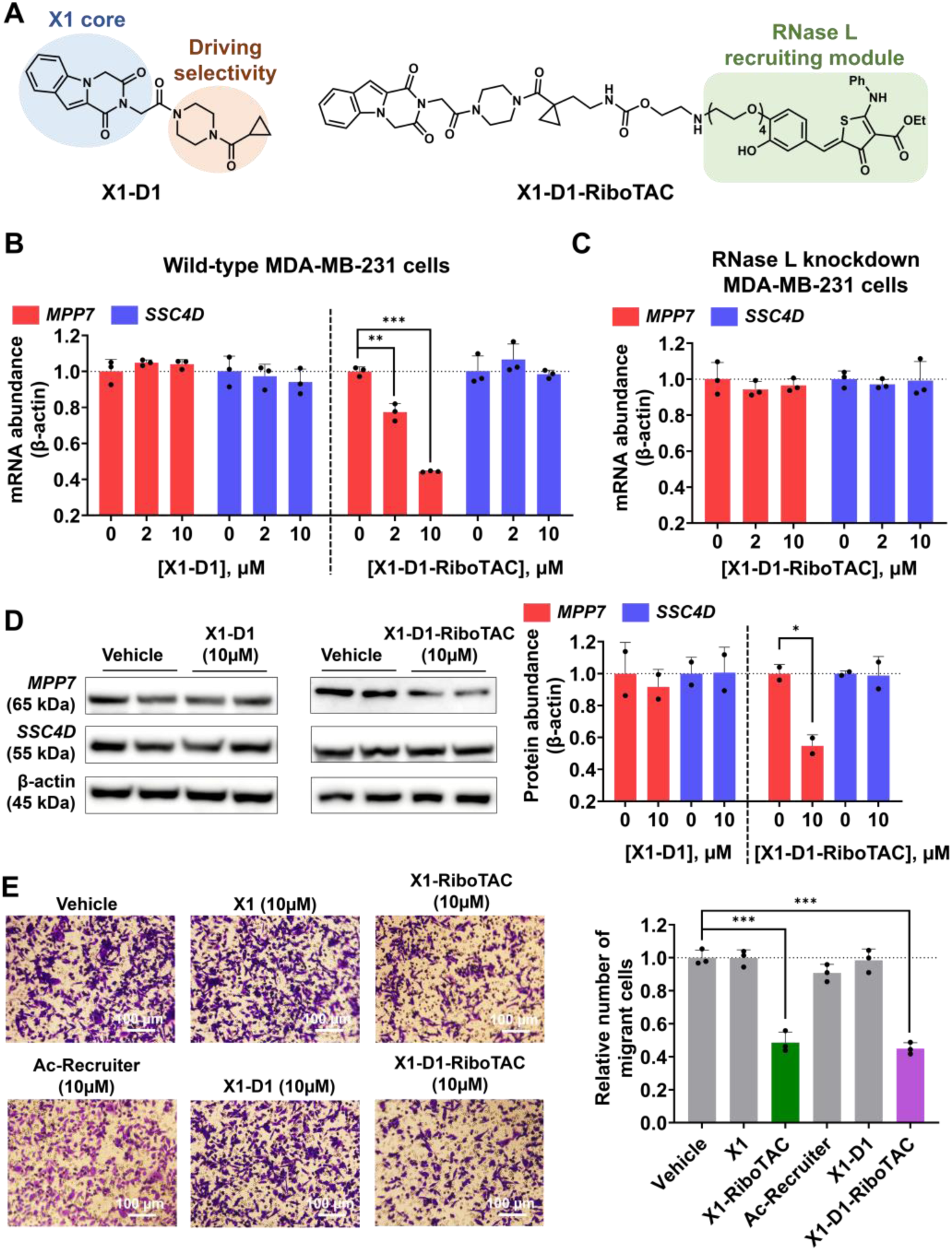
X1-D1-RiboTAC selectively cleaves *MPP7* mRNA and alleviates its associated oncogenic migrant phenotype. (A) Structures of **X1-D1** and **X1-D1-RiboTAC**. (B) Effect of **X1-D1** and **X1-D1-RiboTAC** on the mRNA levels of *MPP7* and *SSC4D* in wild-type MDA-MB-231 cells, as measured by RT-qPCR (*n* = 3 biological replicates). (C) Effect of **X1-D1-RiboTAC** on *MPP7* and *SSC4D* transcript abundance in RNase L knockdown MDA-MB-231 cells, as measured by RT-qPCR (*n* = 3 biological replicates). (D) Effect of **X1-D1** and **X1-D1-RiboTAC** on MPP7 and SSC4D protein levels in WT MDA-MB-231 cells, as measured by Western blot (*n* = 2 biological replicates). (E) Effect of **X1**, **X1-RiboTAC**, **Ac-Recruiter**, **X1-D1**, and **X1-D1-RiboTAC** on the number of migratory WT MDA-MB-231 cells, as measured by a Boyden chamber assay (*n* = 3 biological replicates). * p < 0.05; ** p < 0.01; *** p < 0.001, as determined by two-tailed Student’s t-test. All data are reported as the mean ± SD.

The selectivity of **X1-D1** was profiled transcriptome-wide by using the same C-Chem-CLIP workflow, followed by global RNA-seq to quantify its competition for all targets bound by the parent molecule **X1**. In agreement with the RT-qPCR results, at equimolar concentration (20 µM), **X1-D1** reduced the enrichment for *MPP7* by ∼ 36% with no effect on the other target *SSC4D* (**Figure S11A**). Amongst a total of 282 RNA targets bound by **X1**, only 18 targets (6.4%) were competed by **X1-D1** (>25% reduction in their fold enrichment), supporting the improved selectivity of **X1-D1** compared to the parent molecule **X1**. Notably, at a dose of 5 µM, **X1-D1** could not compete with the enrichment of *MPP7* by **X1** (20 µM). Increasing the concentration of **X1-D1** to 40 µM reduced enrichment of *MPP7* by ∼50%, while still no effect was observed on the enrichment for *SSC4D* (**Figure S11A**). To further validate *MPP7* target engagement of **X1-D1**, the latter was converted into a Chem-CLIP probe by appending it with the diazirine and alkyne groups (**Figure S11B**). In WT MDA-MB-231 cells, **X1-D1-Chem-CLIP** dose-dependently enriched *MPP7* mRNA, with an enrichment of 2.9 ± 0.2- and 3.9 ± 0.1-fold at the 5 µM (p < 0.01) and 20 µM (p < 0.001) doses, respectively, as compared to the control probe lacking the RNA-binding module (16 h treatment; **Figure S11B**). No enrichment of *SSC4D* mRNA was observed (2 – 20 µM; 16 h treatment; **Figure S11B**).

To assess if the selectivity of the RNA-binding module translates to an enhancement of RiboTAC selectivity, the RNase L-recruiting module was appended to **X1-D1**, affording heterobifunctional **X1-D1-RiboTAC** (**Figure 8A**). As observed for **X1**, **X1-D1** had no effect on the abundance of *MPP7* and *SSC4D* in WT MDA-MB-231 cells (**Figure 8B**). In contrast, **X1-D1-RiboTAC** dose-dependently reduced the abundance of *MPP7* mRNA (56 ± 4% at 10 µM, 48 h treatment, p < 0.001), with no effect observed on the abundance of *SSC4D* mRNA (**Figure 8B**). Compared to the parent **X1-RiboTAC** that cleaves both targets to similar extents (**Figure 6A**), these results suggest that improving the selectivity of the RNA-binding module can enhance the selectivity of target cleavage by its corresponding RiboTAC. The RNase L-dependency of **X1-D1-RiboTAC** was verified by its lack of activity in CRISPR RNase L knockdown MDA-MB-231 cells (**Figure 8C**) and reduction of *MPP7* mRNA in control CRISPR MDA-MB-231 cells where RNase L is expressed (65 ± 2% reduction at 10 µM; p < 0.001).

Similarly to **X1,** the binding of **X1-D1** is biologically silent, as no changes in MPP7 or SSC4D protein levels were observed in WT MDA-MB-231 cells (10 µM, 48 h treatment, **Figure 8D**) while **X1-D1-RiboTAC** reduced MPP7 protein levels by 46 ± 7% at a dose of 10 µM (p < 0.05), with no effect on the *SSC4D* protein levels (**Figure 8D**).

As *MPP7* is oncogenic, promoting a migratory phenotype of MDA-MB-231 TNBC cells,^60, 61^ the effects of **X1**, **X1-D1**, and their corresponding RiboTACs on this cellular phenotype were assessed by using a Boyden chamber assay. As expected, the two binders, **X1** and **X1-D1**, had no effect on the migration of WT MDA-MB-231 cells (10 µM, 48 h treatment; **Figure 8E**), as neither compound affected *MPP7* transcript or protein levels (**Figure 8D**). Likewise, control compound **Ac-Recruiter**, which lacks the RNA-binding module, was inactive (**Figure 8E**). In contrast, both **X1-RiboTAC** and **X1-D1-RiboTAC** reduced the number of migratory MDA-MB-231 cells, by 51 ± 6% and 55 ± 4% (p < 0.01), respectively (**Figure 8E**). [An antisense oligonucleotide (ASO) targeting the *MPP7* mRNA was used to support the on-target effect of reducing *MPP7* protein levels to inhibit the migration phenotype of MDA-MB-231 cells (**Figure S12**).]

Collectively, these studies showcase how the conversion of a promiscuous inactive RNA binder into a RiboTAC drives both bioactivity and target selectivity, which can be further reprogrammed by optimization of the RNA-binding module to precisely tune the effect towards the desired oncogenic target in a predictable manner.

### 3D Structure Modeling of X1 Bound to Its Transcript Targets by Molecular Docking

To gain insight into how the small molecule **X1** recognizes its RNA targets across the transcriptome, a multi-step computational approach integrating RNA secondary structure prediction, three-dimensional modeling, and molecular docking was employed (**Figure 9A**). Three-dimensional RNA models were generated for 106 transcripts (from 58 genes) identified by Chem-CLIP-Map-Seq using the FARFAR2 algorithm within the Rosetta suite.^64^ FARFAR2 assembles realistic RNA structures from known structural fragments and refines them to low-energy conformations. For each transcript, ten of the most energetically favorable models were selected and used for ensemble docking with AutoDock-GPU^65^ to identify potential **X1** binding sites (**Figure 9B**). Complete modeling details are provided in the Supporting Information.

**Figure 9.**
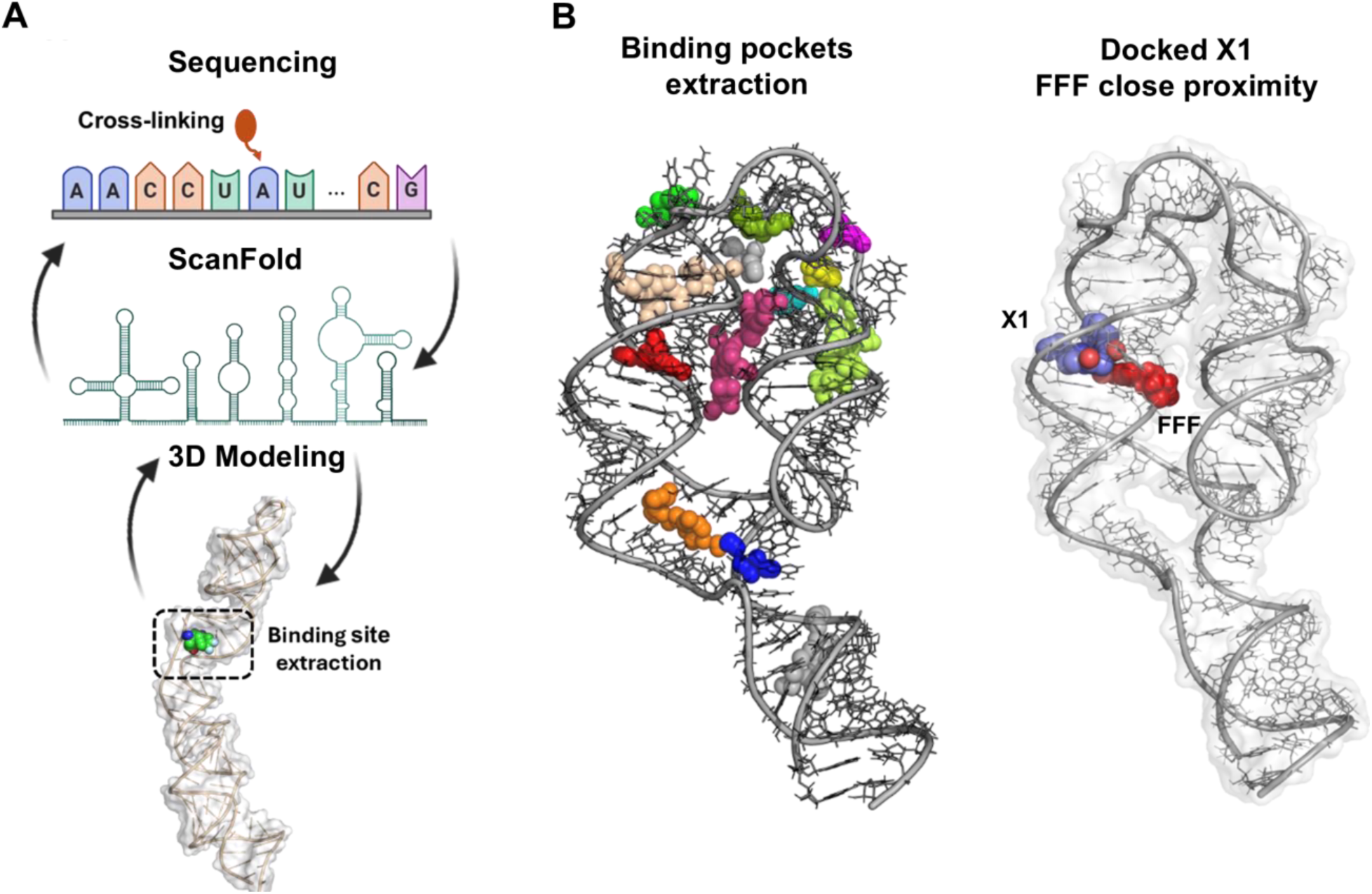
3D modeling from sequencing, ligandable pocket identification and blind docking. (A) Schematic representation of the workflow used for binding site extraction. The RNA sequence of the PTP4A2 gene is shown with the location of the FFF site (**X1**) highlighted. Structural modeling of the RNA allowed for the extraction of a defined binding region surrounding the FFF site (boxed). (B) Three-dimensional visualization of binding pockets within the RNA tertiary structure of the PTP4A2 transcript. Left: identified binding pockets are highlighted in color on the RNA structure, illustrating distinct cavities distributed along the RNA fold. Right: **X1** was docked into the RNA structure (shown in sphere), revealing ligands positioned in close proximity to the FFF site (shown in red spheres). RNA is shown in sticks, cartoons and surface.

To evaluate the accuracy of docking predictions, the positions of docked **X1** molecules were compared to experimentally mapped cross-linked nucleotides from Chem-CLIP data. Docking poses were considered correct when the ligand center was within 10 Å of the experimental cross-link site. Using this criterion, approximately 69% of poses were consistent with experimental data, indicating strong agreement between modeling and experiment (**Figure S13A**). Further analysis of the first three lowest energy docked poses per each gene targeted by **X1** shows for all of the genes except for AK6, HMGB1, MGAT4B and PFN1 there is one pose correctly positioning the **X1** close to the FFF site (**Figure S13B**).

In about 39% of these validated poses, the reactive diazirine group of **X1** was oriented directly toward the cross-linked site, further supporting the physical plausibility of the docking models.

Analysis of all modeled RNA–ligand complexes revealed consistent modes of molecular recognition. Nearly every structure contained at least two hydrogen bonds between **X1** and its RNA target (average = 3 ± 0.3 hydrogen bonds per complex), primarily involving nucleobases rather than the sugar–phosphate backbone (**Figure S14A-C**). Hydrophobic and van der Waals interactions were also commonly observed, reinforcing the importance of shape complementarity within RNA pockets. π-stacking interactions were present in approximately one-third of the docked complexes, most often involving purine bases such as adenine and guanine, whose extended π-surfaces favor aromatic stacking (**Figure S14C**). These findings suggest that **X1** recognizes RNA through a combination of base-specific hydrogen bonding, hydrophobic packing, and π-stacking stabilization, with a preference for purine-rich internal loops and junctions.

Consistent with these observations, the majority of modeled binding motifs corresponded to symmetric or asymmetric internal loops and tetraloops, whereas single-nucleotide bulges were less frequent (**Figure S15 A-B**). This pattern indicates that **X1** preferentially engages structured RNA elements that provide both geometric definition and accessible nucleobase surfaces for interaction.

Detailed examination of individual targets further supported these conclusions. In the *MPP7* transcript, the lowest-energy binding poses placed **X1** within a 4×4-nucleotide internal loop, forming multiple hydrogen bonds (notably with U23 and C24) and hydrophobic contacts with neighboring purines (**Figure S17**). The orientation of the diazirine cross-linking group was within 5 Å of the experimentally mapped site, consistent with Chem-CLIP data. Similarly, for the SSC4D transcript, **X1** was predicted to bind within a three-way junction stabilized by hydrogen bonding to A7 and hydrophobic contacts with surrounding purines (G6, G8, A24) (**Figure S18**).

In summary, molecular docking and cross-linking analyses suggest that **X1** recognizes structured, purine-rich RNA motifs through a combination of hydrogen bonding, hydrophobic, and aromatic interactions. These models provide a structural rationale for **X1**’s transcriptome-wide selectivity and define key interaction features that can guide the design of improved RNA-targeting small molecules. However, these conclusions are predictive, and the precise mode of binding will require further experimental and structural validation through higher-resolution biophysical and biochemical analyses. Complete computational and parameter details are described in the Supporting Information.

### 3D structure modeling of X1 analogs bound to the *MPP7* binding site by molecular docking

To gain insight into why only **X1-D1 and X1-D2** competed with **X1** for binding to *MPP7* mRNA in cells, molecular docking was carried out. For each compound, the top three docking poses with the lowest predicted binding energies were extracted and analyzed (**Figure S19-22**). Derivatives **X1-D1 – X1-D4** had similar or lower binding energies than parent compound **X1** (−6.97 ± 0.1 kcal/mol), −7.41 ± 0.07 (kcal/mol), −7.99 ± 0.03 (kcal/mol), −6.99 ± 0.15 (kcal/mol), and −6.85 ± 0.41 (kcal/mol), respectively (**Table S5**). Further analysis of the top-ranked docking poses for **X1-D1 – X1-D4** each have a binding pose that is like **X1** pose #3, making hydrogen bonds with U23, C24 and a combination of vDW (van Der Waals) and hydrophobic interactions with G22, U23, A9. In contrast, the remaining analogs had significantly lower binding energies, averaging −5.60 ± 0.22 kcal/mol (**Table S4**) and have localized to an alternate binding pocket, primarily the 2×3 nt asymmetric internal loop (**Figure S23-28**). Interestingly, only **X1-D1** and **X1-D2** compete with **X1** for binding *MPP7* mRNA, both of which have lower binding energies than the parent compound, while **X1-D3** and **X1-D4** have similar binding energies, and the remaining analogs have significantly less favorable binding energies (**Table S4**).

For comparison, molecular docking was also carried for the **X1** analogs and the *SSC4D* mRNA binding site (**Figure S29-S38**). As for the *MPP7* docking studies, the top three docking poses for each analog with the lowest predicted binding energies were extracted and analyzed. Derivatives **X1-D1** – **X1-D4** exhibited binding energies averaging –7.49 ± 0.04 kcal/mol, significantly more favorable than that of the parent compound **X1** (−6.47 ± 0.1 kcal/mol) (**Table S5**). In contrast, compounds **X1-D5** – **X1-D10** showed similar binding energies as parent **X1**, averaging –6.43 ± 0.11 kcal/mol (**Table S5**). Further analysis of the top-ranked docking poses revealed that compounds **X1-D1**– **X1-D4** bind in the same binding pocket as **X1** molecule (**Figure S29-32**) making hydrogen bond with A7 and a combination of vdW and hydrophobic interactions with G6, A24, A7, G6 and G8 preserving key molecular contacts. In contrast, all other analogs, except for **X1-D7**, bound to a different pocket (**Figure S33-38**). This suggests that the modifications in these analogs not only reduced binding affinity but also altered the binding site preference.

## DISCUSSION

This study establishes a general framework for decoding how small molecules recognize RNA in living cells and for translating those interactions into functional outcomes. By combining chemoproteomic mapping, transcriptome-wide sequencing, machine learning, and rational degrader design, we reveal that RNA–small molecule recognition follows definable physicochemical and structural principles rather than idiosyncratic chance.

A major insight emerging from these data is that small molecules preferentially engage structured regions within untranslated regions (UTRs) of mRNAs—domains already enriched in regulatory elements and evolutionary conservation. This bias underscores that RNA structure, not abundance alone, governs ligandability in cells. Moreover, integration of thermodynamic modeling with experimental mapping revealed that ligandable sites coincide with unusually stable, low z-score folds, consistent with functional selection pressures on RNA architecture. These observations extend the concept of *druggable structure* from proteins to RNA, providing an empirical basis for identifying functional RNA folds that can be engaged by chemical matter.

The machine-learning analysis unifies diverse chemotypes into quantifiable rules for RNA recognition. Aromaticity, heteroatom content, charge distribution, and defined molecular shape—rather than a single scaffold—emerged as shared determinants of binding. Importantly, models integrating multiple chemical fingerprints and three-dimensional descriptors achieved predictive power across distinct compound classes, offering a path toward prospective discovery of RNA binders by computation. Together, these data define a chemical grammar of RNA recognition that can guide rational design rather than reliance on serendipity.

Conversion of these fragments into ribonuclease-targeting chimeras (RiboTACs) further revealed that broad binding does not imply broad degradation. Despite interacting with hundreds of transcripts, only two targets—MPP7 and SSC4D—were cleaved, underscoring the inherent selectivity conferred by spatial proximity between the RNA-binding site and an RNase L–accessible pyrimidine motif. By coupling unbiased mapping with cellular degradation assays, we show that selective RNA knockdown can be achieved through design rather than random screening. Structural optimization of the X1 RNA-binding module produced an MPP7-selective RiboTAC that reduced oncogenic MPP7 expression and suppressed migration of triple-negative breast cancer cells— demonstrating that chemical fine-tuning of RNA recognition can reprogram biological specificity.

Collectively, these findings establish a scalable and generalizable paradigm for studying RNA–small molecule interactions in living systems. The integration of high-throughput covalent profiling, thermodynamic and structural modeling, and predictive computation creates a roadmap for discovering, modeling, and functionalizing ligandable RNA structures. More broadly, this work defines actionable rules of RNA recognition that bridge chemical biology and transcriptomics—laying the groundwork for systematic development of RNA-targeted small molecules and degraders that modulate disease-relevant RNAs with precision.

## CONCLUSION

This work defines a data-driven, experimentally validated roadmap for discovering, modeling, and functionalizing small-molecule interactions with RNA in living cells. By integrating covalent chemptranscriptomcs, transcriptome-wide mapping, and machine learning, we show that RNA–ligand recognition follows reproducible physicochemical principles rather than stochastic binding. The discovery that small molecules preferentially engage thermodynamically stable structures within mRNA untranslated regions provides a molecular basis for identifying ligandable RNA elements across the transcriptome.

Just as large-scale chemoproteomic platforms and predictive tools such as AlphaFold have revolutionized protein ligandability and structure prediction, this work establishes a parallel foundation for RNA. Yet, the field remains at an early stage. Developing computational and experimental frameworks that can translate transcriptome-wide ligandable patterns into accurate three-dimensional RNA–small molecule interaction models remain one of the key challenges ahead. Addressing this will require integration of chemical biology, structural modeling, and high-throughput profiling to illuminate how RNA folds and functions can be chemically controlled in cells. They will also require that the predicted ligand-RNA structures are further validated by, for example, cryoEM or other structural analysis to obtain complex structural information. These studies open the path to systematic exploration of RNA structures as druggable entities, predictive modeling of RNA binding pockets, and the rational design of small-molecule modulators and degraders that can reprogram RNA biology with precision.

## MATERIALS AND METHODS

### Cell Culture

All cells were cultured at 37 °C with 5% CO_2_. MDA-MB-231 cells (ATCC, HTB-26) were cultured in 1× RPMI Medium 1640 (Sigma Aldrich, Cat# R7388) supplemented with 10% (v/v) fetal bovine serum (FBS; Sigma Aldrich, Cat# F2442) and 1× Penicillin/Streptomycin (Gibco, Cat# 15140122). Cells were used up until passage 20 and verified to be free of mycoplasma contamination (PromoKine, Cat# PK-CA91-1024) before performing cellular experiments.

To treat cells with compounds, the growth medium was replaced with fresh growth medium that contained the compound at the treatment concentration with a final concentration of 0.1% (v/v) DMSO in all samples, except for C-Chem-CLIP studies were the final concentration of DMSO was 0.2% (v/v). For antisense oligonucleotide^66^ treatment, Lipofectamine RNAiMAX Reagent (Invitrogen, Cat# 13778150) was used for transfection following manufacturer’s recommendations. For 100 mm diameter dishes, 12 mL of growth medium was used per dish. For 6-well, 12-well, and 96-well plates, the volumes of growth medium used were 2 mL, 1 mL, and 0.1 mL per well, respectively.

### In vitro Screening of FFFs That Cross-link to Human RNA

Total RNA was isolated from WT MDA-MB-231 cells cultured in 100 mm diameter dishes as described in **Cell Culture** by using a Quick-RNA Miniprep Kit (Zymo, Cat# R1054) per manufacturer’s protocol, including on-column DNase I treatment. For each biological replicate, 10 µg of total RNA in 100 µL of 1× Folding Buffer (20 mM HEPES, pH 7.5, 150 mM NaCl, and 5 mM KCl) was folded by heating at 90 °C for 3 min followed by cooling to room temperature on the bench top. The sample was incubated with the compound (100 µM, 0.2% (v/v) DMSO) at room temperature for 1 h, followed by irradiation with 350-365 nm UV light for 10 min in a photo-crosslinker (UV Stratalinker 2400 with Eiko F15T8/BL bulbs) while leaving the cap of the tubes open. Each sample was supplemented with TAMRA (5-carboxytetramethylrhodamine) azide dye (200 µM) and a pre-mixed click reaction solution (pH = 7.5) containing CuSO_4_ (5 µL of 10 mM aqueous solution), THPTA (5 µL of 50 mM aqueous solution) and sodium ascorbate (5 µL of 250 mM aqueous solution). After mixing well by pipetting, the samples were incubated at 37 °C for 1 h, followed by the addition of 11.5 µL of 3 M sodium acetate (pH = 5.2) and 400 µL of 100% ethanol. The samples were incubated at −80 °C for 16 h, followed by centrifugation at 14,000 rpm at 4 °C for 10 min. The supernatant was carefully removed by pipetting, and the precipitated RNA was washed with 400 µL of 70% (v/v) ethanol, followed by centrifugation at 14,000 rpm at 4 °C for 10 min. After removing the supernatant by pipetting, the RNA was dissolved in 20 µL of 1× Loading Dye (NEB, Cat# B7024S) and separated on a 1.5% (w/v) agarose gel in 1× TBE buffer (100 V, 1 h. The TAMRA signal was imaged by a Typhoon FLA 9500 variable mode imager (GE Healthcare Life Sciences), followed by SYBR Green staining to visualize total RNA. The TAMRA signal was normalized to the SYBR Green signal in each lane to normalize for the loading of total RNA. The ratio of TAMRA/SYBR Green signal for the **Control probe** was set to 1.

### Chem-CLIP Profiling by Global RNA-seq

WT MDA-MB-231 cells were seeded into 100 mm diameter dishes at ∼70% confluency. After allowing the cells to fully adhere overnight (16 h), the cells were treated with vehicle (DMSO, 0.2% v/v) or the compound of interest. For competitive Chem-CLIP with **X1**, cells were treated with growth medium containing **X1** derivatives lacking diazirine and alkyne groups at the indicated concentrations and the **X1** probe (20 µM), that is, co-treatment; the final DMSO concentration was 0.2% (v/v) in all samples. After 16 h, the growth medium was removed, and cells were washed with 1× DPBS twice (5 mL each time), followed by adding 2 mL of 1× DPBS to each dish to cover the entire bottom. Cells (no lid) were irradiated with 350-365 nm UV light for 10 min in a photo-crosslinker (UV Stratalinker 2400 with Eiko F15T8/BL bulbs). After UV irradiation, the liquid was removed by pipetting, and total RNA was extracted by using Quick-RNA Miniprep Kit (Zymo, R1054) per manufacturer’s protocol, including the on-column DNase I treatment. The RNA samples were random fragmented to ∼150 nt lengths by using an NEBNext Magnesium RNA fragmentation module (NEB, Cat# E6150S) per manufacturer’s recommendations. The fragmented RNA was purified by using RNAClean XP beads (Beckman Coulter, Cat# A66514) by following the manufacturer’s instructions and eluted in 50 µL of RNase-free water. An aliquot of RNA samples (200 ng) was saved and stored at −80 °C for RNA-seq library preparation to measure target abundance before Chem-CLIP enrichment.

For each biological replicate, 10 µg of RNA was added to 100 µL of azide-disulfide agarose beads (Click Chemistry Tools, Cat# 1238-2) pre-washed with 200 µL of 25 mM HEPES (pH 7.0). A click reaction solution (pH 7.5) was added to each sample, which was prepared by mixing 30 µL of 250 mM sodium ascorbate, 30 µL of 10 mM CuSO_4_, and 30 µL of 50 mM THPTA. The samples were then incubated while rotating at 37 °C for 1 h, followed by centrifugation for 1 min at 160 × g. After removing the supernatant by pipetting, the beads were washed six times (500 µL each) with Washing Buffer (10 mM Tris-HCl, pH 7, 4 M NaCl, 1 mM EDTA, and 0.05% (v/v) Tween-20). The beads were then incubated with 100 µL of Elution Buffer (50 mM TCEP and 100 mM K_2_CO_3_, pH 10) at 37 °C for 30 min, followed by the addition of 100 µL iodoacetamide (200 mM). After another 30 min incubation at 37 °C, the samples were centrifuged for 1 min at 160 × g, and the supernatant were transferred to a new microcentrifuge tube. The RNA eluted in the supernatant was purified by using RNA CleanXP beads (Beckman Coulter, Cat# A66514) per manufacturer’s recommendations and eluted in 30 µL of RNase-free water.

To prepare the library for RNA-seq, the RNA samples before and after Chem-CLIP enrichment (200 ng per sample) were depleted of ribosomal RNA with an NEBNext rRNA Depletion Module (NEB, Cat# E6310) per manufacturer’s recommendations. The library was prepared using a NEBNext Ultra II Directional RNA kit (NEB, Cat# E7760) by following the manufacturer’s protocols. Briefly, first strand cDNA synthesis was completed by using random priming, followed by second strand synthesis by using dUTP to replace dTTP. The 3’ ends of cDNA were end repaired and adenylated, followed by adaptor ligation. The strand information of the RNA was preserved by using USER (Uracil-specific excision reagent) to degrade the second strand containing dUTP. The cDNA was then PCR amplified with barcoded Illumina-compatible primers to obtain the final libraries, which were loaded into a NextSeq 500 v2.5 flow cell and sequenced with 2 x 40 bp paired-end chemistry (> 20 million reads received per sample).

For data analysis, which was completed as previously described,^18, 67^ the raw sequencing fastq files were aligned to the human genome by STAR (all samples contain >80% uniquely mapped reads by STAR alignment).^68^ The output files after alignment were processed by Genrich (available at github.com/jsh58/Genrich) to identify the peaks (*i.e.*, regions of enrichment) by comparing the sample after Chem-CLIP enrichment to the sample that was not subjected to enrichment steps. A minimum enrichment of 1.5-fold and a minimum read count of 5 was applied to all identified peaks to manually remove low-confidence targets. The binding sites of the small molecules within each target were determined by the end of each peak, whose chromosomal coordinates were reported by Genrich based on statistical confidence. No unique codes were used for these analyses. To model the RNA structures around the binding site, the entire transcript was folded by using ScanFold^69^ (window size = 120 nucleotides, randomization = 100) by following the instructions on the user interface.

### Construction and Validation of RNase L CRISPR Knockdown MDA-MB-231 Cells

The CRISPR-edited MDA-MB-231 cell lines used in this study were previously reported and characterized.^55^ To construct these cell lines, lentiviral constructs containing Cas9 and guide RNA (gRNA) targeting RNase L mRNA (ATTTGTACTGCGTTATGCAG) or a control non-targeting gRNA (GGAGCGCACCATCTTCTTCA) were purchased (CAHS101, Transomic Tech) and transfected into HEK293T cells (ATCC, CRL-3216). The virus was harvested and transduced into WT MDA-MB-231 cells in the presence of 6 mg/mL polybrene (Millipore, Cat# TR-1003-G). The selection of successfully transduced cells was performed by treating cells with 1 µg/mL puromycin (Gibco, Cat# A1113802), which was no longer added during compound- or vehicle-treatment. The mRNA and protein levels of RNase L in the CRISPR knockdown cell line were reduced by ∼60% compared to WT MDA-MB-231 cells, as measured by RT-qPCR and Western blot, respectively, as previously published.^55^ The mRNA and protein levels of RNase L in the control guide CRISPR cells are similar to the levels in wild-type MDA-MB-231 cells (<10% difference), as measured by RT-qPCR and Western blot, respectively, as previously published.^55^

### Measuring mRNA Abundance by RT-qPCR

Cells were seeded into 12-well plates at ∼40% confluency and allowed to adhere to the plate surface for 16 h. They were then treated with vehicle (DMSO, 0.1% (v/v)) or compound as described in **Cell Culture**. After 48 h, total RNA was extracted by using a Quick-RNA Miniprep Kit (Zymo, Cat# R1054) per manufacturer’s protocol, including the on-column DNase I treatment. Reverse transcription (RT) was performed by using a Qscript cDNA synthesis kit (QuantaBio, Cat# 95047) per manufacturer’s protocols with 500 ng of input total RNA in a total volume of 20 µL. The qPCR amplification was performed in 384-well plates in 1× Power SYBR Green Master Mix (Life Technologies, Cat# A46109) containing 570 nM of the corresponding forward and reverse primers (Table S1) using a Quant Studio 5 Real-Time PCR system. For each biological replicate, 2 µL of cDNA product from the RT-step was added into 34 µL of reaction mixture, followed by aliquoting into three technical replicates (10 µL per replicate). The relative abundances of mRNA were calculated by using the ΔΔCt method as described previously.^70^

### Chem-CLIP in Cells by RT-qPCR

The experimental workflow is same as described in **Chem-CLIP Profiling by Global RNA-seq** except there is no random fragmentation step, as it can impair analysis by RT-qPCR. After Chem-CLIP enrichment, RT-qPCR amplification was performed by following the same procedure as described in **Measuring mRNA Abundance by RT-qPCR.** The relative abundances of mRNA were calculated by using the ΔΔCt method as previously described.^70^ To calculate the fold of enrichment, the following equation was used:

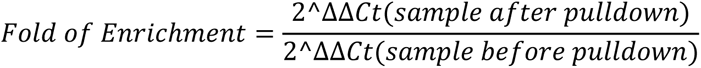

### Library Preparation and Data Analysis for Global RNA-seq

Cells were seeded into 12-well plates at ∼40% confluency and allowed to fully adhere overnight (16 h). Cells were treated with vehicle (0.1% (v/v) DMSO) or compound for 48 h as described in **Cell Culture**. After treatment, total RNA was harvested as described in **Measuring mRNA abundance by RT-qPCR**. The RNA-seq library preparation was performed by following the same procedure as described in **Chem-CLIP profiling by global RNA-seq**. For data analysis, the raw sequencing fastq files were aligned to the human genome by STAR (all samples contain >80% uniquely mapped reads by STAR alignment).^68^ The output files after alignment were processed by featureCounts^71^ and the differential gene analysis was performed by Deseq2 (minimum read count of 10 as the default cut-off)^72^ in R Studio. No unique codes were used for these analyses.

### Measuring Protein Abundance by Western Blotting

Cells were seeded into 6-well plates to afford ∼40% confluency, and the cells were allowed to adhere fully to the plate surface for 16 h. The cells were then treated with vehicle (DMSO; 0.1% (v/v)) or compound as described in **Cell Culture**. After a 48 h treatment period, the cells were washed twice with ice-cold 1× DPBS. The DPBS was removed, and total protein was harvested by adding 100 µL of Mammalian Protein Extraction Reagent (M-PER, Thermo Scientific, Cat# 78501) to each well. Protein concentration was quantified by using a Micro BCA Protein Assay Kit (Pierce, Cat# 23235) following the manufacturer’s protocol. The proteins were separated by gel electrophoresis using 10% SDS-polyacrylamide gel (30 µg total protein loaded per lane) in 1× Running Buffer (50 mM MOPS, 50 mM Tris-base, pH 7.5, 1 mM EDTA, and 0.1% (w/v) SDS) at 120 V for 1 h. The proteins were then transferred to a PVDF membrane (0.45 µm, Cytiva) at 300 mA for 1 h in 1× Transfer Buffer (25 mM Bicine, 25 mM Tris-base, pH 7.5, 1:4 MeOH:H_2_O).

The membrane was blocked for 1 h with 5% (w/v) non-fat dry milk in 1× TBST [1× TBS (20 mM Tris-HCl, pH 7.6, and 150 mM NaCl) containing 0.1% (v/v) Tween-20] at room temperature. The membrane was incubated with the primary antibody for MPP7 protein (ProteinTech, Cat# 12983-1-AP) or SSC4D (Sigma Aldrich, Cat# HPA062611) using 1:1000 dilution in 1× TBST containing 5% (w/v) non-fat dry milk at 4 °C for 16 h. The membrane was then washed by 1× TBST four times (5 min each wash) and incubated with the secondary antibody (Cat# 7074S, Cell Signaling Technology, CST) with 1:3000 dilution in 1× TBST containing 5% (w/v) non-fat dry milk at room temperature for 2 h. The membrane was washed with 1× TBST four times (5 min each wash) and imaged by using a SuperSignal West Pico Chemiluminescent Substrate (Pierce, Cat# 34580) per manufacturer’s protocol.

After imaging, the membrane was stripped by washing in 1× Stripping Buffer (200 mM glycine, pH 2.2, 4 mM SDS, 1% (v/v) Tween 20) at room temperature for 20 min. The membrane was blocked again in 1× TBST containing 5% (w/v) non-fat dry milk for 1 h at room temperature. The blot was then incubated with an antibody for β-actin protein (Abcam, Cat# 8226) at a 1:5000 dilution in 1× TBST containing 5% (w/v) non-fat dry milk. The membrane was imaged by following the same procedure as described in the paragraph above. Expression levels of proteins were quantified based on band intensity normalized to β-actin by using ImageJ.^73^

### Boyden Chamber Migration Assay

MDA-MB-231 cells were seeded in 60 mm diameter dishes at ∼40% confluency, allowing cells to fully adhere at the incubator for 16 h. The cells were then treated with compound or DMSO (vehicle, 0.1% (v/v)) for 40 h as described in **Cell Culture**. After treatment, serum starvation was performed by replacing the growth medium with fresh medium that contains the same concentration of compound or DMSO but lacking FBS for 8 h. The treated, serum-starved cells were trypsinized and then seeded into ThinCert 24-well inserts (GBO, Cat# 662638) with 8 µm pores at 50,000 cells per well (100 µL per insert). Fresh growth medium containing 10% (v/v) FBS was added to a 24-well plate (600 µL each well), and the inserts containing the treated, serum-starved cells were placed on top, allowing the medium to cover the bottom of the insert. After 16 h, all medium was removed by pipetting, and cells in the insert were gently washed twice by 1× DPBS (600 µL per well each time). The cells were fixed by addition of 3% (w/v) paraformaldehyde (PFA) in 1× DPBS (600 µL per well) at room temperature for 30 min. After fixing, the cells were stained with Crystal Violet (10 mg/mL; 4:1 H_2_O:MeOH, 600 µL per well) at room temperature for 20 min. A cotton swab was used to gently remove non-migratory cells from the top of the insert. The inserts were then imaged by using a bright-field microscope (10X magnification, 3 views per well), and the number of migratory cells was counted manually.

### Chemotype analysis

A machine learning pipeline integrating cheminformatics and feature interpretation was developed to classify RNA-binding and non-binding small molecules. Molecules were represented by their SMILES strings and encoded into three complementary molecular fingerprints—Morgan, RDKit, and atom-pair fingerprints— capturing local atomic environments, bonding topology, and interatomic distances, respectively. These representations together provided a comprehensive description of chemical structure for modeling RNA recognition. To assess predictive performance, each fingerprint was first evaluated independently, yielding modest discriminative power (area under the ROC curve [AUC] = 0.60–0.68). Therefore, an ensemble approach combining all three fingerprints was used as input for a Random Forest classifier, chosen for its robustness to high-dimensional data. The dataset of 200 compounds (23 binders, 177 non-binders) was partitioned into five stratified folds, ensuring balanced class representation. Four folds were used for model training and one for testing, with the process repeated across all folds for cross-validated performance estimates. To correct for class imbalance and reduce model bias, three strategies were implemented: (i) SMOTE (Synthetic Minority Over-sampling Technique) to generate synthetic binders through interpolation between structurally similar compounds (Tanimoto = 0.8–0.9); (ii) Tomek Links to remove overlapping non-binders near class boundaries; and (iii) stratified K-fold cross-validation to maintain class balance during training and testing. Hyperparameter optimization was performed via grid search, systematically testing combinations of six key Random Forest parameters to identify configurations maximizing predictive accuracy while minimizing overfitting. Cross-validated performance scores were visualized as heatmaps, identifying optimal parameter regions. Model predictions were evaluated using receiver operating characteristic (ROC) and confusion matrix analyses. To refine classification, the optimal decision threshold was determined by maximizing the F1-score, which balances precision and recall. Rather than applying the default 0.5 threshold, model performance was assessed across thresholds, with 0.34 identified as optimal. This threshold achieved the best trade-off between binder sensitivity and non-binder specificity. The final Random Forest model achieved a mean AUC of 0.92 ± 0.04, demonstrating excellent discrimination between binders and non-binders. Confusion matrix analysis confirmed strong classification accuracy (34/35 true negatives and 11/14 true positives). Feature contributions were quantified using SHapley Additive exPlanations (SHAP) values, which assign importance scores to each fingerprint bit, indicating how specific molecular substructures influence RNA binding probability. Comparative enrichment analysis of features unique to binders revealed distinct chemotypes and substructural motifs that favor RNA recognition (see note S1 for details).

### Automated 3D RNA Modeling and Molecular Docking

To investigate the structural basis of X1–RNA recognition, a multi-step computational workflow integrating secondary structure prediction, 3D modeling, and molecular docking was implemented. Minimum-free-energy secondary structures for 106 transcripts (58 genes) identified by Chem-CLIP-Map-Seq were first obtained using ScanFold. Corresponding 3D RNA models were generated with the Fragment Assembly of RNA with Full-Atom Refinement v2 (FARFAR2) protocol in Rosetta, which assembles RNA structures from experimentally derived fragments followed by all-atom refinement. Hundreds of candidate (“decoy”) conformations were produced per transcript and ranked by Rosetta energy units (REU); the ten lowest-energy models (within 2 REU of the global minimum) were retained to represent the conformational ensemble. Each RNA model was subjected to blind docking with AutoDock-GPU using X1 as the ligand to identify potential binding pockets across the entire RNA surface. Although Chem-CLIP-Map-Seq data provided the known binding region, these sites were not used as restraints. For validation, predicted poses were compared with experimentally identified cross-linked nucleotides from Chem-CLIP studies. For each transcript, the three lowest-energy docking poses were selected, and the Euclidean distance between the ligand and cross-linked residue centers of mass (COMs) was calculated. Poses with COM distances ≤ 10 Å were considered successful (68.7 % success across all RNAs). For geometrically valid poses, the diazirine orientation was examined by defining a vector from the diazirine center to the local site COM (± 5 nt window). Poses with ≤ 8 Å separation were deemed spatially consistent with cross-linking data (39 % success) (see note S2-3 for details).

### Molecular descriptor calculations, binding motif and interaction extraction

Approximately 1,800 two- and three-dimensional molecular descriptors were calculated for each fragment using the Mordred descriptor engine. Statistical differences between binder and non-binder fragments were assessed using the Mann–Whitney U test to identify descriptors with significant variation between the two groups. Descriptor categories included physicochemical, topological, and 3D geometric properties such as partial surface area, charge distribution, and moment of inertia, providing a comprehensive representation of molecular features relevant to RNA recognition (see note S4 for details). RNA secondary structures were analyzed using a Python pipeline that parses dot-bracket notation from Excel input, identifies motifs such as internal loops and tetraloops, maps cross-linked nucleotides, and generates quantitative and graphical summaries of motif distributions across transcripts (see note S5 for details). RNA–ligand interaction profiling was performed using a Python-based automated pipeline. The workflow (i) separated RNA and ligand coordinates from complex .pdb files using PyMOL^74^, (ii) converted ligand structures to .sdf format with Open Babel^75^, and (iii) executed the fingeRNAt^76^ toolkit to detect and classify RNA–ligand interactions, including hydrogen bonding, hydrophobic, and π-stacking contacts (see note S6 for details).

### Statistical Analysis

The statistical analyses were performed by using GraphPad Prism (version 10.3.0). Detailed information for each statistical analysis and its corresponding number of biological replicates was indicated in figure legends.

## DATA AVAILABILITY

RNA-seq data were deposited at Mendeley Data (doi: 10.17632/kkhxcdz3mc.1). All data can be found at Supplemental Documents associated with this manuscript.

## ACKNOWLEDGMENT

This work was supported by the National Institutes of Health (R01 CA249180A to M.D.D.; R01 GM133810 to W.N.M.; and F31 CA257090 to W.B.R.), and the Muscular Dystrophy Association Development Grant 963835 (to A.T.).

## CONFLICT OF INTEREST

M.D.D. is the founder of Expansion Therapeutics. M.D.D. and J.L.C. are co-founders of Ribonaut Therapeutics.

## Notes

### Competing Interest Statement

MDD Is a founder of Expansion and Ribonaut therapeutics. JLC-D is a founder of Ribonaut therapeutics.

