## Supplemental data for "Live-Cell Covalent Profiling Reveals Principles of RNA-Small Molecule Recognition across the Human Transcriptome"

### Supplementary Tables & Figures

**Table S1. Sequences of oligonucleotides used in this study.** "F" denotes a forward primer while "R" denotes a reverse primer.

| Oligonucleotide | Sequence 5' to 3' | Experiment |
|---|---|---|
| SSC4D-F | GTCCCCAGCTGGATGAGAAG | qPCR |
| SSC4D-R | GCCAGTGGCAGGAGGAGA | qPCR |
| MPP7-F | GTAGACAACGTGGCTGCAGGCT | qPCR |
| MPP7-R | CTGGCATGATGCAAGGTGTAGG | qPCR |
| ACTB-F | CATGTACGTTGCTATCCAGGC | qPCR |
| ACTB-R | CTCCTTAATGTCACGCACGAT | qPCR |
| MPP7-ASO[a] | **GAA**TTCCAATTCAA**CAG** | Cellular assays |
| Control-ASO[a] | **GUG**AGGG**UCA** | Cellular assays |

[a] Linkages of all nucleotides are phosphorothioate.
2'-O-Methoxyethyl (MOE) nucleotides are indicated in bold.
All oligonucleotides in this table were purchased from Integrated DNA Technologies (IDT).

| **Table S2. Abbreviations and descriptions of statistically significant molecular descriptors that distinguish binding and non-binding fragments.** | |
|---|---|
| **Descriptor** | **Description** |
| Mor01p | 3D pattern emphasizing atoms with high polarizability at signal 01 |
| Mor01v | 3D pattern emphasizing atoms with large van der Waals volumes at signal 01 |
| Mor02se | 3D signal capturing electronegativity-weighted atom positions (signal 02) |
| Mor02v | 3D pattern showing how atoms with large van der Waals volumes are arranged (signal 02) |
| Mor08 | Basic 3D spatial signal of the molecule (signal 08, unweighted) |
| Mor08se | 3D spatial signal emphasizing atoms with higher electronegativity (signal 08) |
| Mor16 | Basic 3D shape pattern of the molecule at signal 16 (unweighted) |
| Mor16p | 3D signal focused on more polarizable atoms and their spatial distribution (signal 16) |
| Mor16v | 3D signal showing how bulky atoms (van der Waals volume) are arranged (signal 16) |
| Mor16se | 3D signal emphasizing electronegative atoms at signal 16 |
| Mor21v | 3D signal highlighting spatial pattern of large atoms (signal 21) |
| Mor22p | 3D signal showing arrangement of polarizable atoms (signal 22) |
| Mor22v | 3D signal highlighting bulky atoms' positions (signal 22) |
| Mor23m | 3D signal weighted by atom mass (signal 23) |
| Mor28 | General 3D signal of the molecule (signal 28, unweighted) |
| Mor28m | 3D pattern focused on heavier atoms (signal 28) |
| Mor28p | 3D signal showing positions of polarizable atoms (signal 28) |
| Mor28se | 3D signal showing how electronegative atoms are arranged in space (signal 28) |
| Mor28v | 3D signal highlighting spatial distribution of bulky atoms (signal 28) |
| Mor32 | Basic 3D molecular signal (signal 32, unweighted) |
| Mor32se | 3D pattern emphasizing electronegative atoms (signal 32) |
| FNSA5 | Fraction of negative surface area relative to total surface area (bin 5) |
| FPSA2 | Fraction of positive surface area weighted by partial charges (bin 2) |
| PPSA2 | Total positive surface area weighted by partial charges (bin 2) |
| WPSA2 | Weighted positive surface area using partial atomic charges (bin 2) |
| WPSA4 | Weighted positive surface area using partial atomic charges (bin 4) |
| WPSA5 | Weighted positive surface area using partial atomic charges (bin 5) |
| DPSA2 | Difference between total positive and negative surface areas (bin 2) |
| RPCS | Relative negative partial surface area — how much of the surface is negative compared to total charge |
| [a] Statistically significant descriptors as calculated with the boundaries of ***: $1 \times 10^{-4} < p \leq 1.00 \times 10^{-3}$ and ****: $p \leq 1 \times 10^{-4}$. | |

| **Table S3. Grid search parameters for Random Forest model optimization. Summary of the hyperparameter ranges used in the grid search to tune the Random Forest classifier.** | |
|---|---|
| N_estimators | 100,200 |
| Max_depth | 5,10, None |
| Min_samples_split | 2,5 |
| Max_features | 2,4,6 |
| bootstrap | True |
| Min_samples_leaf | 2,4,6 |

**Table S4. Binding energy of X1 and XD1-D10 derivatives against MPP7 mRNA. The first lowest energy docked poses were extracted with an energy difference less than 1 kcal/mol.**

| **X1 derivative** | **ΔG (kcal/mol)** | **X1 derivative** | **ΔG (kcal/mol)** |
|---|---|---|---|
| X1 | -6.97±0.1 | X1D6 | -5.71±0.67 |
| X1D1 | -7.41±0.07 | X1D7 | -5.75±0.27 |
| X1D2 | -7.99±0.03 | X1D8 | -5.78±0.05 |
| X1D3 | -6.99±0.15 | X1D9 | -5.63±0.14 |
| X1D4 | -6.85±0.41 | X1D10 | -5.18±0.0 |
| X1D5 | -5.56±0.2 | | |

**Table S5. Binding energy of the first three lowest energy poses of X1 and X1 derivatives to SSC4D (1 to 10).**

| **X1 derivative** | **ΔG (kcal/mol)** | **X1 derivative** | **ΔG (kcal/mol)** |
|---|---|---|---|
| X1 | -6.47±0.1 | X1D6 | -6.38±0.35 |
| X1D1 | -7.02±0.02 | X1D7 | -6.68±0.09 |
| X1D2 | -7.61±0.11 | X1D8 | -6.75±0.0 |
| X1D3 | -7.67±0.0 | X1D9 | -6.36±0.12 |
| X1D4 | -8.12±0.14 | X1D10 | -5.98±0.01 |
| X1D5 | -6.05±0.14 | | |

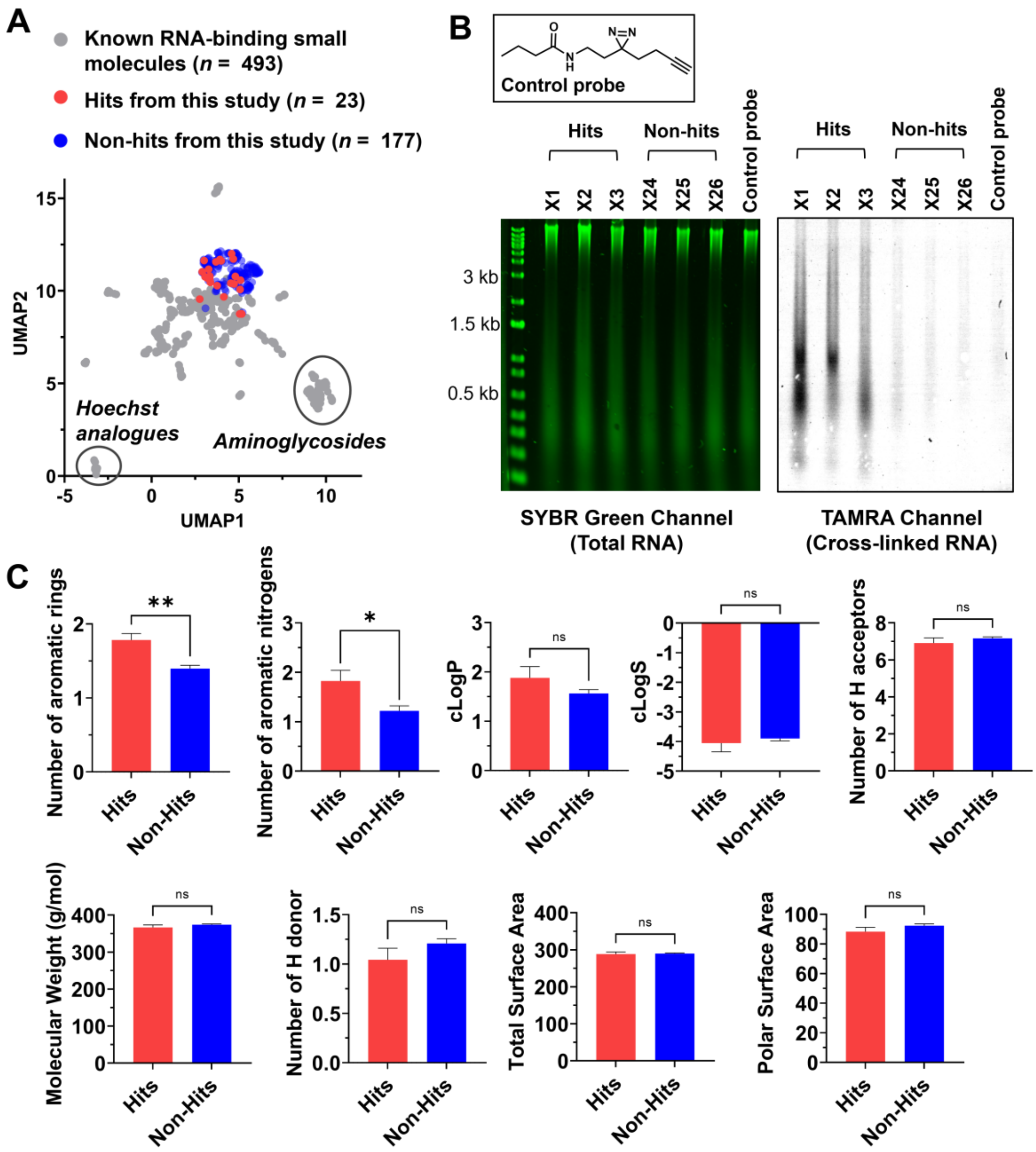


**Figure S1. In vitro screening of functionalized small molecules that interacting with human transcripts.** (A) UMAP (Uniform Manifold Approximation and Projection for Dimension Reduction) analysis comparing published known RNA-binding small molecules[1, 2] to the 200 small molecules used in this study. (B) Representative gel images of the *in vitro* screening of small molecules that interact with total human RNA isolated from WT MDA-MB-231 cells. After cross-linking the fragments to bound RNAs via UV irradiation, an azide-functionalized fluorescent dye, TAMRA (5-carboxytetramethylrhodamine), was conjugated to the alkyne handle of the FFFs by click chemistry.[3] Fluorescence signal from the TAMRA dye (Excitation 550 nm, Emission 580 nm) was used to visualize the RNA bound by small molecules. This signal was normalized to the total amount of RNA loaded into each well, afforded by post-SYBR

Green staining. The normalized TAMRA:SYBR Green signal was set to 1 for the control probe (see Methods for details and equations). Binding fragments, or hits, are defined as small molecules with normalized TAMRA signal that is >1σ (2.5) than the average (2.9) of all screened small molecules. (C) Physicochemical properties of small molecule hits (n = 23) and non-hits (n = 177) from *in vitro* screening. Data in panel C are reported as average ± S.E.M.

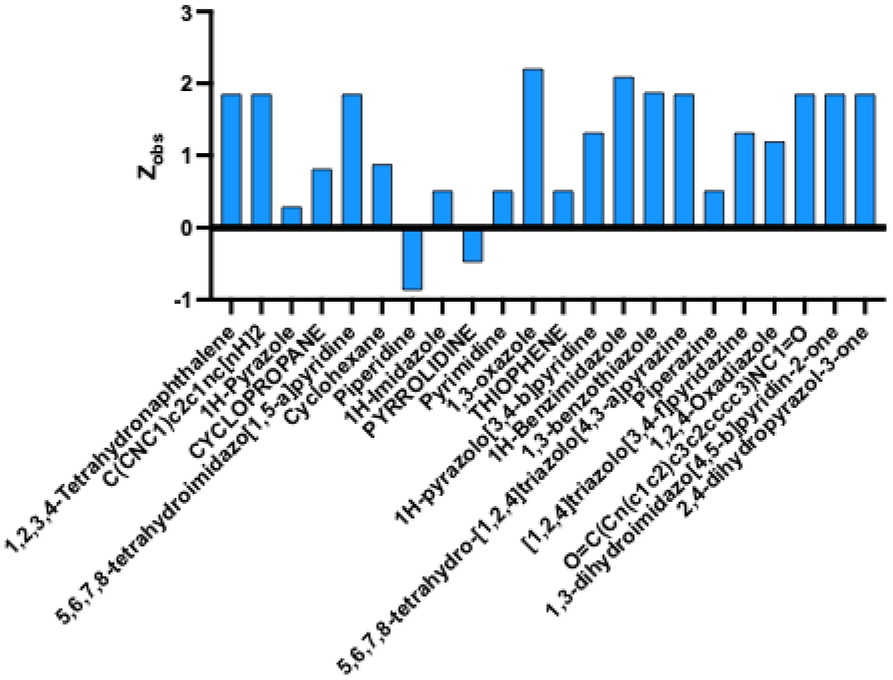


**Figure S2**. **Scaffold comparison between RNA-binding and non-binding fragments**. Core scaffolds for each fragment were identified using DataWarrior[4], and enrichment was quantified as Zobs based on scaffold occurrence and sample size in each pool.

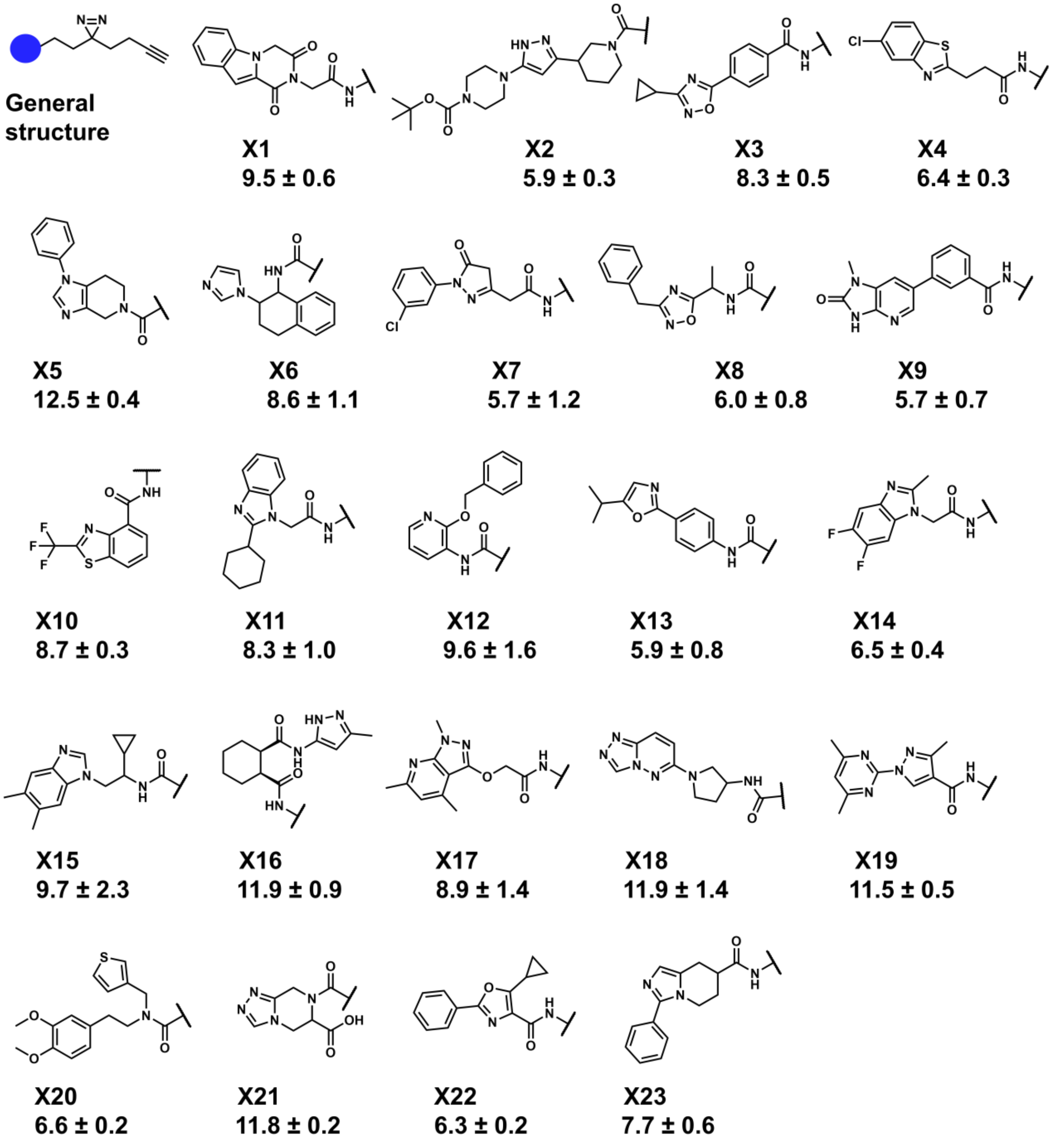


**Figure S3. Structures of hit small molecules that bind human transcripts from *in vitro* screening.** The values below the compound identifiers are the TAMRA signal normalized to the control probe (set as 1) from the *in vitro* screening assay, reported as average ± S.D. (n = 2 biological replicates).

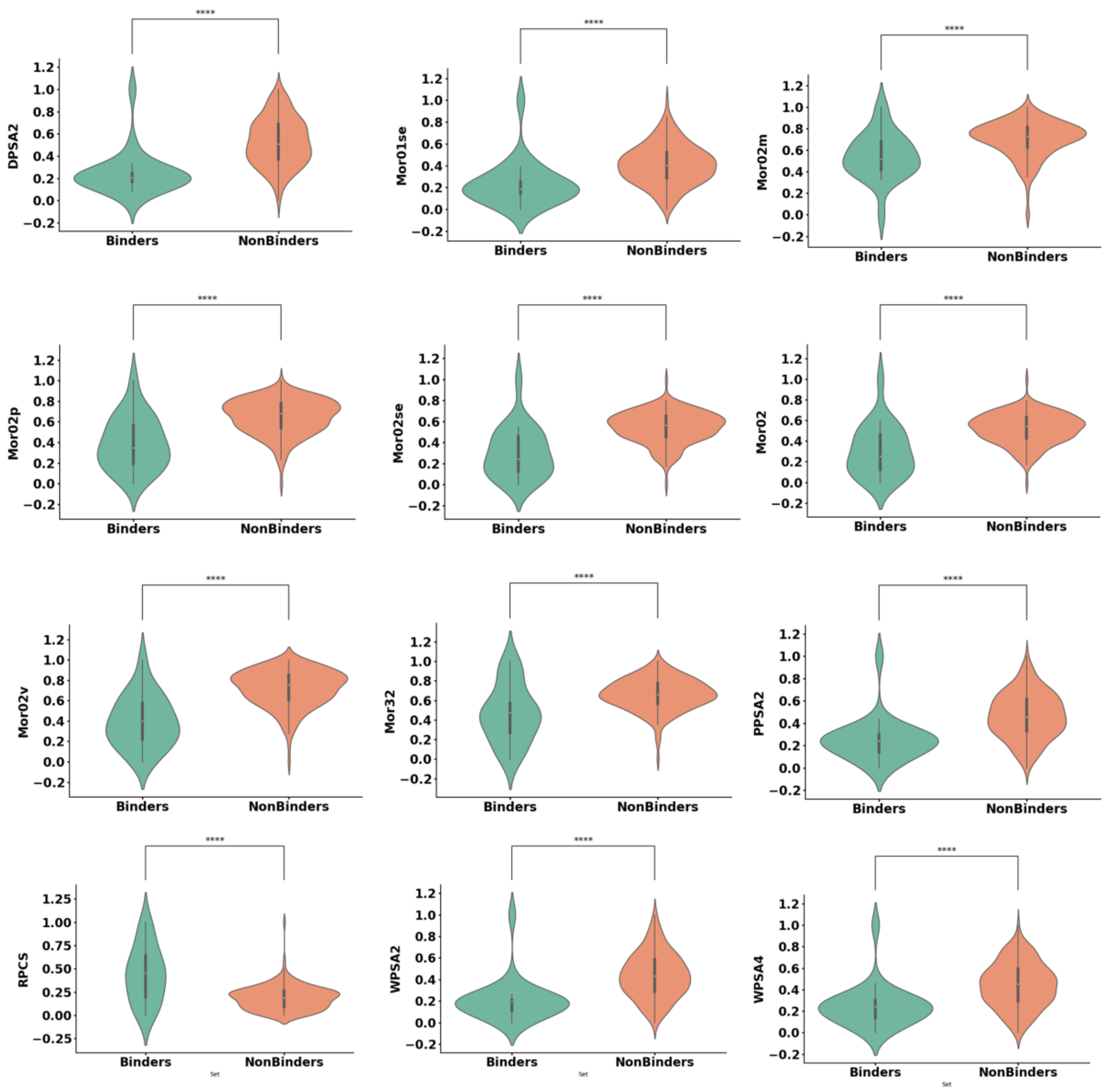


**Figure S4. Comparison of molecular descriptors that distinguish between RNA binders and non-binders.** A selection of 13 molecular descriptors—DPSA2, Mor01se, Mor02m, Mor02p, Mor02se, Mor02, Mor02v, Mor32, PPSA2, RPCS, WPSA2, WPSA4—with highest distinction between RNA-binding and non-binding fragments. A total of 1800 molecular descriptors were evaluated. These descriptors capture a range of physicochemical, geometrical, and surface area-based properties including partial charge distribution (PPSA/WPSA series), 3D molecular autocorrelation (Mor series), and relative polar surface characteristics (DPSA2, RPCS). Statistical analysis (Mann–Whitney) revealed that several descriptors show statistically significant differences between the two groups, suggesting distinct geometric and electrostatic profiles associated with RNA-binding activity. These features may serve as useful predictors in machine learning models for RNA-targeted ligand discovery.

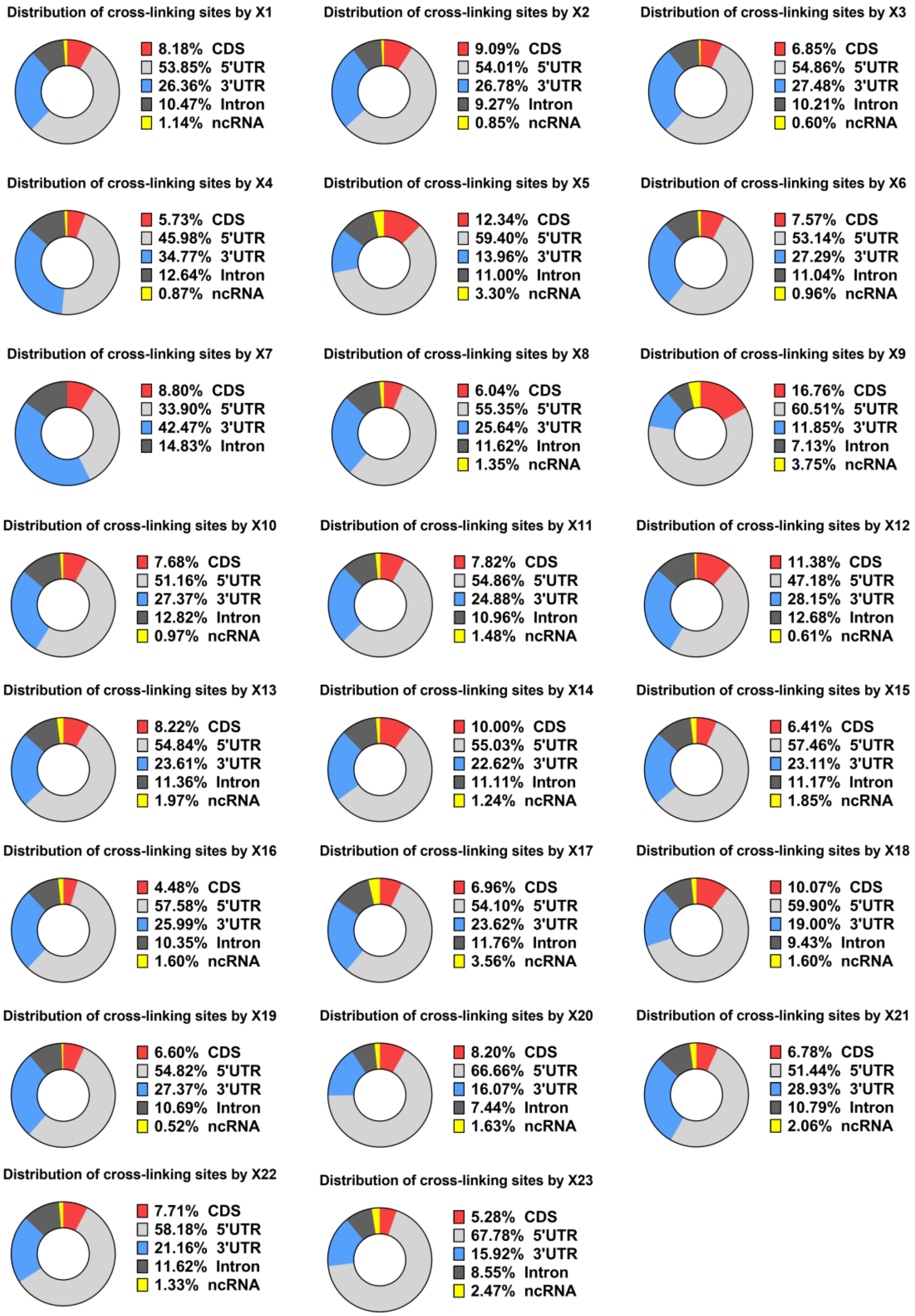

**Figure S5. Distribution of the location of cross-linked sites across the transcriptome.** The RT stop sites mapped by each small molecule to their bound RNA targets are categorized based on the types of RNA regions. CDS: coding sequences; UTR: untranslated regions; ncRNA: non-coding RNA.

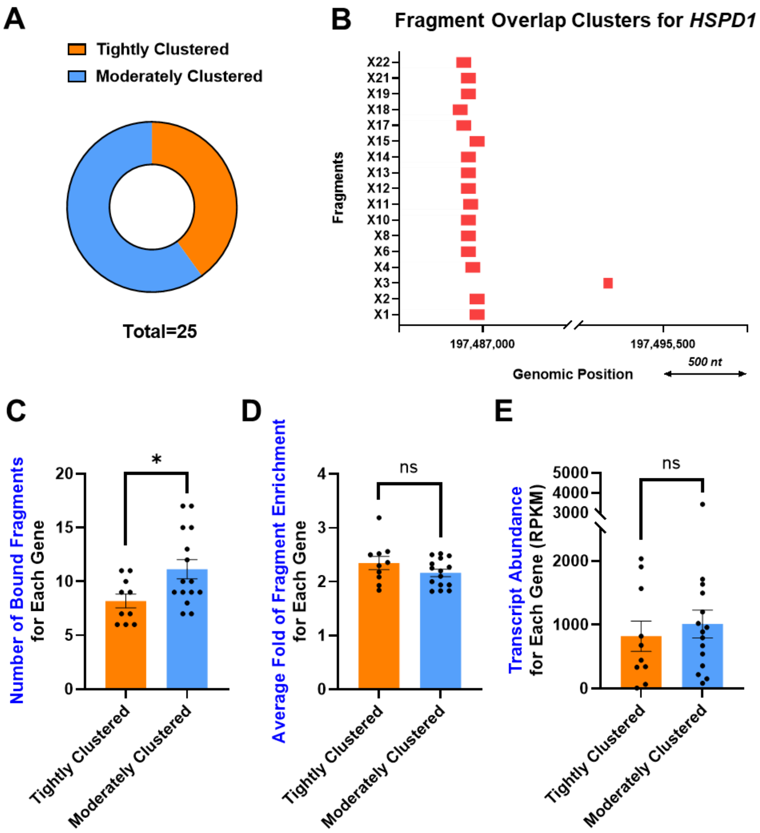


**Figure S6. Overlap analysis of compound binding regions for the top 25 genes, as ranked by the number of bound fragments.** **(A)** Distribution of genes with tightly or moderately clustered fragment binding regions. Tightly clustered (orange): for each bound fragment, there exists at least another fragment showing partially or fully overlapped binding regions with it. Moderately clustered (blue): there is at least one fragment showing unique binding region among all fragments bound to the same gene. **(B)** Exemplar plot showing genomic locations for clustered binding regions for the top 1 gene, HSPD1, which were enriched by 17 profiled fragments and moderately clustered. **(C)** The degree of binding region clustering is influenced by the number of bound fragments for each gene. **(D)** The average fold of fragment enrichment for each gene did not significantly differ between the tightly clustered gene and the moderately clustered genes. **(E)** Transcript abundance for each gene did not influence its degree of binding region clustering within the profiled dataset.

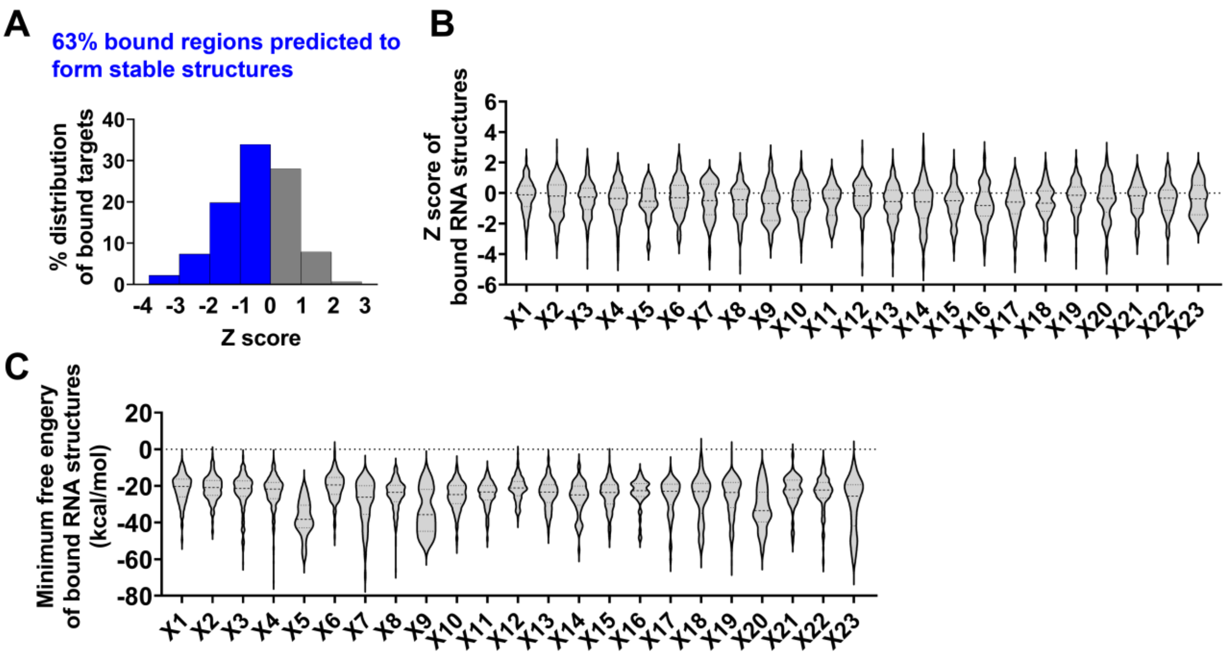


**Figure S7. Thermodynamic stability of RNA structures bound by small molecules.** (A) The distribution of z-scores of the bound RNA regions (window size = 120 nt) for 718 RNA targets enriched by small molecules (**X1** to **X23**). The z-scores were calculated by ScanFold[5] (window size = 120 nt; randomization = 100 sequences). (B) Violin plots showing the z-scores of RNA structures bound by each small molecule (**X1** – **X23**). (C) Violin plots showing the minimum free energy (MFE) of RNA structures bound by each small molecule (**X1** – **X23**).

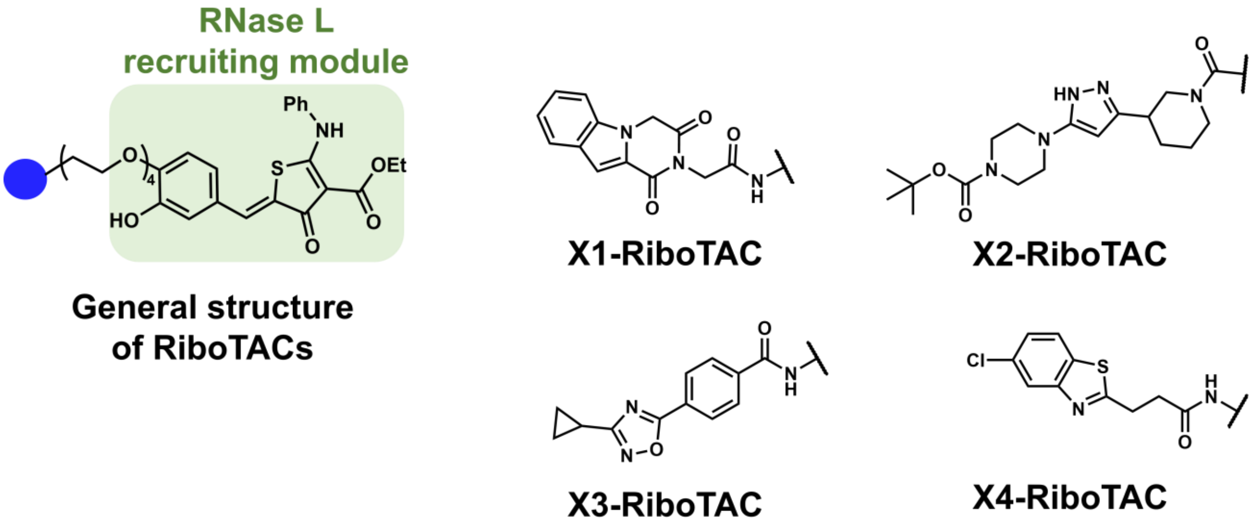


**Figure S8. Conversion of RNA binders into RiboTACs.** The general structure of RiboTACs and the structures of RiboTACs derived from the four prioritized RNA-binding small molecules (**X1**-**X4**).

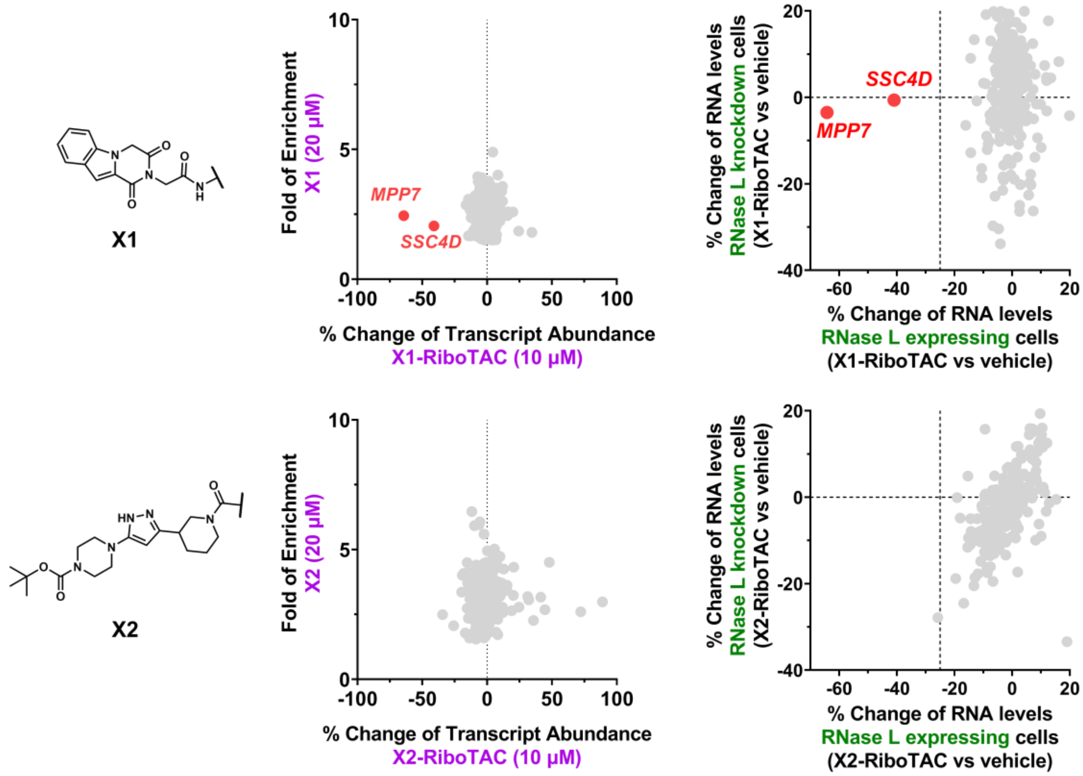

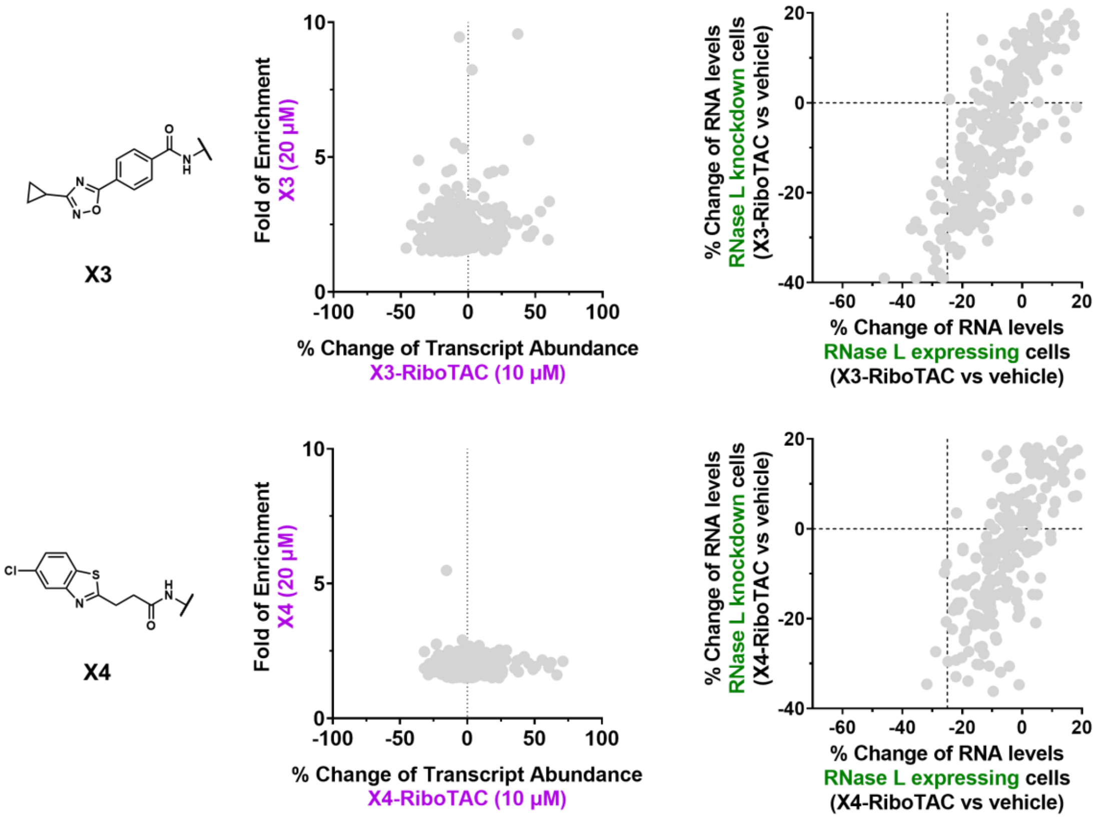


**Figure S9. Transcriptome-wide profiling of the four RiboTACs.** Left: structures of the RNA-binding modules of **X1** to **X4**. Middle: effects of RiboTACs on RNA targets bound by their corresponding FFFs. The x-axis shows the percent change in transcript abundances when treating the RiboTACs in CRISPR control MDA-MB-231 cells (express RNase L). The y-axis shows the fold enrichment for each target as identified by Chem-CLIP-Seq. Right: the x-axis shows the percent change in transcript abundances when treating the RiboTACs in CRISPR control MDA-MB-231 cells (express RNase L) while the y-axis shows the percent change in transcript abundances upon treatment of CRISPR modified MDA-MB-231 cells where RNase L is knocked down. These plots identify transcripts with RNase L-dependent reduction.

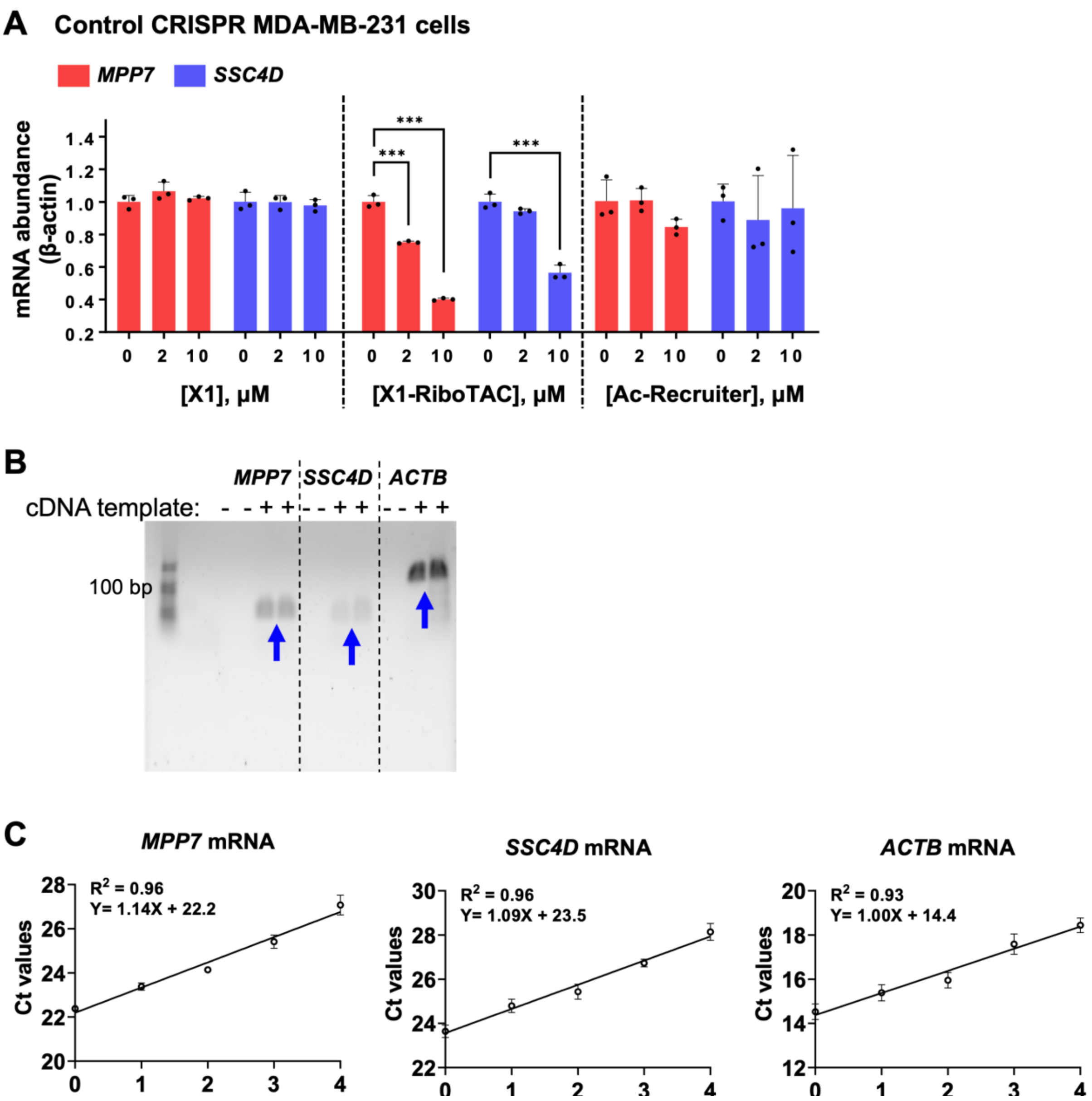


**Figure S10. Effect of X1 and X1-RiboTAC in control CRISPR MDA-MB-231 cells (express RNase L) and primer validation for RT-qPCR.** (A) Effect **X1**, **X1-RiboTAC**, and **Ac-Recruiter** in control CRISPR MDA-MB-231 cells (48 h treatment), as measured by RT-qPCR ($n$ = 3 biological replicates). (B) Representative gel image to assess the size and number of RT-qPCR products, where the blue arrows indicate the expected size ($n$ = 2 biological replicates). (C) Linear correlation between Ct values and the number of 1:2 cDNA dilutions used for RT-qPCR assay ($n$ = 3 biological replicates). *** $p < 0.001$, as determined by two-tailed Student's t-test. All data are reported as the mean ± SD.

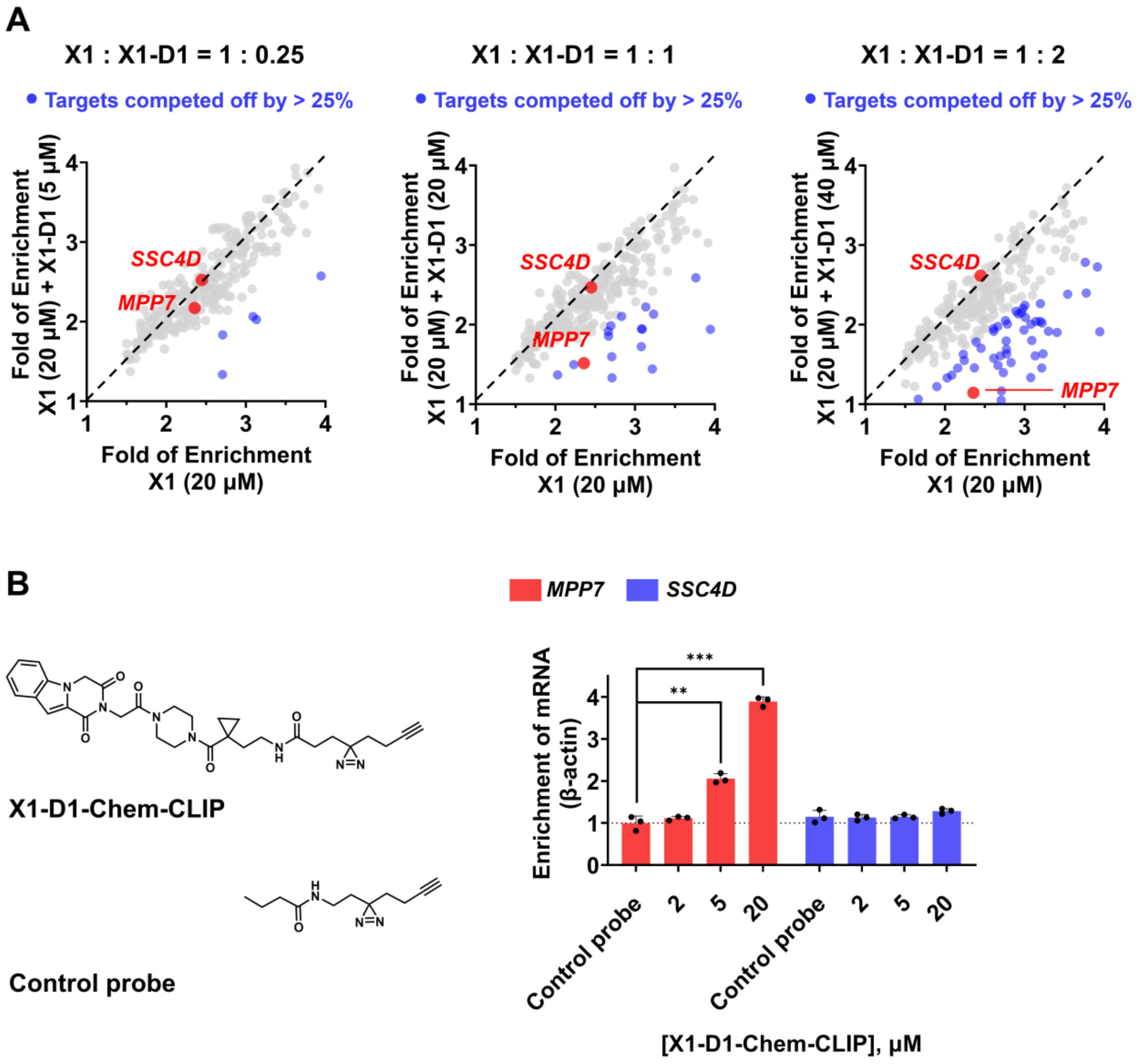


**Figure S11. Competitive Chem-CLIP between X1 and X1-D1 and direct target engagement by X1-D1-Chem-CLIP.** (A) RNA-seq profiling of competitive Chem-CLIP by co-treatment of **X1** (20 μM) with its derivative **X1-D1** at increasing concentrations (5 μM (1:0.25), 10 μM (1:1), 40 μM (1:2)) in WT MDA-MB-231 cells. The x-axis shows the fold enrichment for 282 RNA targets enriched by **X1** in the absence competing **X1-D1** (16 h treatment, 20 μM). The y-axis shows the fold enrichment for the same 282 RNA targets bound by **X1** when co-treated with the competitor, **X1-D1**, at different concentrations. Each dot represents one RNA target, and the blue dots represent the targets whose enrichment was reduced by >25% when co-treated with **X1-D1**. (B) Direct target engagement (enrichment) of *MPP7* mRNA as a function of **X1-D1-Chem-CLIP** concentration in WT MDA-MB-231 cells (16 h treatment, 20 μM, *n* = 3 biological replicates), as measured by RT-qPCR.  ** $p < 0.01$, *** $p < 0.001$, as determined by two-tailed Student's t-test.  All data are reported as the mean ± SD.

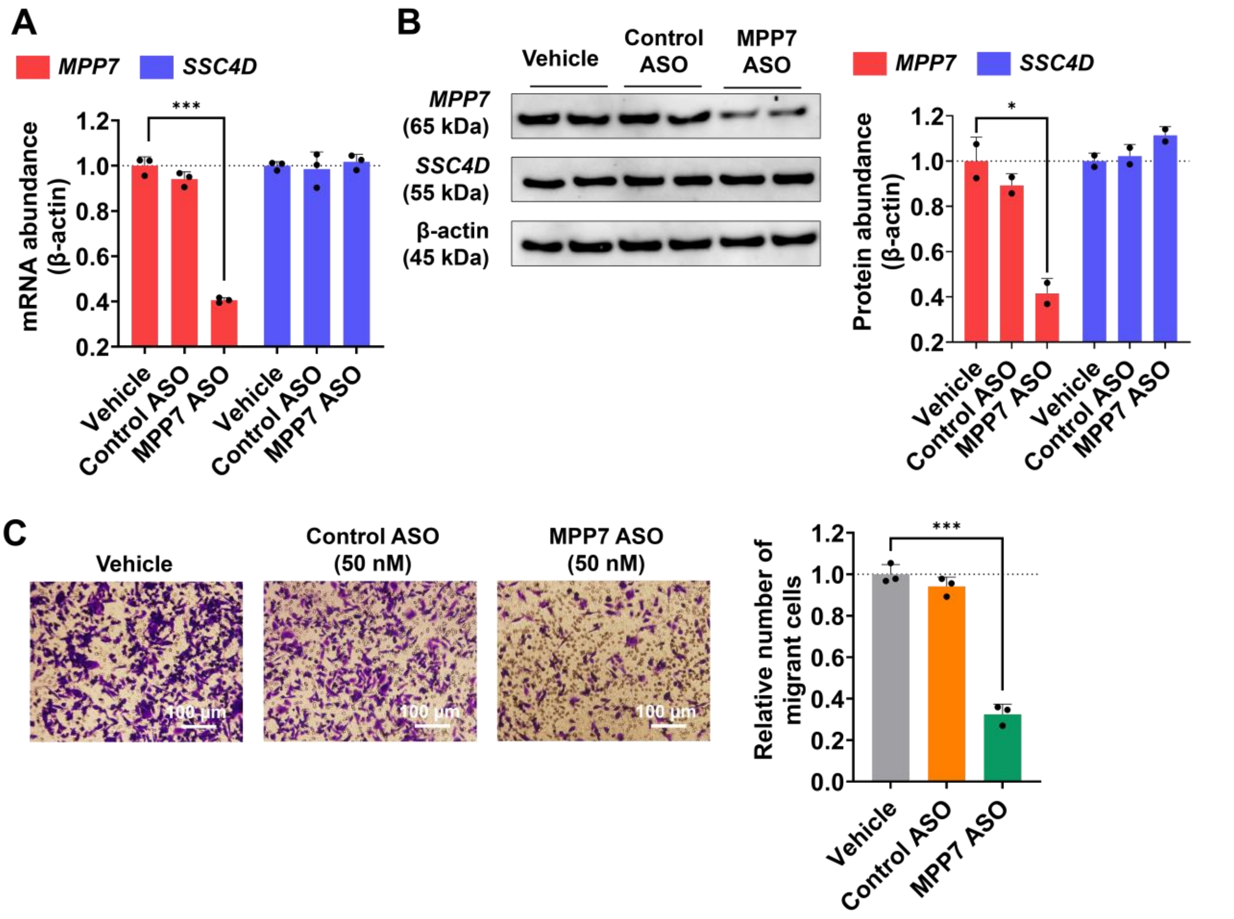


**Figure S12. Antisense oligonucleotide[6] targeting *MPP7* inhibits the migration of wild-type MDA-MB-231 cells.** (A) Effect of an *MPP7*-targeted ASO and a non-targeting control ASO on the abundance *MPP7* and *SSC4D* transcripts in WT MDA-MB-231 cells after 48 h treatment, as measured by RT-qPCR ($n$ = 3 biological replicates). (B) Effect of an *MPP7*-targeted ASO and a non-targeting control ASO on MPP7 and SSC4D protein levels in WT MDA-MB-231 cells after 48 h treatment, as measured by Western blotting ($n$ = 2 biological replicates). (C) Effect of an *MPP7*-targeted ASO and a non-targeting control ASO on the number of migratory MDA-MB-231 cells, as measured by the Boyden chamber assay ($n$ = 3 biological replicates). * $p < 0.05$, *** $p < 0.001$, as determined by two-tailed Student's t-test. All data are reported as the mean ± SD.

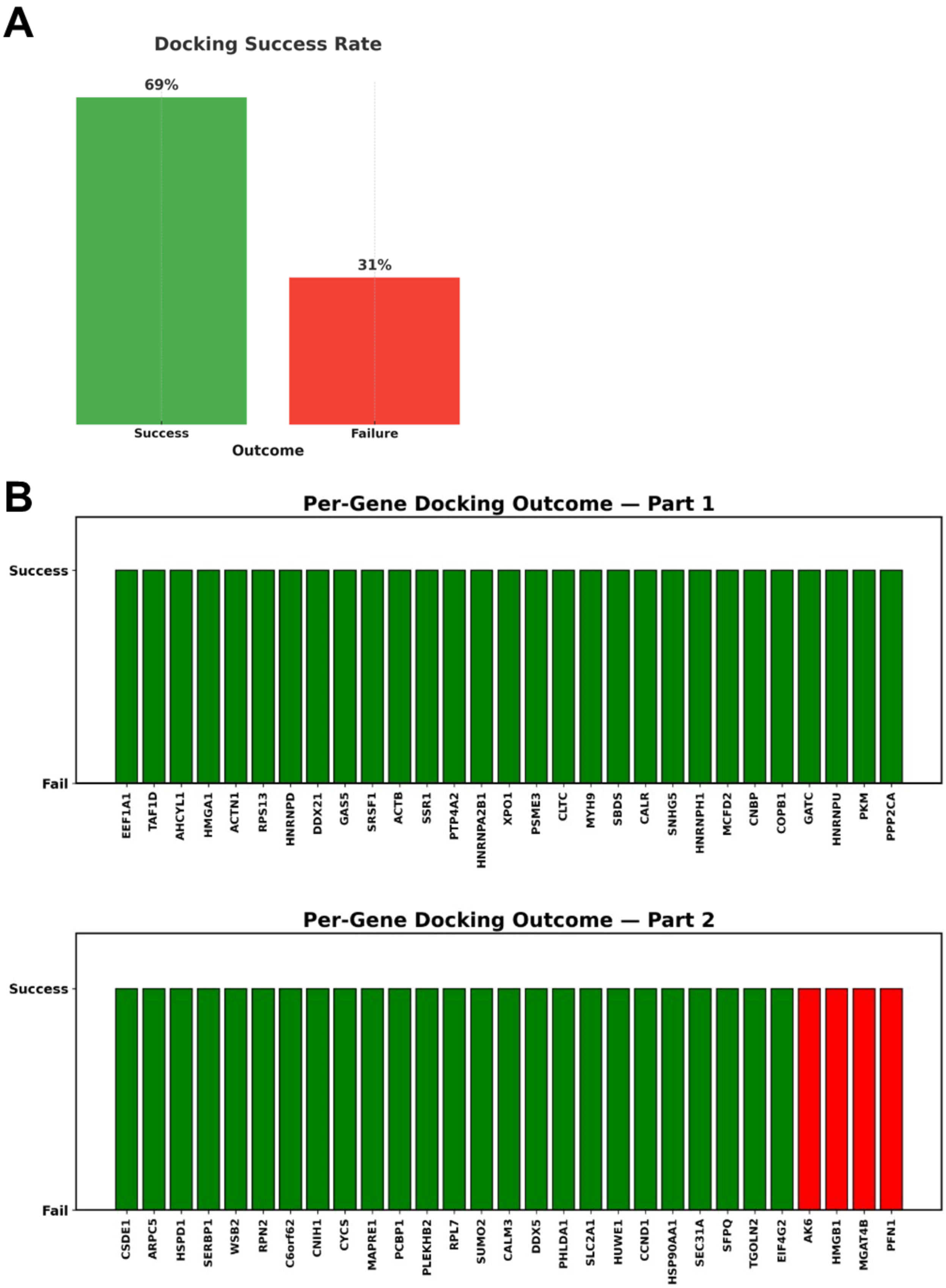
A
Docking Success Rate
69%
31%
Success
Failure
Outcome
B
Per-Gene Docking Outcome — Part 1
Success
Fail
EEF1A1
TAF1D
AHCYL1
HMGA1
ACTN1
RPS13
HNRNPD
DDX21
GAS5
SRSF1
ACTB
SSR1
PTP4A2
HNRNPA2B1
XPO1
PSME3
CLTC
MYH9
SBDS
CALR
SNHG5
HNRNPH1
MCFD2
CNBP
COPB1
GATC
HNRNPU
PKM
PPP2CA
Per-Gene Docking Outcome — Part 2
Success
Fail
CSDE1
ARPC5
HSPD1
SERBP1
WSB2
RPN2
C6orf62
CNIH1
CYCS
MAPRE1
PCBP1
PLEKHB2
RPL7
SUMO2
CALM3
DDX5
PHLDA1
SLC2A1
HUWE1
CCND1
HSP90AA1
SEC31A
SFPQ
TGOLN2
EIF4G2
AK6
HMGB1
MGAT4B
PFN1

**Figure S13. Success rate of RNA-ligand docking poses mapped to functional binding sites.** (A) The bar chart summarizes the number of successful (green) and failed (red) docking poses based on their spatial proximity to experimentally defined RNA sites. A docking pose was classified as successful if the center of mass[7] of the ligand was within 10 Å of the COM of the target RNA residue. (B) analysis of the first three lowest energy docked poses per each gene targeted by **X1** shows for all of the genes except for AK6,HMGB1,MGAT4B and PFN1 there is one pose correctly positioning the **X1** close to the FFF site.

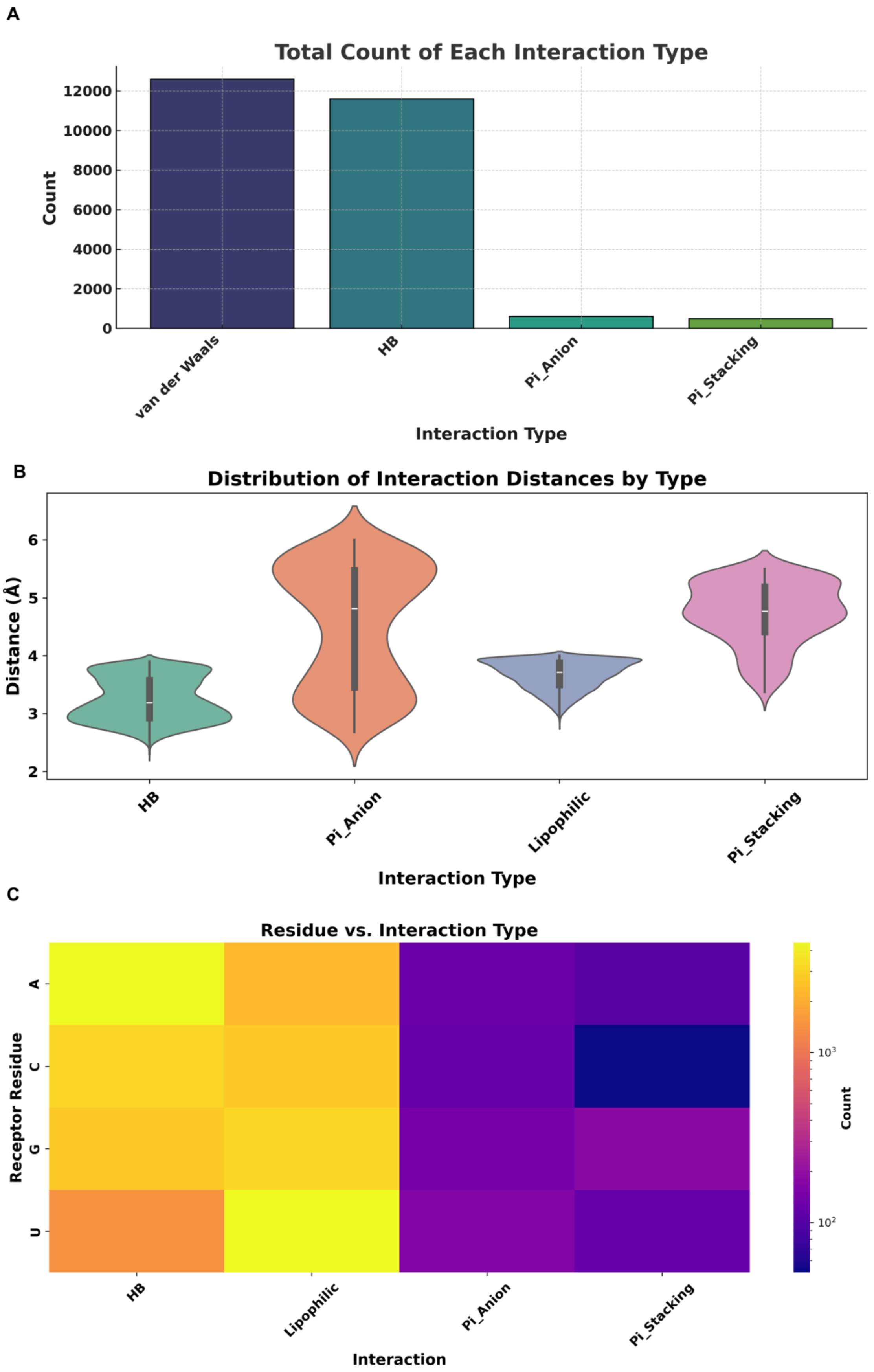
A
Total Count of Each Interaction Type
Count
12000
10000
8000
6000
4000
2000
0
van der Waals
HB
Pi_Anion
Pi_Stacking
Interaction Type
B
Distribution of Interaction Distances by Type
Distance (Å)
6
5
4
3
2
HB
Pi_Anion
Lipophilic
Pi_Stacking
Interaction Type
C
Residue vs. Interaction Type
Receptor Residue
A
C
G
U
HB
Lipophilic
Pi_Anion
Pi_Stacking
Interaction
Count
10^3
10^2

**Figure S14. Summary of ligand–RNA interaction types, distances, and residue preferences.** (A) The bar plot shows the total count of each interaction type observed between small molecules and RNA targets. Lipophilic and hydrogen bonding (HB) interactions are the most prevalent, each occurring over 11,000 times, whereas π-anion and π-stacking interactions are comparatively rare. (B) The violin plot depicts the distribution of interaction distances for each interaction type. Hydrogen bonds tend to occur at shorter distances (~2.5–3.5 Å), consistent with canonical bonding geometries. Lipophilic interactions also occur at short ranges (~3.5 Å), while π-anion and π-stacking interactions show broader distributions and slightly longer typical distances (~4.5–5.5 Å), reflecting the delocalized nature of these interactions. (C) The heatmap illustrates the frequency of each interaction type per receptor RNA residue (A, C, G, U). Adenine (A) and guanine (G) participate more frequently in hydrogen bonding and lipophilic interactions, while uracil (U) shows relatively lower overall interaction counts. Notably, cytosine (C) has fewer π-anion interactions compared to other nucleobases.

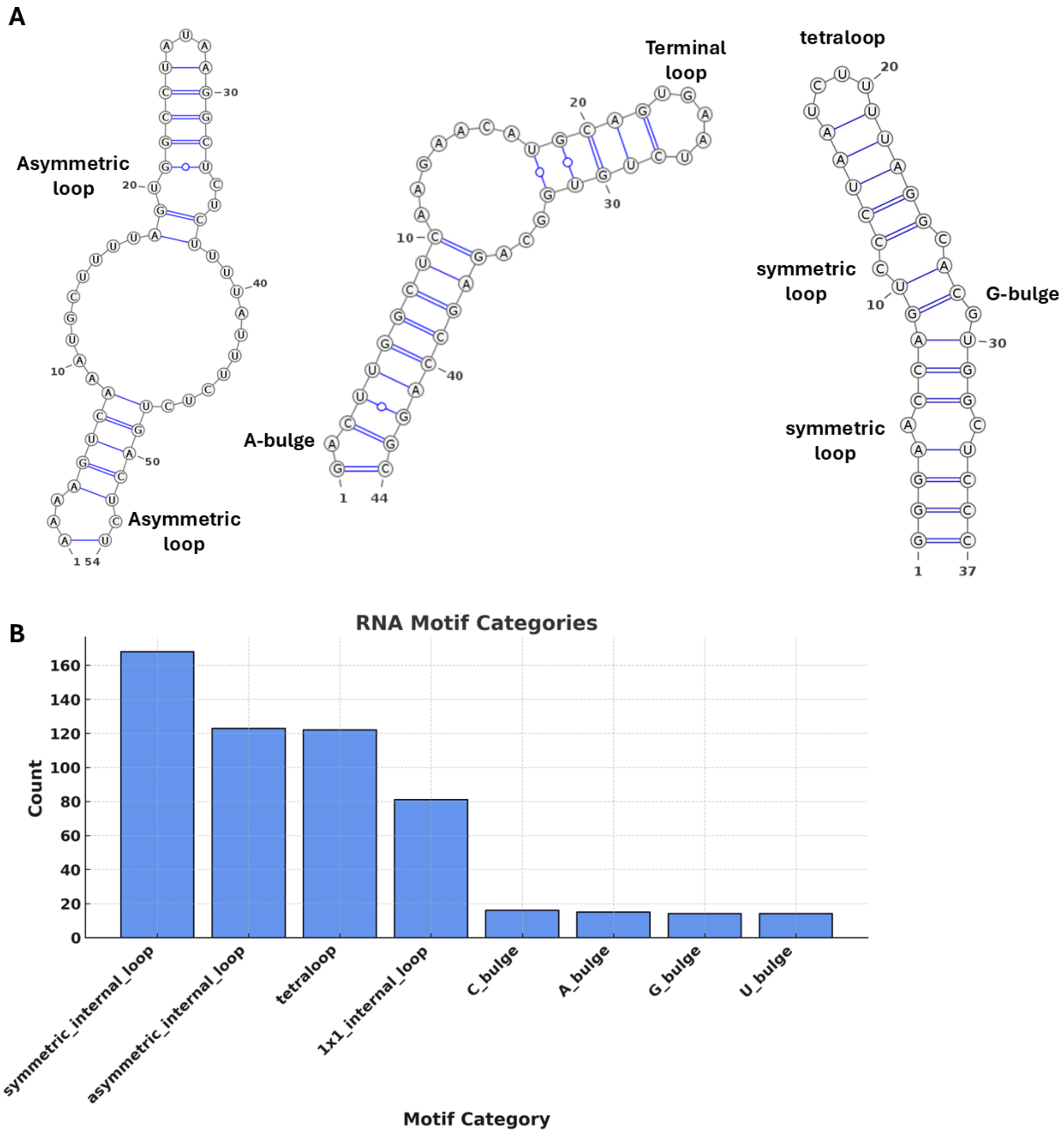


**Figure S15. Secondary structure categorization of short, medium and long binding sites**. Thermodynamically most stable secondary structure representatives bound by **X1** in short (<30 nt), medium (30<Y<60 nt), and long (>65 nt). The name of each gene is written under the secondary structure. Y stands for the RNA chain length. (B) Bar plot showing the frequency of distinct RNA motif categories observed at ligand interaction sites. Symmetric internal loops were the most common motif type, followed by asymmetric internal loops, GNRA tetraloops, and 1×1 internal loops, each comprising a substantial fraction of the dataset. In contrast, single-nucleotide bulges (C, A, G, and U) were markedly less frequent, each occurring fewer than 20 times.

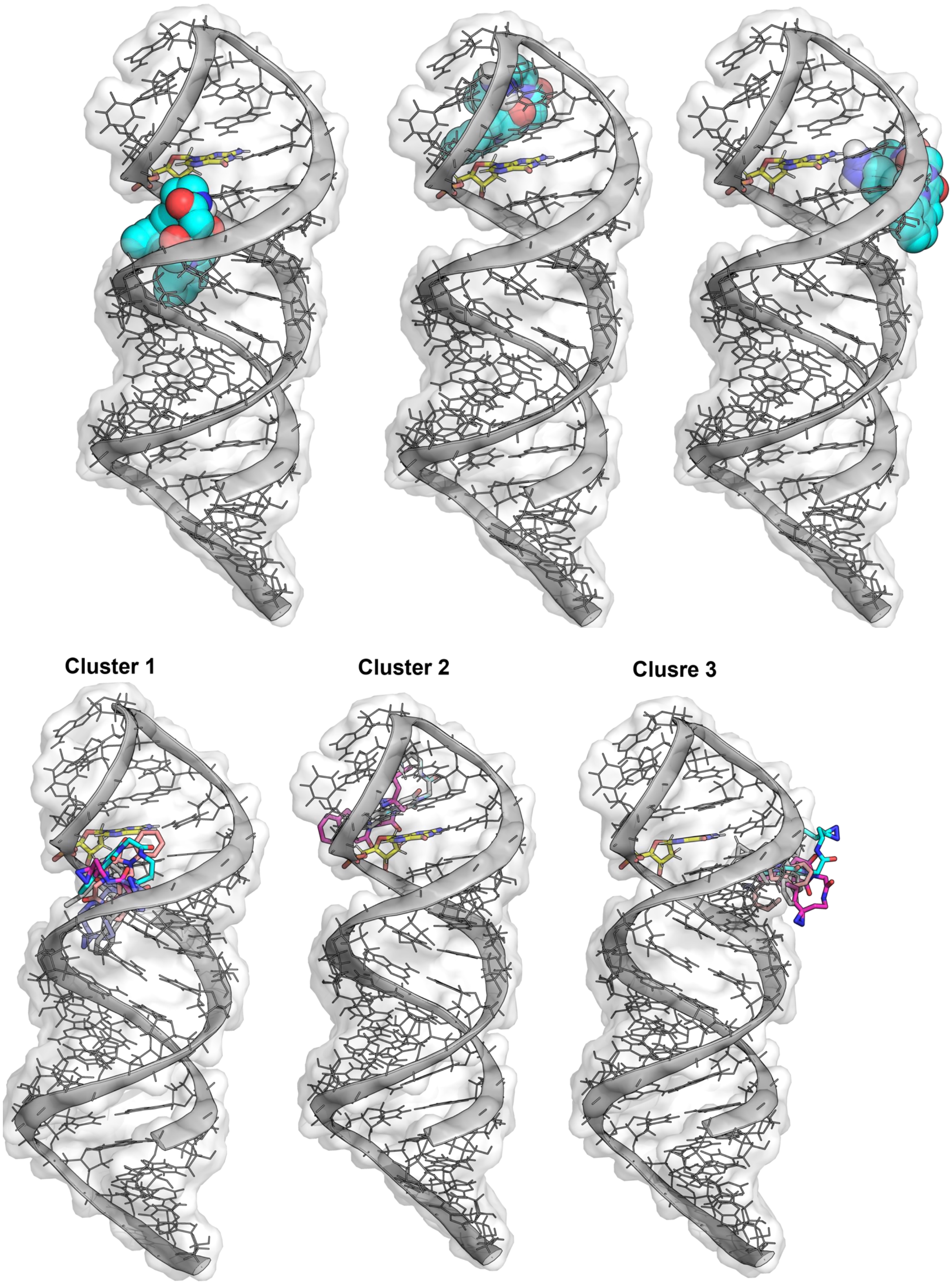
Cluster 1
Cluster 2
Clusre 3

**Figure S17. First three lowest energy docked poses of X1 against the MPP7 mRNA.** The RNA is shown in cartoon, surface and lines. The cross-linking site is highlighted in yellow (sticks). Cluster analysis of the first 10 poses of X1 docked against the MPP7. The first three lowest energy poses (from right to left respectively) were used to cluster the similar poses.

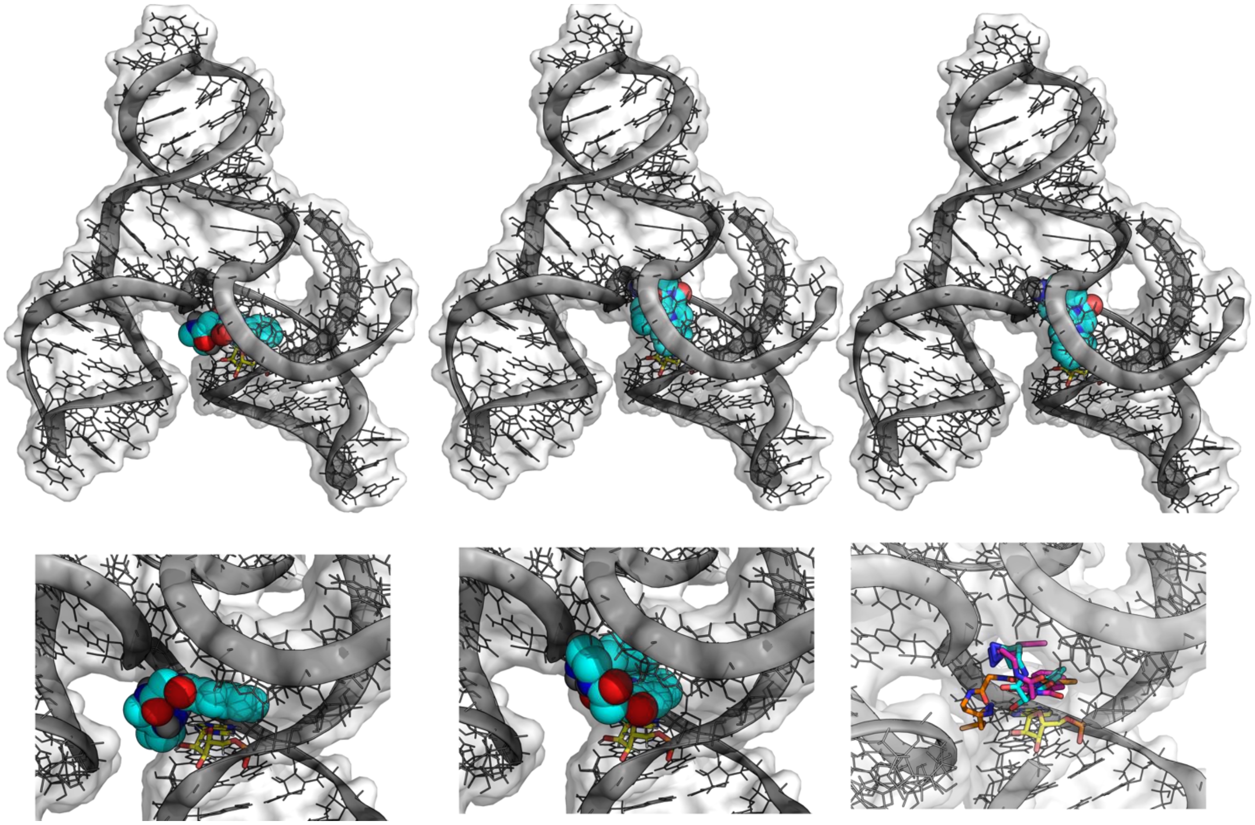

**Figure S18. Docking poses of the first three lowest energy states of X1 against the SSCD4.** The RNA is shown in cartoon, surface and lines. The cross-linking site is highlighted in yellow (sticks).

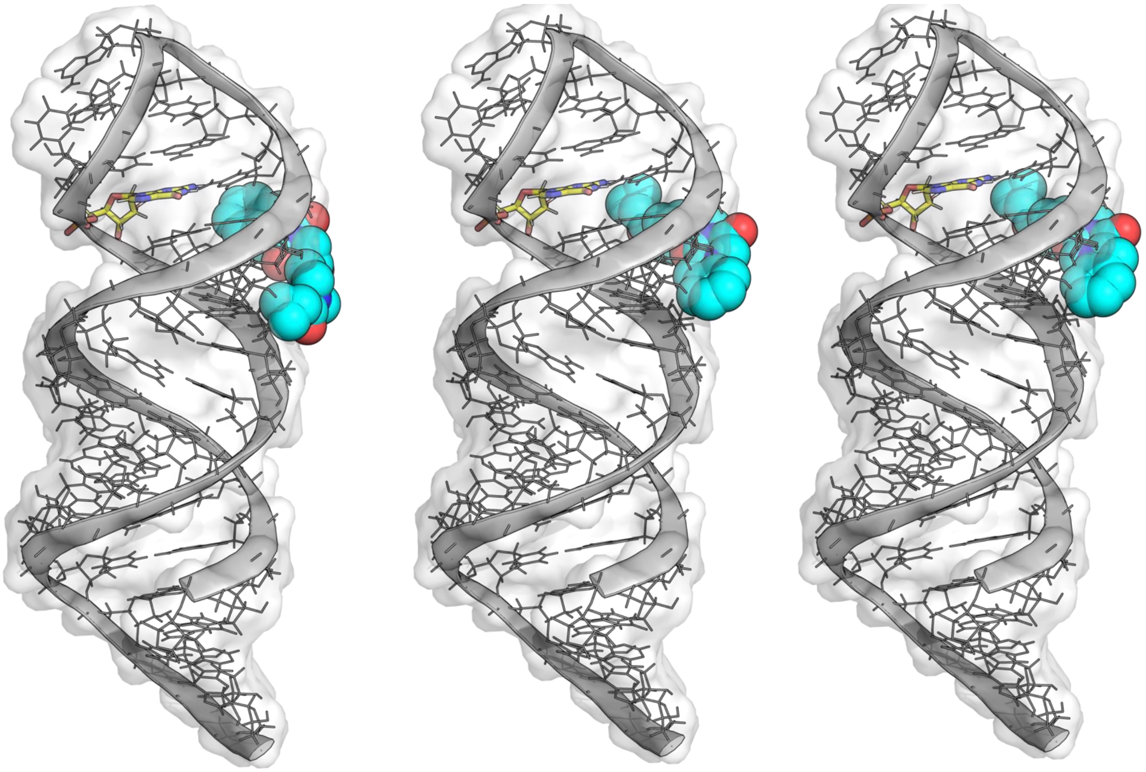

**Figure S19. First three lowest energy docked poses of X1-D1 against the MPP7 mRNA.** The RNA is shown in cartoon, surface and lines. The cross-linking site is highlighted in yellow (sticks).

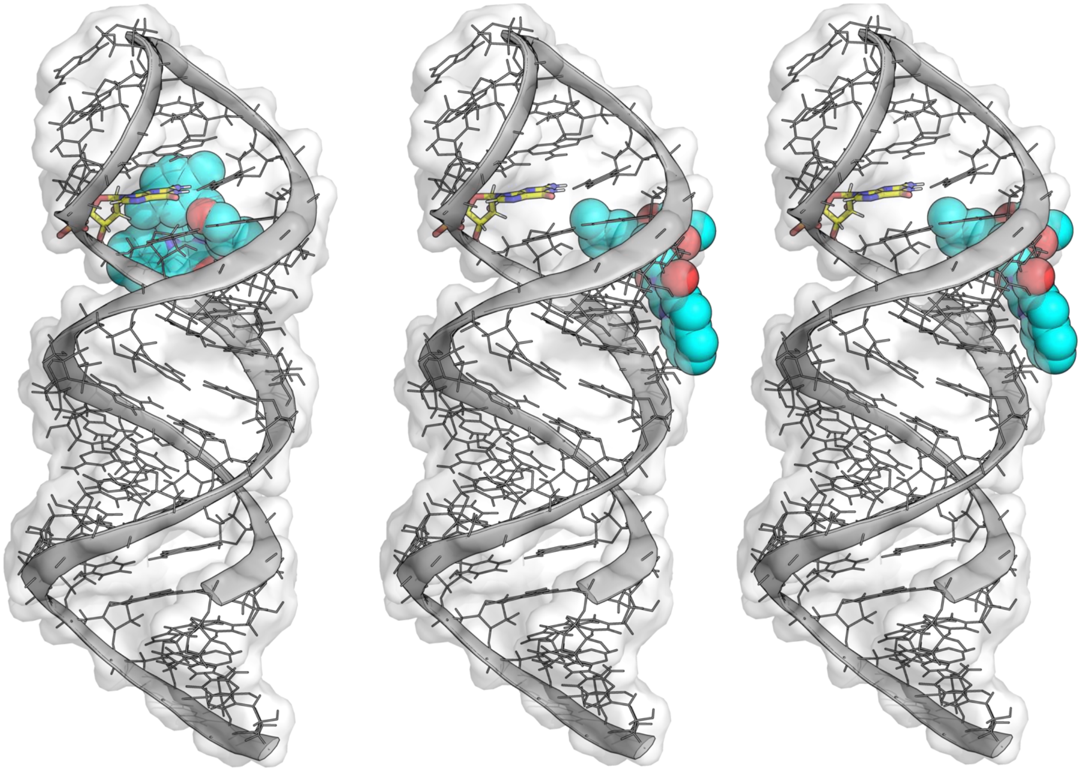

**Figure S20. First three lowest energy docked poses of X1-D2 against the MPP7 mRNA.** The RNA is shown in cartoon, surface and lines. The cross-linking site is highlighted in yellow (sticks).

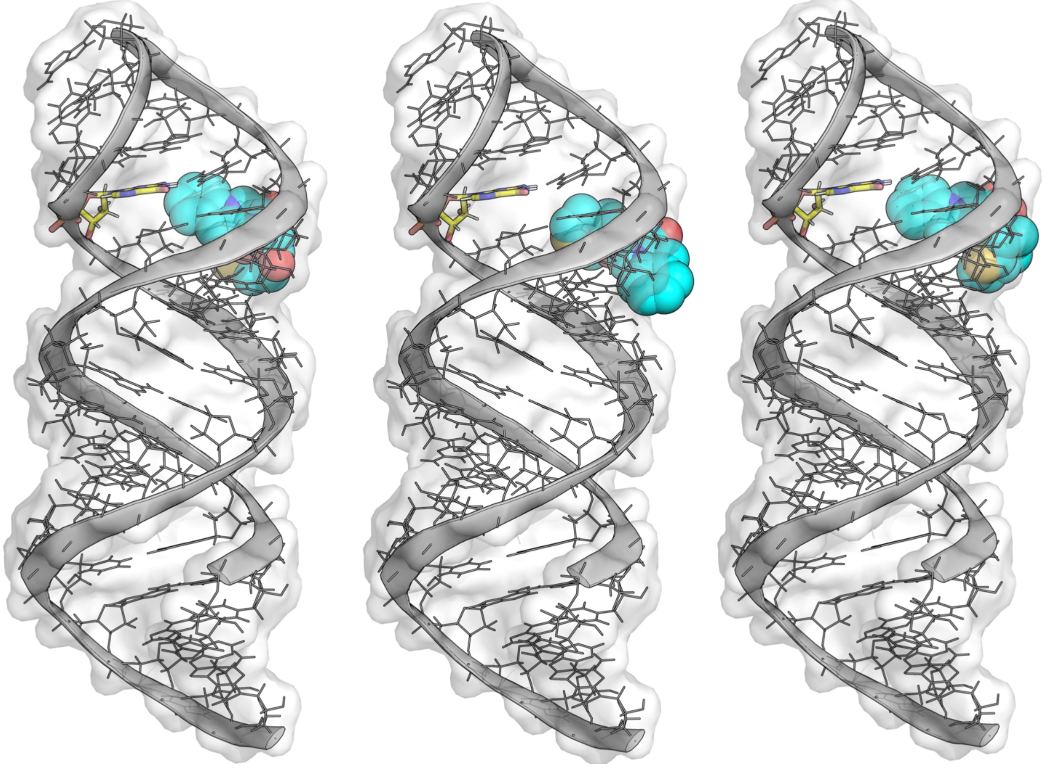

**Figure S21. First three lowest energy docked poses of X1-D3 against the MPP7 mRNA.** The RNA is shown in cartoon, surface and lines. The cross-linking site is highlighted in yellow (sticks).

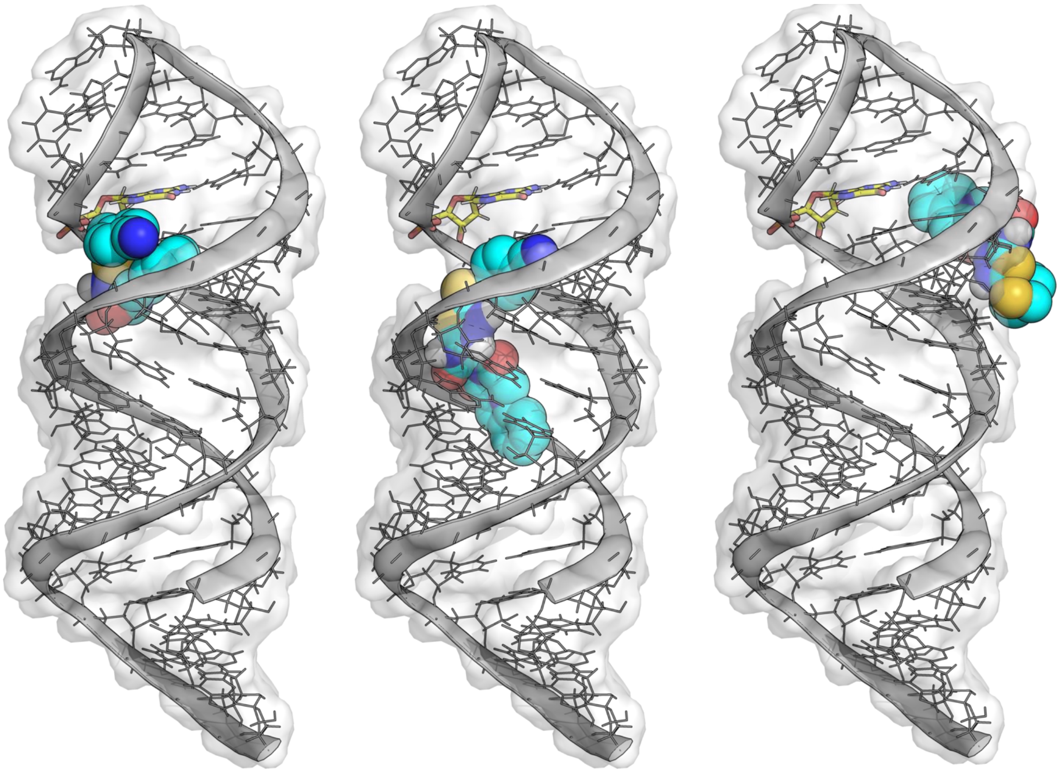

**Figure S22. First three lowest energy docked poses of X1-D4 against the MPP7 mRNA.** The RNA is shown in cartoon, surface and lines. The cross-linking site is highlighted in yellow (sticks).

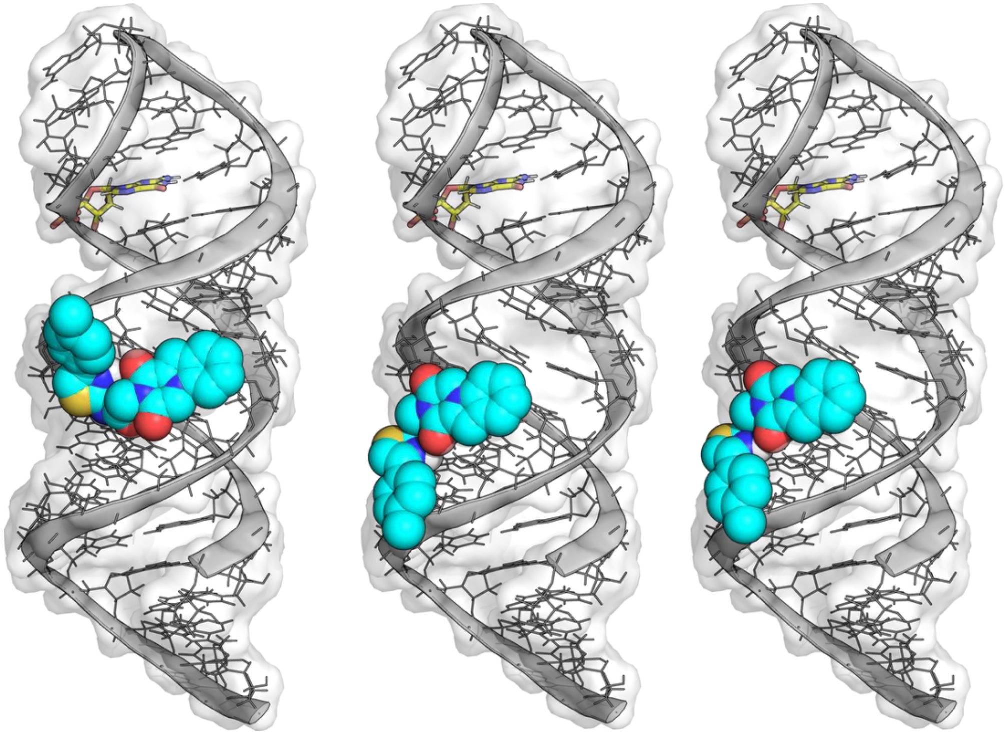

**Figure S23. First three lowest energy docked poses of X1-D5 against the MPP7 mRNA.** The RNA is shown in cartoon, surface and lines. The cross-linking site is highlighted in yellow (sticks).

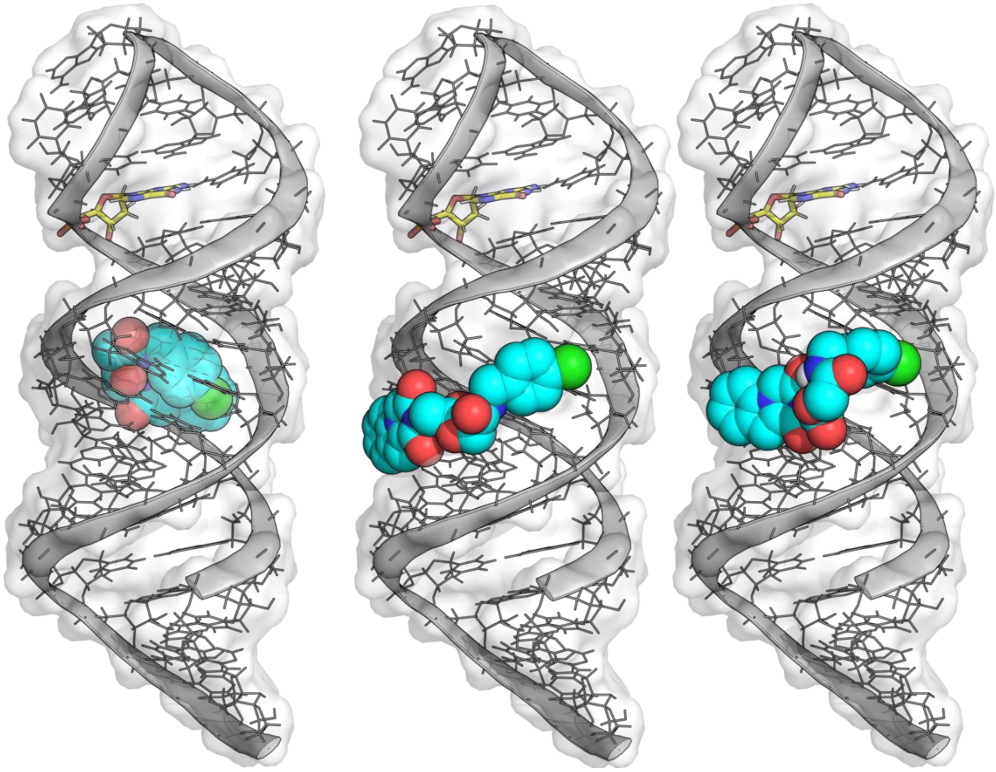

**Figure S24. First three lowest energy docked poses of X1-D6 against the MPP7 mRNA.** The RNA is shown in cartoon, surface and lines. The cross-linking site is highlighted in yellow (sticks).

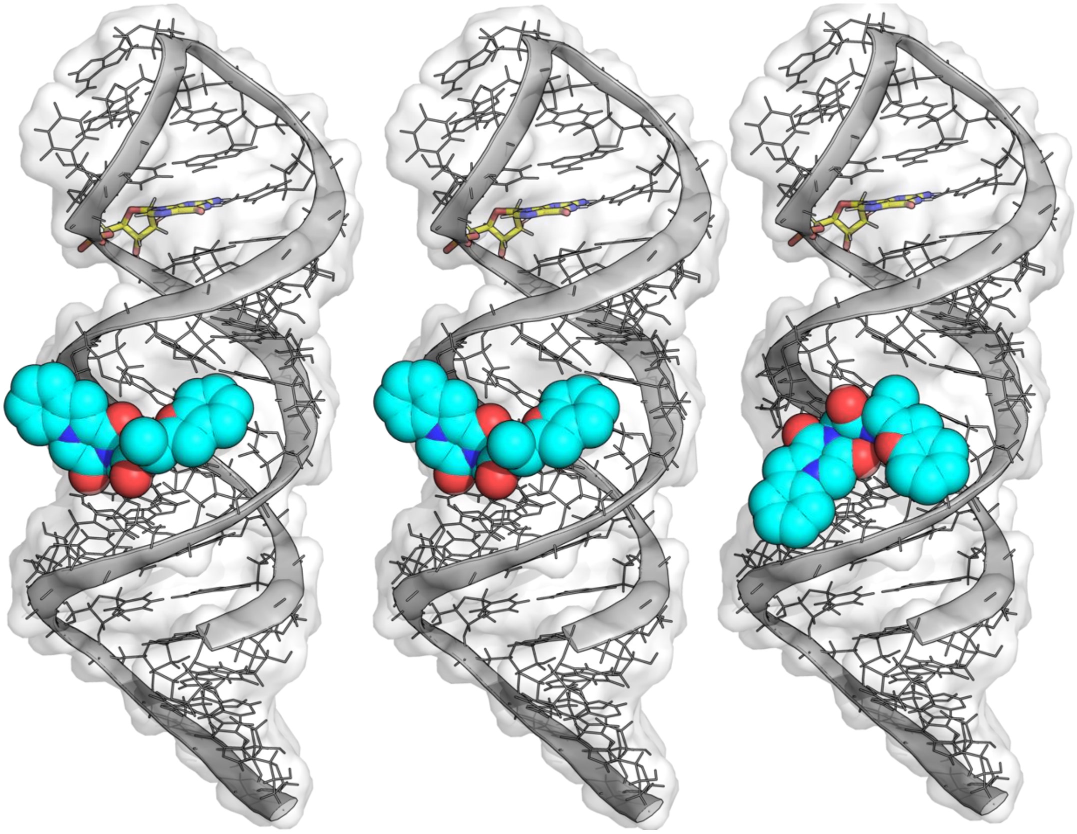

**Figure S25. First three lowest energy docked poses of X1-D7 against the MPP7 mRNA.** The RNA is shown in cartoon, surface and lines. The cross-linking site is highlighted in yellow (sticks).

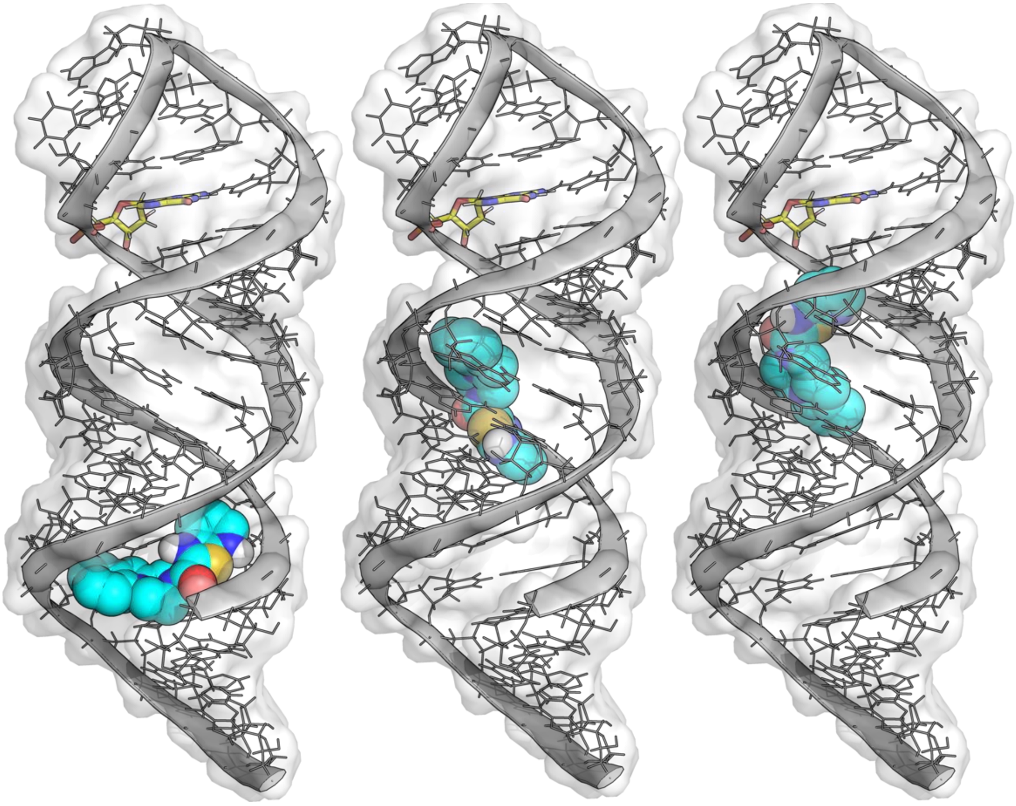

**Figure S26. First three lowest energy docked poses of X1-D8 against the MPP7 mRNA.** The RNA is shown in cartoon, surface and lines. The cross-linking site is highlighted in yellow (sticks).

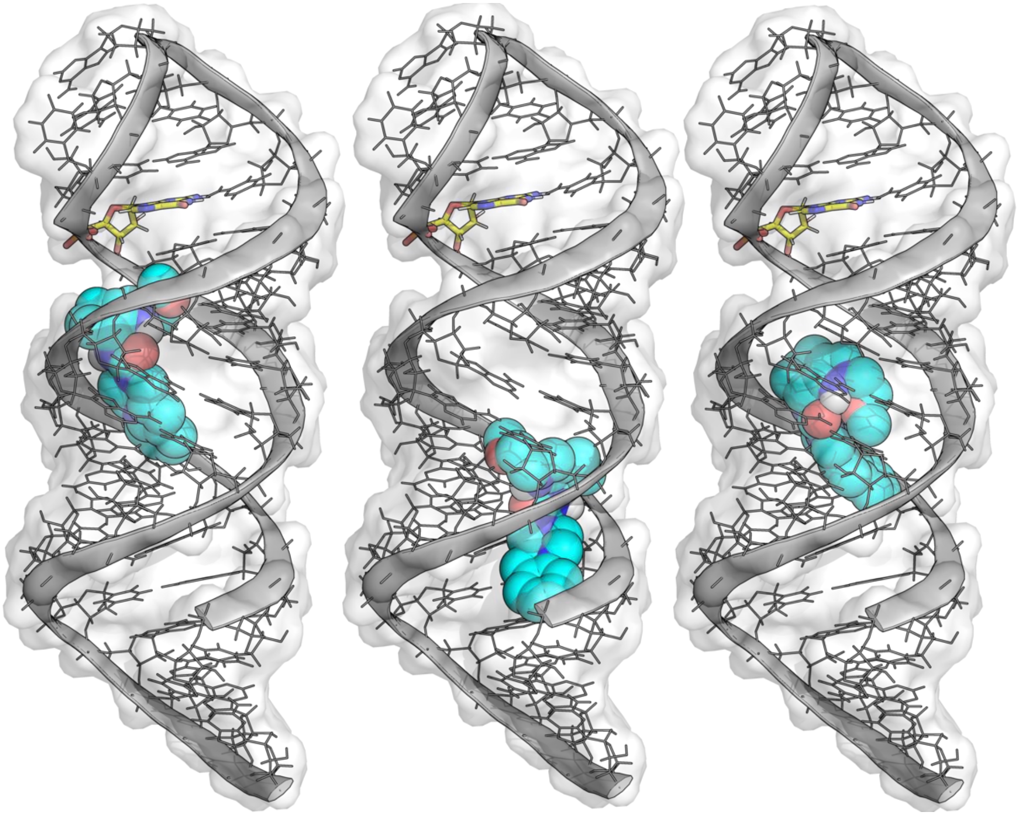

**Figure S27. First three lowest energy docked poses of X1-D9 against the MPP7 mRNA.** The RNA is shown in cartoon, surface and lines. The cross-linking site is highlighted in yellow (sticks).

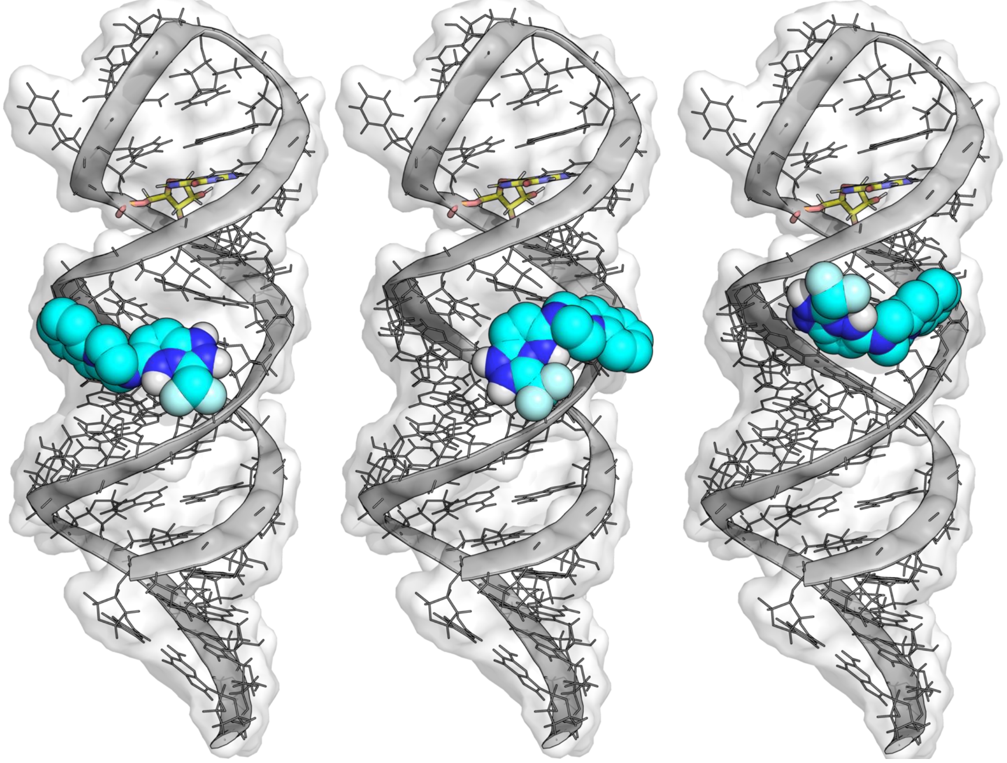

**Figure S28. First three lowest energy docked poses of X1-D10 against the MPP7 mRNA.** The RNA is shown in cartoon, surface and lines. The cross-linking site is highlighted in yellow (sticks).

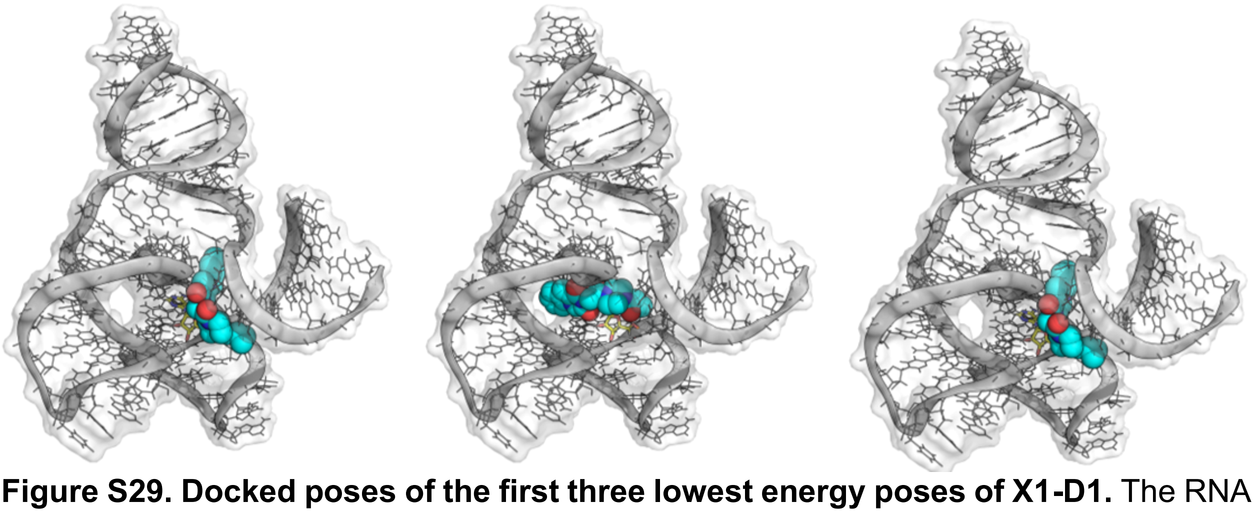

**Figure S29. Docked poses of the first three lowest energy poses of X1-D1.** The RNA is shown in cartoon, surface and lines. The cross-linking site is highlighted in yellow (sticks).

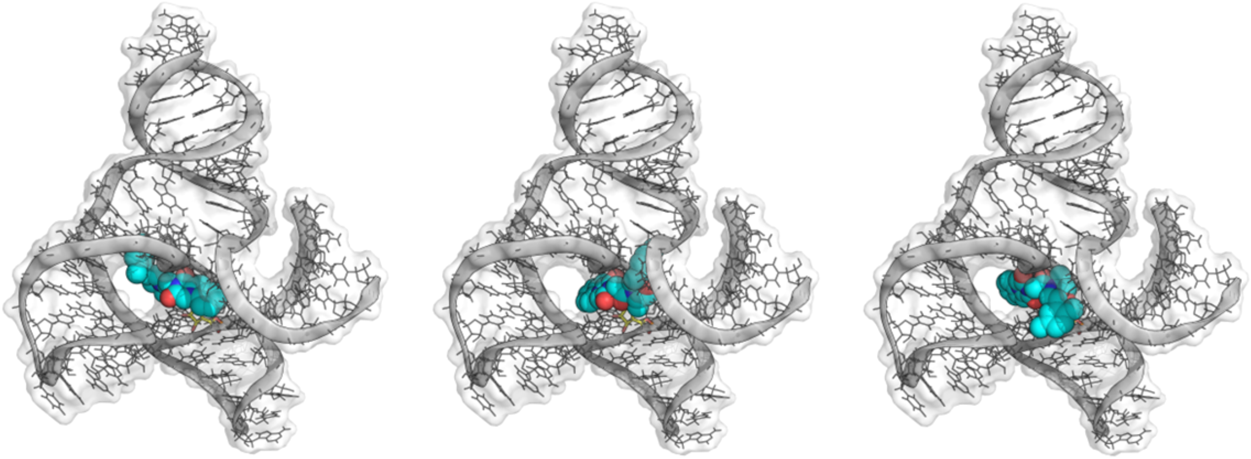

**Figure S30. Docked poses of the first three lowest energy poses of X1-D2.** The RNA is shown in cartoon, surface and lines. The cross-linking site is highlighted in yellow (sticks).

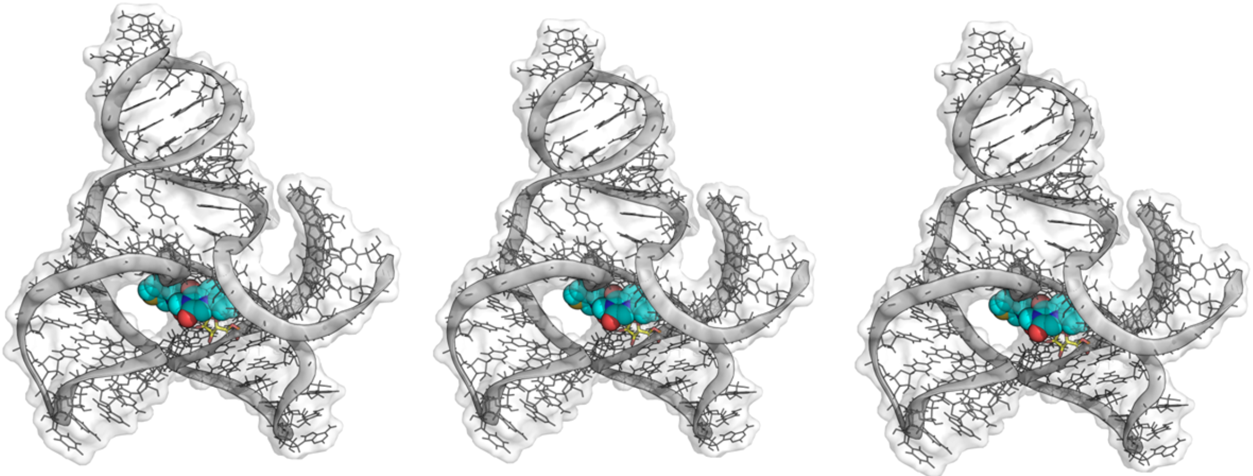

**Figure S31. Docked poses of the first three lowest energy poses of X1-D3.** The RNA is shown in cartoon, surface and lines. The cross-linking site is highlighted in yellow (sticks).

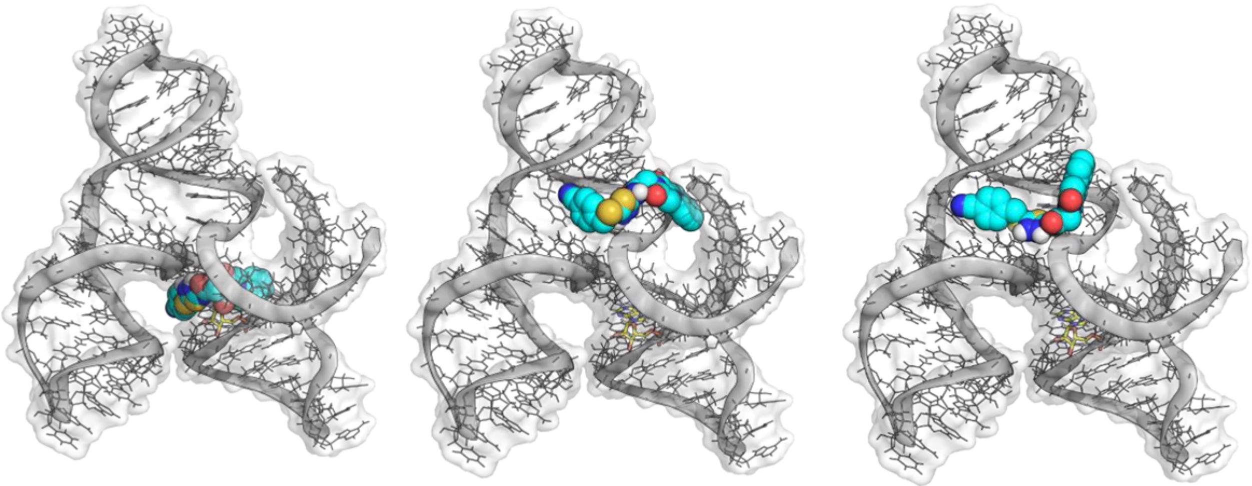

**Figure S32. Docked poses of the first three lowest energy poses of X1-D4.** The RNA is shown in cartoon, surface and lines. The cross-linking site is highlighted in yellow (sticks).

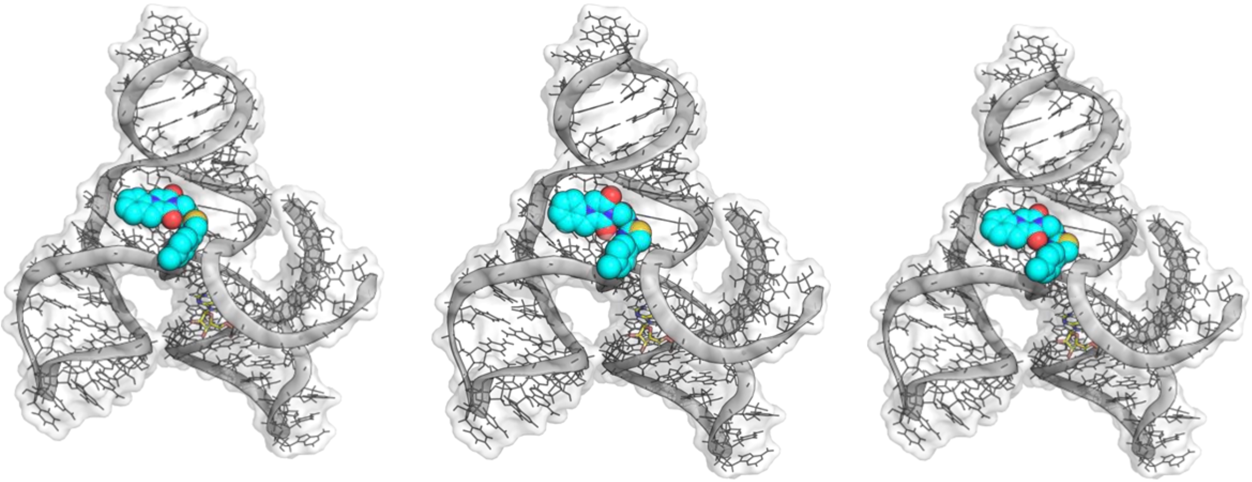

**Figure S33. Docked poses of the first three lowest energy poses of X1-D5.** The RNA is shown in cartoon, surface and lines. The cross-linking site is highlighted in yellow (sticks).

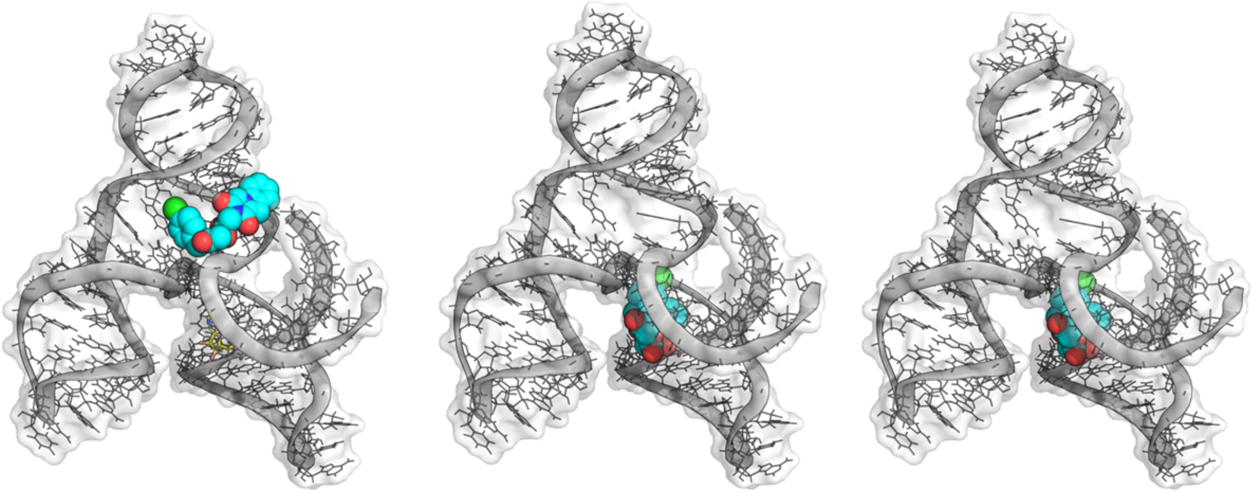

**Figure S34. Docked poses of the first three lowest energy poses of X1-D6.** The RNA is shown in cartoon, surface and lines. The cross-linking site is highlighted in yellow (sticks).

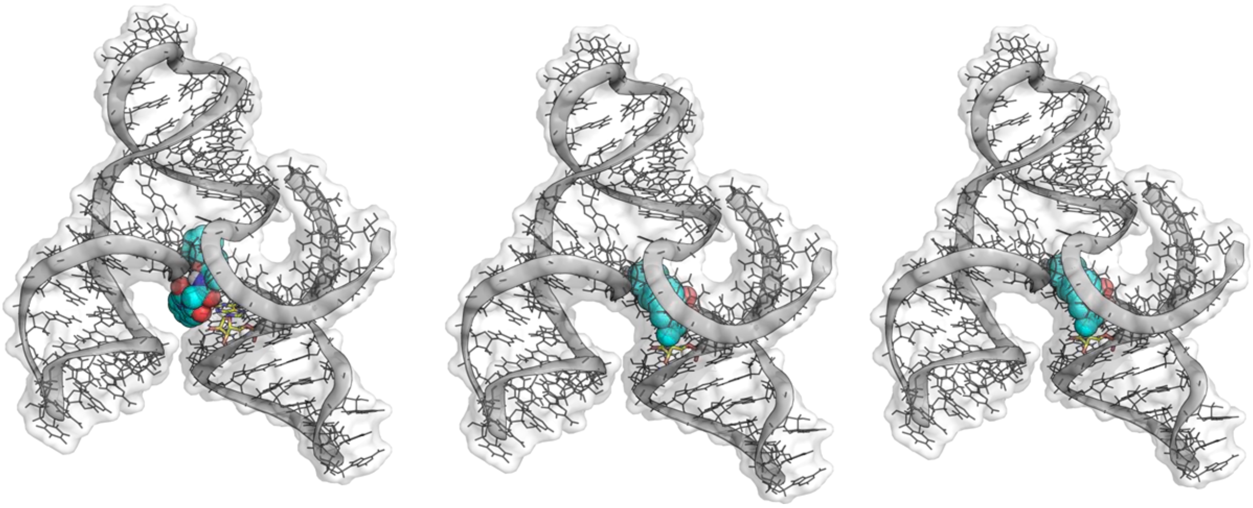

**Figure S35. Docked poses of the first three lowest energy poses of X1-D7.** The RNA is shown in cartoon, surface and lines. The cross-linking site is highlighted in yellow (sticks).

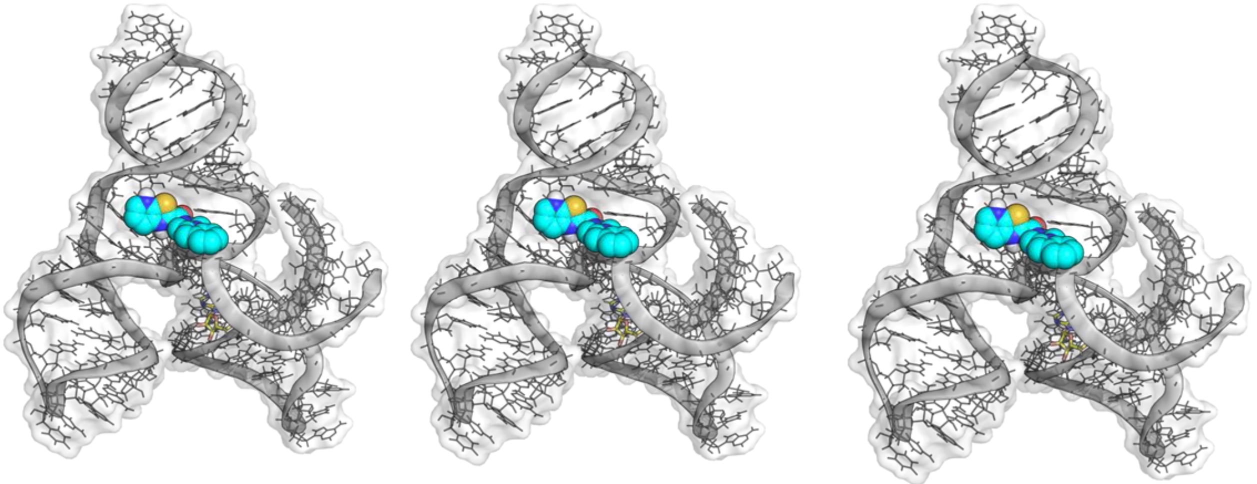

**Figure S36. Docked poses of the first three lowest energy poses of X1-D8.** The RNA is shown in cartoon, surface and lines. The cross-linking site is highlighted in yellow (sticks).

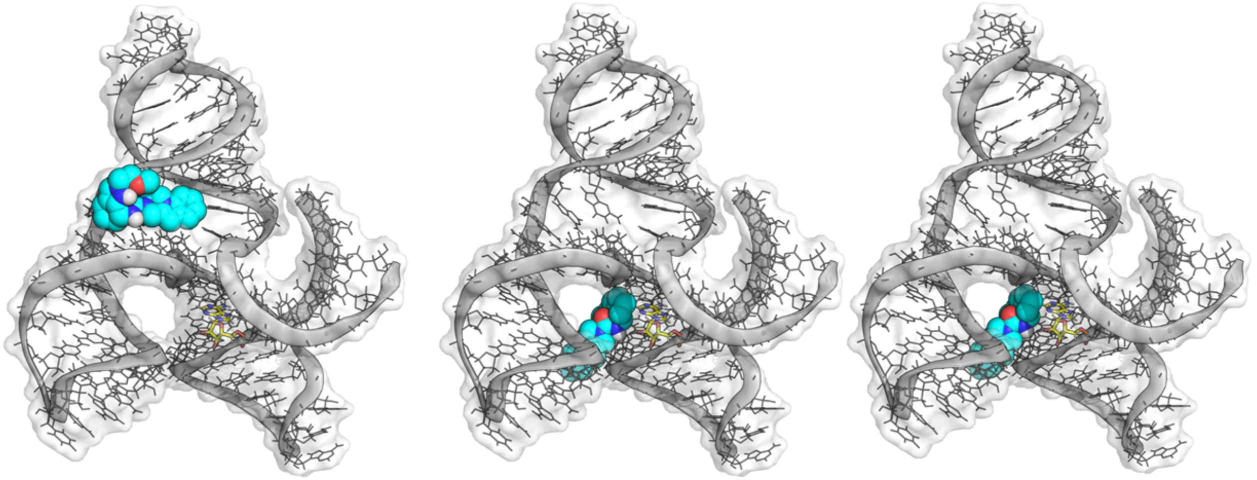

**Figure S37. Docked poses of the first three lowest energy poses of X1-D9.** The RNA is shown in cartoon, surface and lines. The cross-linking site is highlighted in yellow (sticks).

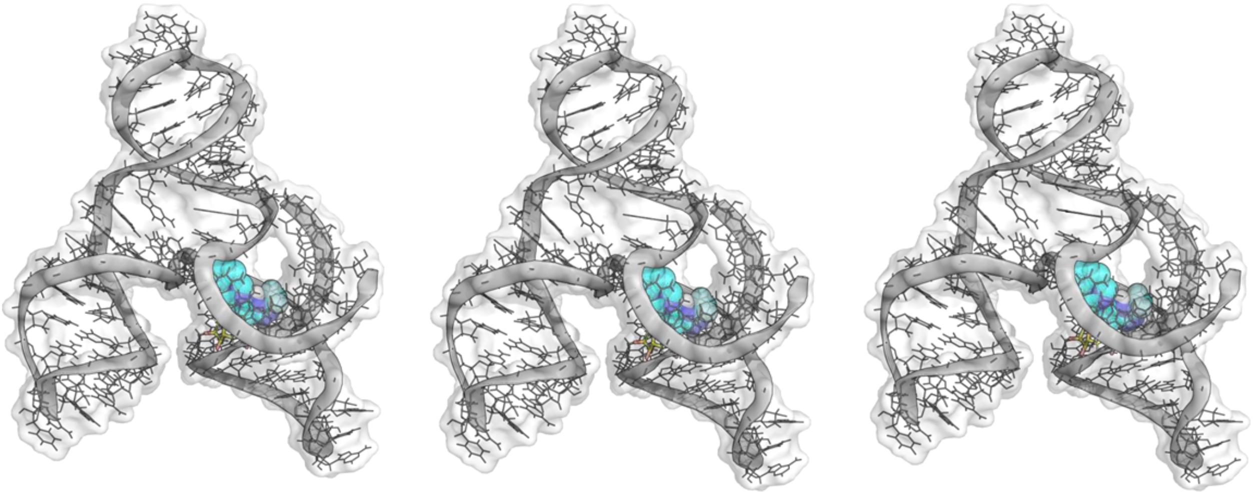

**Figure S38. Docked poses of the first three lowest energy poses of X1-D10.** The RNA is shown in cartoon, surface and lines. The cross-linking site is highlighted in yellow (sticks).

**Note S1. Chemotype analysis**

To determine whether binding fragments share common chemotypes, a traditional substructure analysis using DataWarrior (*openmolecules.org*)[8] was completed. Likely due to the small number of binding fragments and their structural diversity, statistically significant common scaffolds were unable to be identified. Therefore, a machine learning pipeline that combines cheminformatics tools and feature interpretation techniques was built. The pipeline uses SMILES (Simplified Molecular Input Line Entry System)[9] strings to represent molecules and extracts molecular features using different types of fingerprints. To represent comprehensively the structural features of small molecules, three distinct types of molecular fingerprints that encode chemical structure in a machine-readable format were employed. These included: (i) Morgan fingerprints, which captures the local structural environment around each atom within a certain radius and hence are also known as circular fingerprints; (ii) RDKit fingerprints, which map how atoms are connected through bonds, providing a detailed picture of the molecule's overall framework; and (iii) atom-pair fingerprints, which describe the types of atoms present and the distances between them, offering insight into the spatial relationships within the molecule. Together, these complementary fingerprints capture a wide range of structural and chemical information for RNA-binding and non-binding fragments.

We first tested whether each molecular fingerprint used alone could accurately classify binders and non-binders. Although each individual fingerprint representation yielded some level of predictive signal, the resulting area under the curve values of 65% (Morgan), 60% (atom-pair), and 68% (RDkit) indicate only modest discriminatory power. Although such values are above random chance (50%), they are not sufficiently reliable to accurately distinguish binders from non-binders. Thus, these molecular fingerprints were combined, that is an ensemble approach, was applied.

All three fingerprints (RDkit, Morgan, and atom-pair) were therefore used as input to develop a machine learning approach, here a random forest model, due to its ability to analyze high-dimensional data. The random forest model was developed by randomly dividing the data, into five parts or “folds”, where four folds were used for training and one for model testing. Our model makes decisions on the contributions of each substructure identified from the three types of molecular fingerprints to RNA binding, as assigned by SHapley Additive exPlanations (SHAP) analysis.

Biases in random forest models can be introduced by imbalances in the number of samples in each data set (minority vs. majority), here the number of binders (n = 23) and non-binders (n = 177), leading to overfitting of minority data. To address data imbalance, three different strategies were employed: (i) SMOTE (Synthetic Minority Over-sampling Technique) was implemented to create synthetic, nearest neighbor binders that were added to the minority data set; Tomek Links were used to remove noisy data near class (binder vs. non-binder) boundaries. That is, if a non-binder and binder were neighbors, the non-binder was removed from the majority data set; and (iii) a stratified K-fold cross-validation approach to avoid overfitting of the binder data set. This method involves dividing the data into multiple parts, here five, while maintaining a balanced representation of the different classes (e.g. binders vs. non-binders) in each split.

In brief, instead of duplicating existing binders, SMOTE interpolates between the feature vectors of two chemically similar binders to create a new, intermediate point. In practical terms, this means that the synthetic binders resemble real binders in their molecular fingerprints but do not correspond to actual tested chemical structures. They provide the model with additional “virtual” binders that improve balance and reduce bias during training. Typically, these synthetic samples show a moderate-to-high similarity to their parent binders, with Tanimoto coefficients in the range of ~0.8–0.9, which is often considered the threshold for meaningful chemical similarity. This ensures that the augmented data remain representative of the binder chemical space while preventing overfitting to a very small minority class. A Tomek Link occurs when a binder and a non-binder are each other’s nearest neighbors in feature space. In such cases, the non-binder (as the majority class member) is removed to reduce class overlap and improve model clarity. The number of non-binders removed by Tomek Links corresponds directly to the number of binder/non-binder nearest neighbor pairs identified, providing a measure of how frequently the two classes occupy overlapping regions in descriptor space.

As aforementioned, the model was trained on four folds while the fifth was used to test model accuracy. The data were randomly assigned five times to each of five folds, followed by training and testing to provide a more accurate estimate of model performance. During model training, a grid search was employed that tests different combinations of model settings, or “hyperparameters”, to identify the configuration that yields the best predictive accuracy. Six hyperparameters were used that define the trees used in the decision-making process (described in **Table S3**). Tuning these hyperparameters balances model complexity, prevents overfitting, and improves predictive performance on imbalanced molecular datasets. To optimize model performance, a grid search approach was employed to systematically explore combinations of hyperparameters. During the grid search, the model was trained and evaluated under each possible parameter combination using cross-validation, ensuring that performance estimates were unbiased. The resulting scores were then compiled into a performance surface (heatmap), where regions of higher accuracy (red) indicated favorable parameter combinations. This systematic search ensured that the final model was both well-calibrated and robust, avoiding suboptimal choices that might arise from manual tuning.

Finally, a confusion matrix was used for model evaluation, which is a classification evaluation tool that summarizes model predictions relative to the true class labels. It is organized into four categories: true positives (TP), correctly predicted binders; true negatives (TN), correctly predicted non-binders; false positives (FP), non-binders incorrectly classified as binders; and false negatives (FN), binders incorrectly classified as non-binders. By capturing both correct and incorrect classifications, the confusion matrix provides a detailed view of model performance beyond overall accuracy. In this study, the confusion matrix was used to quantify the predictive ability of the random forest model to distinguish binders from non-binders.

Model performance was then evaluated both receiver operating characteristic (ROC) analysis and confusion matrix classification outcomes. To determine the optimal classification threshold for the random forest model, we employed F1-score optimization, which balances precision (the fraction of predicted binders that are true binders) and recall (the fraction of actual binders that are correctly identified). Rather than applying the default threshold of 0.5, we systematically evaluated model performance across a range of thresholds using cross-validated predictions. The threshold yielding the maximum F1-score was selected, as it provides the best trade-off between identifying true binders while minimizing false positives. This procedure resulted in an optimal threshold of 0.34, which was subsequently applied throughout the analyses to improve the reliability of binder versus non-binder classification. This cut-off was applied as it allows the model to balance sensitivity and specificity, achieving high true negative accuracy while still capturing most binders. Adjusting this threshold upward or downward can shift the trade-off between false positives and false negatives, allowing prioritization of binder discovery versus strict rejection of non-binders depending on the application. This is a key element in this analysis such that the model assigns a compound as a binder or a non-binder accurately. The average ROC curve yielded a mean area under the curve of 0.92 ± 0.04, indicating excellent discriminative power of the random forest model in distinguishing binders from non-binders. This strong separation was further supported by the confusion matrix, which showed that most non-binders (34/35) and binders (11/14) were correctly classified.

This analysis verified that the random forest model can accurately identify substructures that contribute to RNA binding. Thus, a comparative enrichment analysis of features found exclusively or predominantly in binders revealed several key contributors. In summary, our machine learning approach can pinpoint structural features associated with RNA recognition, offering a tool to guide structure-activity relationship (SAR) studies and optimizing ligand design for improved potency and selectivity.

Machine Learning Pipeline for Predicting RNA Binding Fragments

This Python script implements a comprehensive machine learning pipeline for distinguishing RNA-binding from non-binding small molecules. It utilizes Random Forest (RF) classifiers trained on molecular fingerprints derived from SMILES strings, performs feature selection, evaluates model performance through cross-validation and SHAP analysis, and applies a dual-mode Leave-One-Binder-Out (LOBO) strategy for model validation.

1. Input and Preprocessing

Data Input

The script accepts a CSV file containing two columns:

- SMILES: Canonical SMILES strings representing the molecular structures.
- Label: Binary class label indicating binders (1) and non-binders (0).

Invalid SMILES strings are filtered out using RDKit. Any such SMILES are logged to invalid_smiles.log in the output directory.

2. Feature Extraction

Fingerprint Generation

Molecular features are generated using the following fingerprint types (user-selectable):

- Morgan (ECFP) fingerprints (radius=3, nBits=1024)
- MACCS keys (167-bit)
- RDKit topological fingerprints
- Hashed AtomPair fingerprints (nBits=1024)

Users may also ensemble multiple fingerprint types using the + symbol (e.g., morgan+rdkit).

3. Feature Selection

After fingerprint generation, feature selection is applied using a Random Forest model trained on the full feature set. The SelectFromModel method retains only features with importances above the median threshold, effectively reducing dimensionality and improving interpretability.

4. Model Training and Hyperparameter Optimization

Stratified K-Fold Cross-Validation

The dataset is split into k stratified folds (default k=5). For each fold:

- SMOTE oversampling is applied on the training set (with sampling_strategy=0.75) to mitigate class imbalance.
- A base Random Forest model (class_weight='balanced', n_estimators=100) is trained.

Randomized Hyperparameter Search

Once CV is completed, the script performs a RandomizedSearchCV to optimize the following RF hyperparameters:

- n_estimators, max_depth, min_samples_split, min_samples_leaf, max_features

The best model (based on mean AUC across folds) is retained for downstream evaluation. Average AUC is saved along with the optimal hyperparameter settings.

5. SHAP Analysis

The script performs SHAP (SHapley Additive exPlanations) analysis to interpret the model:

- SHAP values are calculated for each feature.
- Feature importance is computed as the mean absolute SHAP value.
- Visualization includes:
    - Bar and beeswarm summary plots
    - Dependence plots for top 5 features
    - Force plots for three representative samples

SHAP outputs are saved as .csv and .png files.

6. Confidence Threshold Tuning

A threshold for converting predicted probabilities into class labels is optimized based on:

- F1 score (default) or recall
- Thresholds in the range [0.05, 0.95] are evaluated, and the one maximizing the chosen metric is selected as best_threshold.

This value is later used in confirmation-mode LOBO evaluation.

7. Visualization and Diagnostics

The script generates the following performance plots:

- Train vs. Test AUC bar chart
- Learning curve showing AUC vs. training size
- Precision-recall curve

- Confusion matrix (with user-defined threshold)
- t-SNE 2D projection of molecular fingerprints
- Average AUC-ROC curve across cross-validation folds, including shaded error bands

8. Leave-One-Binder-Out (LOBO) Evaluation

LOBO simulates how well the model generalizes to unseen binders by:

- Holding out each binder molecule individually
- Training the model on all remaining data
- Predicting the probability of the held-out sample

Two evaluation modes are implemented:

- Screening mode: Uses a low threshold (0.25) to prioritize sensitivity
- Confirmation mode: Uses the optimized threshold from the confidence tuning step

Both modes output:

- Per-sample predictions and probabilities (lobo_results.csv)
- Confusion matrix and ROC curve

9. Command-Line Usage

bash

CopyEdit

```
python binder_vs_nonbinder_rf_LOBO_SMOTE_PERFOLD_V4_ENSEMBLE_V2.py \
  --input combined_smiles.csv \
  --output FINAL_LOBO_ENSEMBLE \
  --use_smote \
  --fingerprint morgan+rdkit \
  --kfolds 5
```

Arguments:

- --input: Input CSV file with SMILES and Label
- --output: Output directory
- --use_smote: Enable class balancing with SMOTE
- --fingerprint: Type or combination of fingerprints
- --kfolds: Number of K-folds for CV (default: 5)

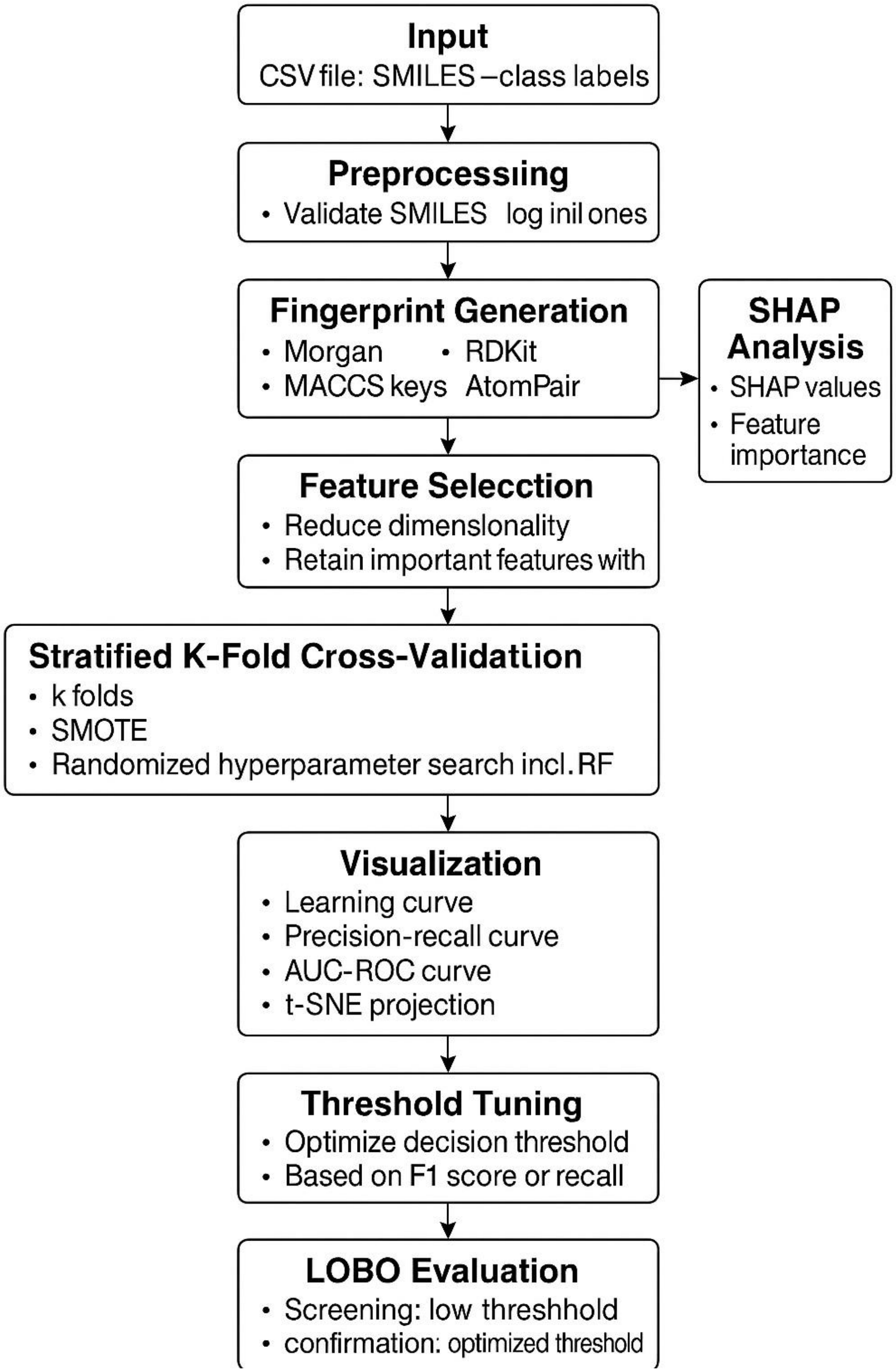
Input
CSV file: SMILES –class labels
Preprocessiing
• Validate SMILES log inil ones
Fingerprint Generation
• Morgan • RDKit
• MACCS keys AtomPair
SHAP Analysis
• SHAP values
• Feature importance
Feature Selecction
• Reduce dimenslonality
• Retain important features with
Stratified K-Fold Cross-Validatiion
• k folds
• SMOTE
• Randomized hyperparameter search incl. RF
Visualization
• Learning curve
• Precision-recall curve
• AUC-ROC curve
• t-SNE projection
Threshold Tuning
• Optimize decision threshold
• Based on F1 score or recall
LOBO Evaluation
• Screening: low threshhold
• confirmation: optimized threshold

**Note S2. Automated 3D structure generation of RNA using FARFAR2 and molecular docking.** To gain insight into the potential molecular recognition of target RNAs by **X1** across the transcriptome, a multi-step computational pipeline integrating secondary structure prediction, 3D structure modeling, and molecular docking was employed. Three-dimensional models of all 106 transcripts (corresponding to 58 genes) for **X1** that could be determined from the Chem-CLIP-Map-Seq pipeline were generated from the MFE secondary structures obtained from ScanFold analysis using the Fragment Assembly of RNA with Full-Atom Refinement version 2 (FARFAR2[10]) protocol within the Rosetta suite. FARFAR2 performs fragment-based assembly of RNA structures, that is 3D models are built by piecing together small structural elements (fragments) derived from known RNA structures. This is followed by high-resolution all-atom refinement to yield energetically favorable conformers. In FARFAR2 modeling, thousands of RNA 3D structures (decoys) are generated during stochastic sampling. Each decoy is assigned an energy score reflecting how well it satisfies Rosetta's all-atom energy function. Selecting the lowest-energy structures isolates those most thermodynamically favorable and structurally consistent with the underlying physics-based model. Extracting the top-scoring (lowest-energy) structures therefore focuses on conformations most likely to represent the native RNA fold, providing a rational basis for assessing model convergence and structural reliability.

For each transcript, ten lowest energy models were generated as ranked by Rosetta energy score. Rosetta score values (REU) were used to rank FARFAR2 models. Stability was evaluated based on $\Delta E$ relative to the minimum-energy model, with models within 2 REU considered energetically similar. These 10 lowest free energy models were then used for ensemble docking. The resulting RNA 3D models were subjected to blind docking using molecular docking algorithms (AutoDock-GPU[11] to probe potential ligand binding pockets across the entire surface of the RNA structure. Although the binding region was used as *a priori* knowledge from Chem-CLIP-Map-Seq studies, the cross-linked nt(s), were not used as restraints in blind docking studies.

To assess the reliability of the blind docking procedure, we compared predicted binding poses with experimental data. Crosslinking-based transcriptome-wide Chem-CLIP studies provide residue-level information on ligand–RNA contacts, which can serve as an orthogonal benchmark for docking results. By measuring the spatial agreement between the docked ligand positions and experimentally identified cross-linked nucleotides, we established a quantitative criterion to validate docking accuracy. To do so, for each transcript targeted by X1, we considered an ensemble of ten FARFAR2-generated RNA structural models. The first three lowest energy docked poses of X1 against each ensemble were selected for further analysis. These poses were then evaluated for congruence with the experimentally identified cross-linked site from transcriptome-wide Chem-CLIP studies. In brief, the ligand and the cross-linked RNA residue were extracted and the center of mass for both groups of atoms was calculated.

The Euclidean distance between these two COMs served as a geometric measure of proximity. If the COMs were within a 10 Å distance, the docking pose was classified as successful. Indeed, substantial majority of docked poses (68.7% success rate), including for *MPP7* and *SSC4D*, was successfully localized near their designated RNA binding sites. For this subset of geometrically valid poses, the relative orientation of the diazirine moiety was then examined by calculating the vector from the diazirine center toward the *trans*-FFF site and comparing it across models. Because the FFF site does not always correspond to a single nucleotide with absolute precision owing to experimental uncertainties in cross-link localization the COM was calculated using a window encompassing the cross-linked residue and its adjacent nucleotides (± 5 positions). Docked poses with a diazirine-to-site COM distance ≤ 8 Å were classified as spatially consistent with the Chem-CLIP data. For this subset of geometrically valid poses, the orientation of the diazirine group was then analyzed by defining a vector from the diazirine center toward the site COM, allowing assessment of whether the reactive group was favorably oriented toward the cross-linked region providing a 39% success rate.

To efficiently generate 3D RNA structures at scale, we implemented a batch-processing pipeline that automates Rosetta's rna_denovo execution using the SLURM workload manager. The pipeline is designed to take in a large collection of RNA sequences and corresponding secondary structures and submit SLURM jobs to generate low-energy RNA 3D models using the FARFAR2 protocol.

Input Data Organization

Each modeling job is contained in its own folder within a common base directory. Every folder must contain:

- A .fasta file: specifying the RNA sequence
- A .secstruct file: providing the corresponding dot-bracket secondary structure

The folder names are arbitrary but are expected to be uniquely named and organized within a single base directory provided to the script.

Script Functionality

The script (create_slurm_script_V3_batch_process.py) performs the following steps:

1. Batch Directory Traversal

The script traverses the provided base directory and groups subfolders into batches of user-defined size (default: 500). Each batch is submitted sequentially, with manual confirmation required before submitting the next batch.

2. Redundancy Check

Before submitting a SLURM job for a folder, the script checks whether the modeling output already exists by looking for a file named rna_output.silent. If this file is present, the job is skipped to prevent reprocessing.

3. SLURM Script Generation

For each valid folder, a job script named run_job.sh is generated with the following parameters:

```
#SBATCH --job-name=rna_record
#SBATCH --ntasks=1
#SBATCH --cpus-per-task=4
#SBATCH --mem=20G
#SBATCH --time=3-00:00
#SBATCH --account=scripps-dept
#SBATCH --qos=scripps-dept-b
```

The SLURM script loads the appropriate Rosetta module and executes the following Rosetta commands:

a. FARFAR2 Modeling

bash

```
rna_denovo.default.linuxgccrelease \
  -fasta [input.fasta] \
  -secstruct_file [input.secstruct] \
  -minimize_rna true \
  -cycles 100 \
  -nstruct 100 \
```

```
  -out:file:silent rna_output.silent \
  -out:level 500
```

- -fasta and -secstruct_file: input sequence and secondary structure
- -minimize_rna: performs full-atom minimization
- -cycles 100: sets the number of Monte Carlo sampling cycles
- -nstruct 100: generates 100 candidate models
- -out:file:silent: stores all models in compressed Rosetta silent file format
- -out:level 500: ensures detailed logging of the output

b. Structure Extraction

After modeling, the script extracts the top 3 lowest-energy structures from the silent file using:

bash

```
extract_pdbs.linuxgccrelease \
  -in:file:silent rna_output.silent \
  -in:file:tags $(grep "SCORE:" rna_output.silent | sort -nk2 | head -n 3 | awk '{print $NF}')
```

This ranks the structures by total Rosetta score and converts the selected models into standard .pdb files for visualization and downstream analysis.

Job Submission

Once the SLURM script is written, the script changes the working directory to the respective folder and submits the job via:

bash

```
sbatch run_job.sh
```

Any output or errors during job submission are logged to the terminal. The script then reverts to the original working directory to continue batch processing.

Command-Line Usage

The pipeline is executed as follows:

bash

python create_slurm_script_V3_batch_process.py /path/to/modeling_directories --batch_size 500

Arguments:

- base_directory: Path to the directory containing modeling subfolders
- --batch_size: Optional parameter specifying how many folders to process per batch (default: 500)

Post-processing

After job completion, each folder will contain:

- rna_output.silent: All 100 models in compressed format
- Top 3 .pdb files: Extracted low-energy 3D models
- farna_rebuild.out: Log file with runtime information and score summaries

**Note S3. Automated extraction of top-scoring RNA 3D structures from FARFAR2 output.** Following 3D RNA structure generation via Rosetta's FARFAR2 protocol, each modeling job produces a large ensemble of candidate conformations stored in a .silent file format. To facilitate analysis, visualization, or benchmarking, we implemented an automated shell script to extract the top 10 lowest-energy 3D RNA models from each modeling directory.

Input Requirements

The script processes a directory of RNA modeling subfolders, each of which must contain:

- rna_output.silent: A Rosetta silent file containing 100 models generated by FARFAR2 (rna_denovo)

Each subdirectory corresponds to one RNA target and is expected to be named arbitrarily but distinctly. The path to the parent directory containing these subfolders is supplied as the script's sole argument.

Script Functionality

The script (Generate_top10_structure_from_farfar.sh) performs the following steps for each modeling directory:

1. Input Validation

- The script confirms that exactly one command-line argument (a valid directory path) is provided.
- If the specified path does not exist or is not a directory, the script exits with an error.

2. Subdirectory Traversal

- It loops through all subdirectories within the specified parent directory.
- For each folder, it checks whether a file named rna_output.silent is present.

3. Rosetta Module Loading

To ensure Rosetta tools are available in the environment, the script loads the appropriate version of the Rosetta module:

bash

```
module purge

ml rosetta/2023.11.30
```

This ensures a clean module environment and loads Rosetta version 2023.11.30.

4. Extraction of Top 10 Structures

If the silent file exists, the script uses Rosetta's built-in tool extract_pdbs.linuxgccrelease to extract the top 10 structures with the lowest total energy scores.

The command used is:

bash

```
extract_pdbs.linuxgccrelease \
  -in:file:silent rna_output.silent \
  -in:file:tags $(grep "SCORE:" rna_output.silent | sort -nk2 | head -n 10 | awk '{print $NF}')
```

This command performs the following:

- Searches for lines beginning with SCORE: in the silent file.
- Sorts them numerically by the total score (2nd column).
- Extracts the top 10 tags (structure identifiers).
- Invokes extract_pdbs with these tags to generate standard .pdb files.

Each .pdb file represents one of the 10 best-scoring FARFAR2 models and is saved in the corresponding subdirectory.

Execution

The script is invoked using the command:

bash

bash Generate_top10_structure_from_farfar.sh /path/to/modeling_folders/

No other arguments are required. Upon execution, the script prints progress for each folder, lists folder contents for debugging, and handles errors gracefully (e.g., missing silent files or inaccessible folders).

Output

For each processed directory:

- Up to 10 .pdb files are generated (one per structure)
- If rna_output.silent is missing, the directory is skipped with a warning message

**Note S4. Analysis of statistically significant molecular descriptors distinguishing binders from non-binders.** Additional molecular features that contribute to the differentiation between binder and non-binder fragments were next investigated using 2D and 3D descriptors. The former includes properties such as molecular weight, number of carbon atoms or aromatic rings, while 3D descriptors include charged partial surface area (CPSA) and moment of inertia (MOI), among others. A total of ~1,800 descriptors were computed for each fragment using the Mordred[12] descriptor engine and the statistical significance of these descriptors for binders vs. non-binders was calculated through univariate statistical testing, particularly a Mann-Whitney U test[13].

To assess chemical property differences between two ligand sets (e.g., RNA binders vs. non-binders), we employed a Python script that calculates, compares, and visualizes molecular descriptors derived from SMILES strings. The pipeline integrates RDKit, Mordred, and 3D optimization routines to generate 2D and 3D descriptors, normalize them, and identify statistically significant differences between the two groups.

### Input

The script accepts two plain-text files containing SMILES strings, one per line, representing two distinct molecular sets. These may correspond to "binders" and "non-binders" or any user-defined categories.

Example input format:

makefile

CopyEdit

File1: binder_smiles.txt

File2: nonbinder_smiles.txt

### SMILES validation and 3D geometry generation

SMILES strings were sanitized using RDKit's MolFromSmiles() and SanitizeMol() functions. Molecules with invalid or non-sanitizable structures were discarded. Valid molecules were then hydrogenated (Chem.AddHs()) and converted into 3D conformations using:

- AllChem.EmbedMolecule(): generates 3D coordinates
- AllChem.UFFOptimizeMolecule(): optimizes geometry with the Universal Force Field

The combined 3D structures for both sets were exported to an .sdf file (structures.sdf) for archival and downstream use.

### Descriptor Calculation

We computed molecular descriptors using Mordred, a comprehensive cheminformatics descriptor calculator.

- 2D descriptors: calculated directly from sanitized SMILES.
- 3D descriptors: calculated from geometry-optimized conformers.

Each descriptor set was filtered to:

- Drop columns containing any NaN values
- Remove descriptors with all-zero values across all molecules

The resulting 2D and 3D descriptor matrices were then concatenated to form comprehensive feature sets for each group.

### Normalization

All descriptor values were normalized using Min-Max scaling independently for each group. This ensured that descriptors were on a uniform scale while preserving relative differences within each set.

### Statistical Comparison

We compared descriptors between the two groups using one of the following user-selectable tests:

- Student’s t-test (default): Welch's correction for unequal variances
- Mann–Whitney U test: non-parametric alternative for ordinal/rank-based comparison

Only descriptors common to both groups and fully numeric were retained. P-values were calculated for each descriptor and sorted in ascending order.

### Visualization

The top N most significantly different descriptors (default N = 5) were visualized using:

- Box plots or violin plots
- Seaborn and statannotations for aesthetic and annotated significance
- Plots were saved as .png images and labeled by descriptor name

For each descriptor, a side-by-side .csv was also generated, containing raw values for the binder and non-binder sets in separate columns for downstream statistical analysis.

### Output

- structures.sdf: Combined 3D structures of both groups
- <output_file>.csv: Summary of all tested descriptors with associated p-values
- /plots/: Directory containing:
    - Annotated plots of top differing descriptors
    - Raw descriptor value tables for top hits

Execution

The script is run from the command line as follows:

bash

CopyEdit

```
python mordred-compare_smiles_plot_2D_3D_V6.py \
    binder_smiles.txt \
    nonbinder_smiles.txt \
    comparison_results.csv \
    --output_dir plots \
    --num_properties 5 \
    --plot_type box \
    --test_type t-test
```

Optional arguments:

- --num_properties: Number of top descriptors to plot (default: 5)
- --plot_type: box or violin (default: box)
- --test_type: t-test or mannwhitneyu (default: t-test)

**Note S5. Automated analysis of RNA secondary structures and motifs from dot-bracket notation**

To evaluate RNA structural motifs across multiple genes, we implemented a Python-based pipeline that parses dot-bracket secondary structures, detects internal loops and tetraloops, classifies crosslink sites, and generates quantitative and graphical summaries. The input is a curated Excel file containing RNA sequences, dot-bracket structures, and annotated crosslinked nucleotide positions.

Input

The input file is an Excel spreadsheet with the following required columns:

- Gene: Gene name or identifier
- seq: RNA nucleotide sequence (5'→3')

- structure: Dot-bracket representation of the RNA secondary structure
- trans_FFF_site: Integer index (0-based) indicating the experimentally crosslinked nucleotide

The script ensures sequence and structure lengths match, and skips entries with mismatches.

Structural Motif Parsing

Using a custom parser, the script traverses the dot-bracket notation and identifies:

1. Base Pairing

- A stack-based algorithm is used to match ( with ) and determine base-paired indices.

2. Tetraloops

Hairpin loops flanked by a single helix and consisting of exactly 4 unpaired nucleotides were evaluated. If the loop sequence matched the following patterns, they were categorized:

- GNRA: regex G.[AG]A
- UUCG and CUUG: recognized explicitly
- Other: all remaining tetraloops

Detected tetraloops were annotated by range, sequence, and category.

3. Internal Loops

Non-base-paired regions flanked by paired bases were classified as internal loops. For each:

- The loop start/end positions were recorded
- Loop length and sequence were saved
- Multiple unpaired regions within stems were handled using a visited index set

Crosslink Location Classification

The crosslinked nucleotide position was examined in the context of its structural role using the dot-bracket notation. Each position was classified as:

- "Loop/Unpaired": corresponds to .
- "Helix (base-paired)": corresponds to ( or )

- "Out of range": invalid index relative to structure length

This enables mapping of experimental crosslinking to structural contexts.

Output and Summary Table

The script creates a structured .csv file summarizing the following for each gene:

| **Column Name** | **Description** |
|---|---|
| Gene | Gene identifier |
| Length | Total length of RNA sequence |
| Crosslink_nt | 1-based index of crosslinked nucleotide |
| Crosslink_location | Structural category (loop, helix) |
| Internal_loops | Total number of internal loops |
| Avg_internal_loop_size | Mean loop size (rounded to 2 decimal places) |
| Tetraloop_GNRA | Count of GNRA tetraloops |
| Tetraloop_UUCG | Count of UUCG tetraloops |
| Tetraloop_CUUG | Count of CUUG tetraloops |
| Tetraloop_Other | Count of other tetraloops |

Graphical Summary and Visualization

The script automatically generates and saves several bar plots:

1. Internal Loops
   - Count per gene
   - Average loop size per gene
2. Tetraloop Subtypes
   - Separate plots for GNRA, UUCG, CUUG, and Other tetraloops
3. Crosslink Location
   - Distribution of crosslinked nucleotide positions by structural type (e.g., loop vs. helix)

Color coding is applied to highlight specific genes of interest (SSC4D = red, MPP7 = blue, others = gray).

Plots are saved in PNG format in the working directory.

**Execution**

The script is run from the command line:

python analyze_structures.py X1_cleaned.xlsx --output structure_motif_summary.csv

Optional arguments:

- --output: Output path for summary CSV file (default: structure_motif_summary.csv)

**Note S6. Automated extraction and interaction profiling of RNA–Ligand complexes using fingeRNAt.** To systematically characterize RNA–ligand interactions across a large set of RNA-ligand complexes, we developed a Python-based pipeline that:

1. Splits RNA and ligand coordinates from a complex .pdb file using PyMOL,
2. Converts ligand files to .sdf format using Open Babel,
3. Executes the fingeRNAt[14] toolkit to identify and classify molecular interactions between RNA and small molecules.

Input Format

The script expects a directory containing RNA-ligand complexes in PDB format. Each file should:

- End with .pdb
- Be named using the pattern RNA_ligand_*.pdb

This naming convention ensures the correct files are filtered and processed automatically.

Workflow Overview

1. Directory Setup

For each RNA-ligand complex, a new folder is created in a processed_complexes/ subdirectory. Each folder is populated with:

- The original .pdb file
- A copy of the required fingeRNAt scripts and modules:
    - fingeRNAt.py, preprocessing.py, fingerDISt.py, DistanceMetrics.py, config.py, gui.py

This ensures self-contained processing environments for each complex.

2. RNA and Ligand Extraction Using PyMOL

Using the PyMOL Python API, the complex structure is split into:

- RNA-only coordinates saved as a .pdb file (residues A, C, G, U, ADE, CYT, GUA, URA)
- Ligand coordinates saved in .mol2 format by selecting residues with name UNL

3. Ligand Format Conversion

The ligand .mol2 file is converted into .sdf format using Open Babel via the following shell command:

bash

CopyEdit

obabel -imol2 ligand.mol2 -osdf -O ligand.sdf

Any failures in ligand conversion raise an error and skip the current complex.

4. RNA–Ligand Interaction Analysis with fingeRNAt

The processed RNA and ligand files are passed to fingeRNAt.py, a toolkit designed to extract and classify interactions such as:

- Hydrogen bonds
- π–π stacking
- Electrostatic interactions
- Hydrophobic contacts

The analysis is run in detailed and verbose mode with the following command:

bash

CopyEdit

./fingeRNAt.py -r RNA.pdb -l ligand.sdf -detail -print -verbose

The script captures and prints the output from fingeRNAt, including:

- Preview of RNA input file
- Success/failure of the interaction analysis
- Any error messages or interaction statistics

Error Handling

The script includes extensive exception handling:

- Checks for missing dependencies (e.g., PyMOL modules or fingeRNAt scripts)
- Skips malformed or incompatible .pdb files
- Captures and logs errors from subprocesses (Open Babel or fingeRNAt)

Execution

The script is run from the command line as:

python process_rna_ligand_complexes_with_fingeRNAt_V5.py /path/to/complexes

Where /path/to/complexes is a directory containing RNA_ligand_*.pdb files.

Output

For each RNA-ligand complex:

- A separate subdirectory is created under processed_complexes/
- It contains the split RNA.pdb, ligand.sdf, fingeRNAt scripts, and any output logs or results
- Interactions are printed to the console and optionally logged for further analysis

**Docking success evaluation**

Ligand Atom Identification

Using BioPython's PDBParser, the script identifies atoms belonging to the ligand by selecting heteroatoms with residue IDs starting with "H" or "H_". These atoms are aggregated across all models in the PDB structure.

Target Site Atom Identification

RNA atoms corresponding to the crosslinked nucleotide (as per the trans_FFF_site) are extracted using standard residue ID matching. Only canonical residues (i.e., not heteroatoms or water) are considered.

Distance Calculation

The minimum Euclidean distance (in Å) between any ligand atom and any RNA atom at the crosslink site is computed using:

```
np.linalg.norm(coords1[:, None, :] - coords2[None, :, :], axis=-1).min()
```

A docking pose is classified as a success if this distance is less than or equal to the user-specified cutoff (e.g., 5.0 Å), and fail otherwise.

Output Summary

CSV File

A summary table is written to docking_success_summary.csv in the output directory. Columns include:

- pdb_file: Name of the docked complex file
- ENST: Transcript ID
- trans_FFF_site: Crosslinked nucleotide index
- min_distance: Minimum computed distance (Å)
- success: Boolean flag indicating docking success based on cutoff

Bar Plot

A bar plot is generated to visualize the number of successful vs. unsuccessful docked poses. The plot:

- Is saved as docking_success_plot.png
- Uses green for success, red for failure
- Displays a count of poses in each category

Command-Line Usage

The script is executed as:

```
python evaluate_docking_and_plot_excel.py \
```

```
--excel X1_cleaned.xlsx \
--pdb_dir docked_models/ \
--cutoff 5.0 \
--output_dir docking_results/
```

Arguments:

- --excel: Path to the Excel file with the X1 sheet
- --pdb_dir: Directory containing docked PDB files
- --cutoff: Distance threshold in Å for defining docking success
- --output_dir: Output directory for CSV and plot (default: docking_output)

**Evaluation of diazerine orientation in docked poses**

To assess the geometric congruence between computationally docked ligand poses and experimentally defined cross-linking sites, we developed a Python-based evaluation workflow implemented in the script evaluate_docking_diazirine_to_site_V3_window.py. This analysis quantified the distance between the diazirine moiety of each docked ligand and the experimentally mapped RNA residue identified as the *trans*-FFF cross-linking site. The approach integrates structural parsing, automated center-of-mass [7] computation, and residue-based filtering, enabling systematic evaluation of docking accuracy across large sets of RNA–ligand complexes.

*Input Data*

The program requires two primary inputs:

1. Docking results — A directory containing all ligand–RNA complex structures in PDB format, where filenames encode the target transcript identifier (ENST).
2. Experimental mapping table — An Excel file containing a sheet named “X1”, which lists each transcript (ENST), its corresponding *trans*-FFF cross-linking residue position (trans_FFF_site), and optionally thermodynamic or statistical descriptors such as minimum free energy (MFE) or Z-score for resolving duplicate entries.

*Algorithmic Workflow*

The script systematically processes all PDB files in the specified directory using the following steps:

1. Parsing and Target Resolution
   For each docked structure, the corresponding ENST identifier is extracted from the filename using a regular expression. The residue number of the experimentally mapped cross-linking site is then determined from the Excel table. If multiple entries exist for the same transcript, the site is resolved according to a user-defined strategy (e.g., lowest MFE or Z-score, or first entry in the table).

2. Site Atom Selection and Center-of-Mass Calculation
   Using the Biopython PDBParser, the program isolates all atomic coordinates from the nucleotide at the trans_FFF_site. Optionally, residues within a user-specified window (±*K* residues) around the cross-linking site can be included to approximate a local binding pocket. The COM (or centroid) of the selected atoms is calculated, either using equal weights or optionally mass-weighted based on atomic species (H, C, N, O, S, P, halogens).

3. Ligand and Diazirine Identification
   The ligand is identified as a non-polymeric residue (HETATM record) within the PDB file. If the ligand residue name is provided (e.g., *X1*), it is selected explicitly; otherwise, the residue with the largest number of atoms is assumed to be the ligand.
   The diazirine reactive group within the ligand is then localized by one of two methods:

   - User-specified atom list: A comma-separated list of atom names (e.g., C7,N8,N9) provided via command-line argument.
   - Automatic detection: Identification of the most compact C–N–N triplet in the ligand, using a geometric constraint that the interatomic distances between the three atoms are ≤1.8 Å (typical for diazirine rings).

The geometric center (or mass-weighted COM) of the selected diazirine atoms is calculated to represent the photoreactive centroid.

4. Distance Computation and Pose Classification
   The Euclidean distance between the ligand's diazirine center and the RNA site COM is computed. Docking poses are classified as *successful* if this distance is less than or equal to a user-defined threshold (default 12 Å), corresponding to a realistic cross-linking range for photoaffinity labeling experiments.
   For each structure, metadata including filename, ENST ID, ligand name, number of diazirine atoms, computed distance, and success/failure status are stored in a results table.

5. Output and Visualization
   All results are saved as a CSV file (diazirine_to_site_summary.csv) in the output directory, accompanied by a bar plot summarizing the fraction of successful

versus unsuccessful docking poses. This summary enables rapid visualization of docking accuracy across transcripts.

| Parameter | Description |
| --- | --- |
| --excel | Input Excel file containing ENST and trans_FFF_site columns |
| --pdb_dir | Directory of docked PDB files |
| --output_dir | Directory for saving output CSV and plots |
| --site_window | Number of neighboring residues (±K) to include in site COM calculation |
| --cutoff | Distance threshold (Å) defining a successful docking pose |
| --ligand_resname | Optional ligand residue name (e.g., X1) |
| --site_chain | Optional chain ID restriction |
| --auto_diazirine | Automatically detect C–N–N diazirine triplet |
| --diazirine_atomnames | Comma-separated custom diazirine atom names |
| --mass_weighted | Apply atomic mass weighting for COM calculations |
| --duplicate_strategy | Rule for resolving duplicate ENST entries (e.g., mfe_min, zscore_min) |

*Implementation Details*

Programming environment: Python 3.

Libraries: numpy, pandas, matplotlib, tqdm, and Bio.PDB from Biopython.

Computational efficiency: Progress tracking is handled by tqdm to monitor per-file processing, and data aggregation is vectorized using pandas and NumPy for minimal overhead.

Visualization: Summary plots are generated automatically in PNG format using Matplotlib.

**Synthetic Methods and Characterization**

**Abbreviations:**

AcOH: Acetic acid

DCM: Dichloromethane

DIPEA: Diisopropyl ethyl amine

DMSO: Dimethyl sulfoxide

DMF: *N,N*-Dimethylformamide

EDCI: 1-Ethyl-3-(3-dimethylaminopropyl)carbodiimide hydrochloride

ESI: Electrospray ionization

HATU: Hexafluorophosphate azabenzotriazole tetramethyl uranium

HOAt: 1-Hydroxy-7-azabenzotriazole

HPLC: High-performance liquid chromatography

MeOH: Methanol

PyBOP: Benzotriazol-1-yloxytripyrrolidinophosphonium hexafluorophosphate

TFA: Trifluoroacetic acid

**General Synthetic Methods**. Chemicals were purchased from Sigma Aldrich, Enamine, or TCI. $^{1}$H NMR and $^{13}$C NMR spectra were collected on a 400 MHz UltraShieldTM or a 600 MHz UltraShieldTM NMR spectrometer (Bruker). All FFFs were acquired from ?. All chemical shifts are reported in ppm and coupling constants are reported in Hz.

Preparative HPLC was performed using a Waters 1525 Binary HPLC pump equipped with a Waters 2487 dual absorbance detector system and a Waters Sunfire C18 OBD 5 μm, 19 ×150 mm S-14 column. A linear gradient from 0-100% MeOH in water containing 0.1% (v/v) TFA and a flow rate of 5 mL/min was applied over 60 min, and absorbances were monitored at 254 and 280 nm in parallel. The purity of final products was assessed by analytical HPLC using a Waters Symmetry C18 5 μm, 4.6 × 150 mm column with a flow rate of 1 mL/min and a linear gradient from 0-100% MeOH in water with 0.1% (v/v) TFA applied over 60 min, and absorbances were monitored at 254 and 345 nm.

Mass spectra were acquired using positive ESI mode on an Orbitrap Exploris 120

(Thermo Fisher Scientific) coupled to a Vanquish HPLC system (Thermo Fisher Scientific).

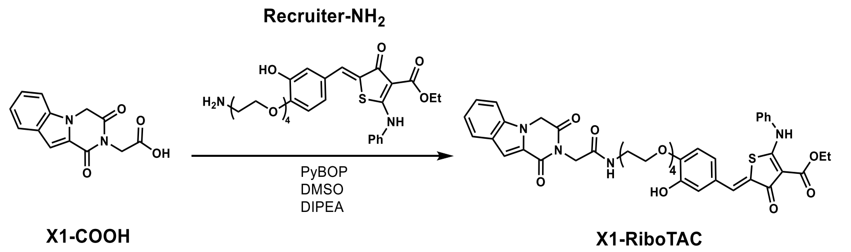


**Scheme S1. Synthesis of X1-RiboTAC.**

**Synthesis of X1-RiboTAC.** **Recruiter-NH2** was synthesized as previously reported.[15] To a solution of **X1-COOH** (12.5 mg, 0.05 mmol) in 0.1 mL of DMSO was added PyBOP (41.6 mg, 0.08 mmol) and DIPEA (27 µL, 0.15 mmol). The reaction was stirred at room temperature for 30 min followed by the addition of **Recruiter-NH2** (35 mg, 0.08 mmol). The reaction was stirred at room temperature for 16 h and then diluted by adding 1 mL of MeOH containing 0.1% (v/v) TFA. The product **X1-RiboTAC** (16% yield) was purified by HPLC as described in the **General Synthetic Methods**. $^{1}$H NMR (600 MHz, DMSO) δ 11.24 (s, 1H), 9.43 (s, 1H), 8.25 (s, 1H), 7.78 (d, J = 8.1 Hz, 1H), 7.63 – 7.59 (m, 1H), 7.52 (s, 4H), 7.48 (s, 1H), 7.46 – 7.43 (m, 1H), 7.41 (ddd, J = 8.3, 6.9, 1.1 Hz, 1H), 7.38 – 7.36 (m, 1H), 7.21 (ddd, J = 8.0, 6.9, 0.9 Hz, 1H), 7.01 (s, 1H), 6.97 (s, 2H), 5.30 (s, 2H), 4.45 (s, 2H), 4.28 (d, J = 7.1 Hz, 2H), 4.11 (s, 2H), 3.76 – 3.73 (m, 2H), 3.58 (d, J = 5.4 Hz, 2H), 3.52 (dd, J = 16.3, 4.0 Hz, 7H), 3.40 (s, 2H), 3.31 (s, 1H), 3.21 (d, J = 5.7 Hz, 2H), 1.29 (t, J = 7.1 Hz, 3H). $^{13}$C NMR (151 MHz, DMSO) δ 181.41, 175.80, 167.38, 166.81, 165.34, 158.56, 149.21, 147.49, 137.97, 136.99, 130.49, 130.11, 128.66, 126.94, 126.77, 126.09, 125.87, 125.23, 125.08, 123.57, 123.20, 121.94, 116.30, 114.19, 111.74, 107.01, 97.33, 70.40, 70.27, 70.19, 70.14, 69.46, 69.30, 68.39, 59.98, 47.00, 41.93, 39.25, 14.88. HRMS calculated for $[C_{41}H_{43}N_4O_{11}S]^+$ 799.2644, found 799.2645.

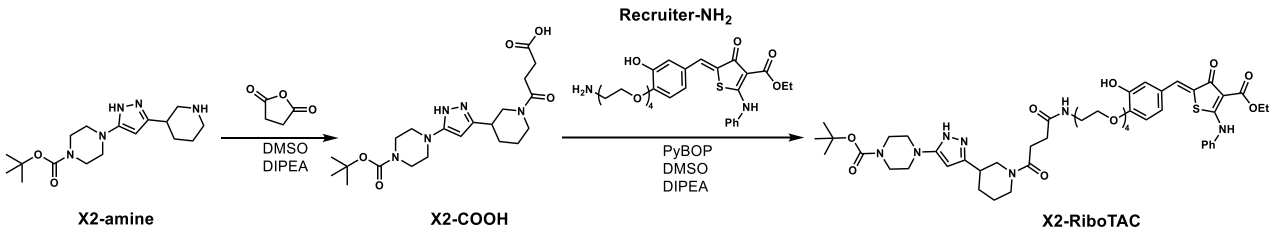


**Scheme S2. Synthesis of X2-RiboTAC.**

**Synthesis of X2-RiboTAC.** To a solution of **X2-amine** (10 mg, 0.03 mmol) in 0.1 mL of DMSO was added succinic anhydride (20 mg, 0.2 mmol) and DIPEA (18 µL, 0.1 mmol). The reaction was stirred at room temperature for 16 h and diluted by adding 1 mL of MeOH containing 0.1% (v/v) TFA, followed by purification by HPLC as described in the **General Synthetic Methods.** The intermediate **X2-COOH** was dissolved in 0.1 mL of DMSO followed by the addition of PyBOP (31.2 mg, 0.06 mmol) and DIPEA (18 µL, 0.1 mmol). The reaction was stirred at room temperature for 30 min followed by the addition of **Recruiter-NH$_2$** (26 mg, 0.06 mmol). The reaction was stirred at room temperature for 16 h amd then diluted by adding 1 mL of MeOH containing 0.1% (v/v) TFA. The product **X2-RiboTAC** (22% yield) was purified by HPLC as described in the **General Synthetic Methods**. $^1$H NMR (600 MHz, DMSO) δ 11.24 (s, 1H), 7.87 (dt, J = 8.8, 4.1 Hz, 1H), 7.59 – 7.40 (m, 6H), 7.04 – 6.94 (m, 3H), 5.81 (d, J = 13.3 Hz, 1H), 4.27 (q, J = 7.1 Hz, 6H), 4.13 – 4.09 (m, 4H), 3.96 (s, 4H), 3.76 – 3.72 (m, 2H), 3.61 – 3.55 (m, 3H), 3.55 – 3.45 (m, 7H), 3.43 – 3.34 (m, 6H), 3.20 – 3.14 (m, 3H), 3.12 (p, J = 3.0 Hz, 3H), 3.04 (tq, J = 11.1, 3.4 Hz, 1H), 2.73 (dtd, J = 24.3, 11.2, 7.4 Hz, 1H), 2.60 (d, J = 6.7 Hz, 1H), 2.33 (dt, J = 16.1, 7.0 Hz, 2H), 1.41 (s, 7H), 1.29 (t, J = 7.1 Hz, 3H). $^{13}$C NMR (151 MHz, DMSO) δ 181.41, 175.80, 172.01, 171.95, 171.19, 170.34, 170.18, 165.34, 159.09, 158.84, 158.60, 158.36, 157.60, 157.33, 154.36, 149.21, 147.51, 137.97, 130.10, 128.65, 126.08, 125.11, 123.56, 116.95, 116.32, 115.02, 114.22, 113.83, 97.35, 89.27, 89.19, 79.59, 70.39, 70.27, 70.20, 70.06, 69.60, 69.31, 68.41, 59.98, 55.53, 51.73, 49.96, 49.07, 47.77, 47.64, 46.08, 45.57, 41.85, 40.51, 39.04, 34.63, 34.05, 30.76, 30.18, 30.02, 29.22, 28.50, 28.21, 25.19, 24.57, 14.87. HRMS calculated for $[C_{49}H_{66}N_7O_{12}S]^+$ 976.4490, found 976.4485.

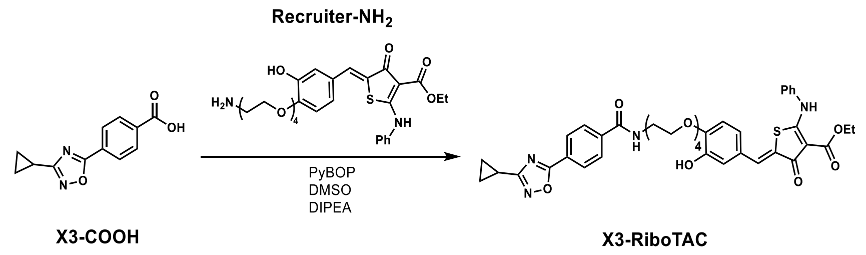


**Scheme S3. Synthesis of X3-RiboTAC.**

**Synthesis of X3-RiboTAC.** **X3-RiboTAC** was synthesized by following the same procedure as described in Scheme S1, except **X3-COOH** was used instead of **X1-COOH**. The product **X3-RiboTAC** (21% yield) was purified by HPLC as described in the **General Synthetic Methods**. $^1$H NMR (600 MHz, MeOD) δ 8.02 (d, J = 8.6 Hz, 2H), 7.90 (d, J = 8.6 Hz, 2H), 7.55 – 7.49 (m, 3H), 7.45 (s, 3H), 6.87 (s, 3H), 4.34 (d, J = 7.1 Hz, 2H), 4.10 (d, J = 4.4 Hz, 2H), 3.83 – 3.78 (m, 2H), 3.63 (dd, J = 4.8, 1.2 Hz, 10H), 3.54 (t, J = 5.5 Hz, 2H), 2.14 – 2.05 (m, 1H), 1.37 (t, J = 7.1 Hz, 3H), 1.08 (d, J = 8.3 Hz, 2H), 1.05 – 1.00 (m, 2H). $^{13}$C NMR (151 MHz, MeOD) δ 182.89, 176.42, 174.31, 174.29, 173.05, 173.03, 167.36, 165.69, 148.68, 146.87, 137.92, 137.29, 131.65, 129.60, 129.58, 129.57, 129.54, 129.52, 129.50, 129.49, 129.47, 127.93, 127.78, 127.73, 127.69, 127.67, 127.66, 127.65, 127.60, 127.58, 127.55, 126.93, 126.32, 124.69, 124.57, 123.41, 116.12, 112.74, 97.52, 70.22, 70.17, 70.12, 69.91, 69.13, 69.02, 67.82, 59.98, 48.46, 48.18, 39.73, 39.02, 36.52, 13.45, 6.89, 6.17. HRMS calculated for $[C_{40}H_{43}N_4O_{10}S]^+$ 771.2700, found 771.2699.

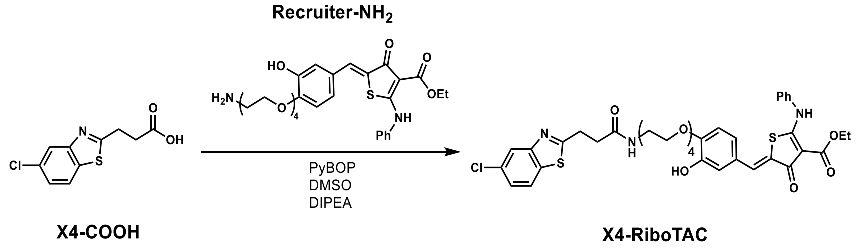


**Scheme S4. Synthesis of X4-RiboTAC.**

**Synthesis of X4-RiboTAC.** **X4-RiboTAC** was synthesized by following the same procedure as described in Scheme S1, except **X4-COOH** was used instead of **X1-COOH**. The product **X4-RiboTAC** (19% yield) was purified by HPLC as described in the **General**

**Synthetic Methods**. $^1$H NMR (600 MHz, DMSO) δ 11.25 (s, 1H), 8.09 – 8.02 (m, 2H), 7.98 (d, J = 2.1 Hz, 1H), 7.58 – 7.47 (m, 5H), 7.47 – 7.41 (m, 2H), 7.04 – 6.93 (m, 3H), 4.28 (q, J = 7.1 Hz, 2H), 4.13 – 4.09 (m, 2H), 3.75 – 3.71 (m, 2H), 3.57 (dd, J = 5.9, 3.7 Hz, 3H), 3.51 (dd, J = 5.9, 3.6 Hz, 2H), 3.50 – 3.44 (m, 4H), 3.38 (t, J = 5.8 Hz, 2H), 3.31 (d, J = 14.6 Hz, 2H), 3.20 (q, J = 5.8 Hz, 2H), 2.66 (t, J = 7.3 Hz, 2H), 1.30 (t, J = 7.1 Hz, 3H). $^{13}$C NMR (151 MHz, DMSO) δ 181.40, 175.79, 174.08, 170.82, 165.35, 154.01, 149.20, 147.50, 137.96, 134.07, 131.23, 130.51, 130.10, 128.64, 126.79, 126.05, 125.36, 125.10, 123.98, 123.54, 122.04, 116.34, 114.19, 97.35, 70.38, 70.25, 70.17, 70.04, 69.55, 69.31, 68.40, 59.98, 40.88, 39.13, 34.31, 29.84, 14.87. HRMS calculated for $[C_{38}H_{41}ClN_3O_9S_2]^+$ 728.1967, found 728.1970.

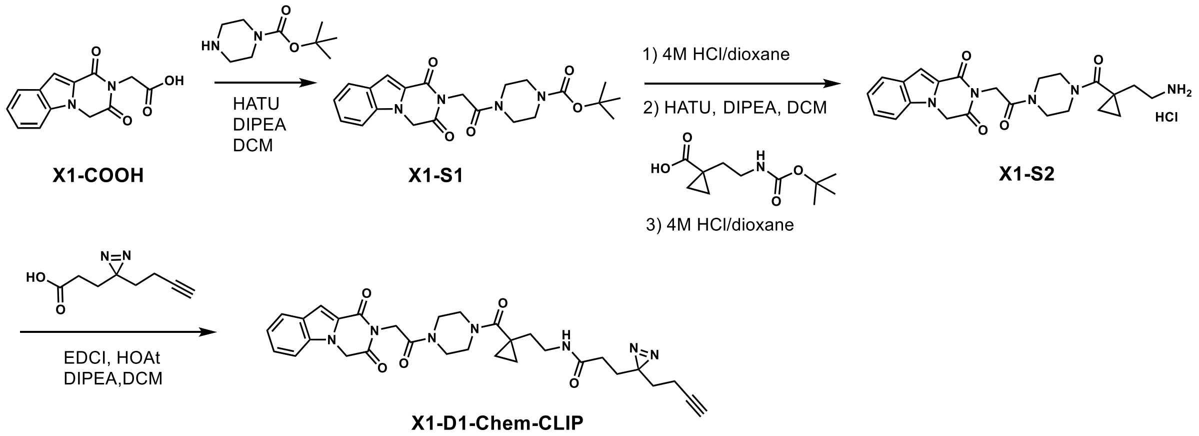


**Scheme S5. Synthesis of X1-D1-Chem-CLIP.**

**Synthesis of X1-D1-Chem-CLIP.** A mixture of **X1-COOH** (50 mg, 0.2 mmol), DCM (5 mL), HATU (103 mg, 0.28 mmol), DIPEA (84 µL, 0.5 mmol) and tert-butyl piperazine-1-carboxylate (50 mg, 0.28 mmol) was stirred at room temperature overnight. The product **X1-S1** was isolated by silica column chromatography (DCM:MeOH increasing from 1:0 to 9:1) and dried under vacuum (66 mg, 80% yield). The isolated **X1-S1** (60 mg, 0.14 mmol) was added 4M HCl/dioxane (1 mL) and stirred at room temperature overnight, followed by drying under vacuum, added EtOAc and triturate. The precipitate was collected by filtration, dried under reduced pressure (51 mg, quantitative yield). A portion of the intermediate (23 mg, 0.06 mmol) was suspended in 2 mL of DCM, followed by the addition of HATU (36 mg, 0.095 mmol), DIPEA (33 μL, 0.19 mmol), and 1-(2-((tert-butoxycarbonyl)amino)ethyl)cyclopropane-1-carboxylic acid (22 mg, 0.95 mmol). The reaction was stirred at room temperature overnight and purified by silica column chromatography (DCM:MeOH increasing from 1:0 to 9:1) and dried under vacuum. The isolated (13 mg, 38% yield) was added 4M HCl/dioxane (1 mL) and stirred at room temperature overnight, followed by drying under vacuum, added EtOAc and triturate. The precipitate was collected by filtration, dried under reduced pressure and get **X1-S2** (9.0 mg, 38% yield). Without further purification, the solid intermediate **X1-S2** (4 mg, 0.01 mmol) was suspended in 2 mL of DCM, followed by the addition of EDCI (3.2 mg, 0.02 mmol), HOAt (2.3 mg, 0.02 mmol), DIPEA (7 μL, 0.04 mmol), and 3-(3-(but-3-yn-1-yl)-3H-diazirin-3-yl)propanoic acid (2.8 mg, 0.02 mmol). The reaction was stirred at room temperature overnight and purified by silica column chromatography (DCM:MeOH increasing from 1:0 to 9:1) and dried under vacuum to obtain the final product **X1-D1-Chem-CLIP** (3.5 mg, 71% yield). $^{1}$H NMR (400 MHz, CDCl3) δ 7.78 (d, J = 8.1 Hz, 1H), 7.49 (s, 1H), 7.45 (dd, J = 8.4, 7.0 Hz, 1H), 7.35 (d, J = 8.4 Hz, 1H), 7.29 – 7.23 (m, 3H), 6.23 (s, 1H), 5.14 (s, 2H), 4.86 (s, 2H), 3.81 – 3.55 (m, 9H), 3.37 (q, J = 6.1 Hz, 2H), 2.07 – 1.99 (m, 4H), 1.95 – 1.81 (m, 5H), 1.26 (s, 4H), 0.98 (q, J = 4.6 Hz, 2H), 0.69 (q, J = 4.6 Hz, 2H). $^{13}$C NMR (151 MHz, CDCl3) δ 172.55, 170.98, 166.32, 164.41, 158.03, 137.10, 127.27, 126.46, 124.09, 123.57, 122.09, 109.88, 109.23, 82.77, 69.26, 46.37, 44.80, 42.19, 40.36, 38.13, 35.66, 32.43, 30.28, 29.71, 29.33, 28.20, 27.94, 23.48, 22.70, 14.13, 13.32, 11.67. HRMS calculated for $[C_{31}H_{36}N_7O_5]^+$ 586.2778, found 586.2772.

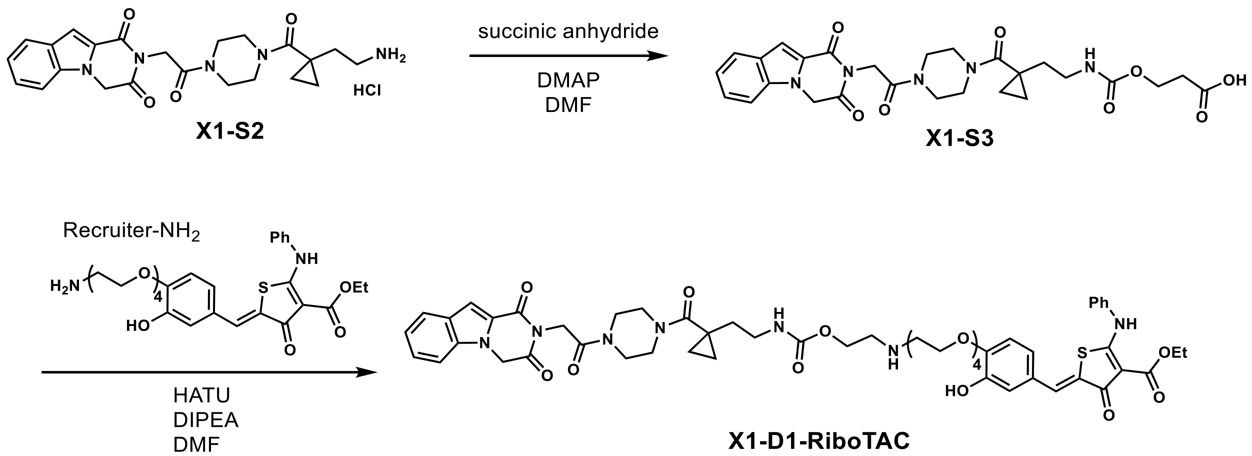


**Scheme S6. Synthesis of X1-D1-RiboTAC.**

**Synthesis of X1-D1-RiboTAC.** **X1-S2** was synthesized as described in Scheme S5. A mixture of **X1-S2** (7 mg, 0.015 mmol), succinic anhydride (1.6 mg, 0.016 mmol), DMAP (2.2 mg, 0.018 mmol) and DMF (1 mL) was stirred at room temperature for 5 h to obtain the intermediate **X1-S3** (not isolated) as detected by LC-MS. The final step of amide coupling with **Recruiter-NH$_2$** was completed by following the same procedure as described in Scheme S1, except for using HATU and DMF, and performing extraction and column purification in the workup, to afford the final product **X1-D1-RiboTAC** (2.2 mg, 14% yield). $^1$H NMR (400 MHz, CDCl3) δ 11.48 (s, 1H), 7.76 (d, J = 8.1 Hz, 1H), 7.71 (s, 1H), 7.57 – 7.30 (m, 10H), 7.12 (d, J = 2.2 Hz, 1H), 7.02 (d, J = 2.2 Hz, 1H), 6.94 (s, 1H), 6.87 (d, J = 8.3 Hz, 1H), 6.51 (s, 1H), 5.12 (s, 2H), 4.82 (s, 2H), 4.39 (q, J = 7.1 Hz, 2H), 4.19 (d, J = 4.9 Hz, 2H), 3.87 (d, J = 5.0 Hz, 2H), 3.80 – 3.49 (m, 23H), 3.44 (d, J = 5.4 Hz, 2H), 3.31 (d, J = 6.4 Hz, 3H), 2.49 (dd, J = 13.3, 5.4 Hz, 4H), 1.43 (t, J = 7.1 Hz, 4H), 1.25 (s, 13H), 0.91 (d, J = 26.1 Hz, 6H), 0.65 (s, 2H). $^{13}$C NMR (151 MHz, CDCl3) δ 182.21, 176.10, 172.40, 172.12, 166.99, 166.31, 164.46, 158.01, 147.90, 147.10, 137.20, 137.08, 131.56, 129.91, 128.08, 127.75, 127.26, 126.39, 125.56, 124.14, 124.07, 123.52, 123.17, 122.04, 116.99, 113.26, 109.90, 109.10, 97.93, 70.52, 70.44, 70.31, 70.07, 69.33, 68.35, 60.60, 46.36, 44.78, 42.13, 40.39, 39.22, 38.26, 35.77, 31.94, 31.60, 31.26, 29.71, 23.48, 22.71, 14.46, 14.13, 11.60. HRMS calculated for $[C_{55}H_{64}N_7O_{14}S]^+$ 1078.4232, found 1078.4222.

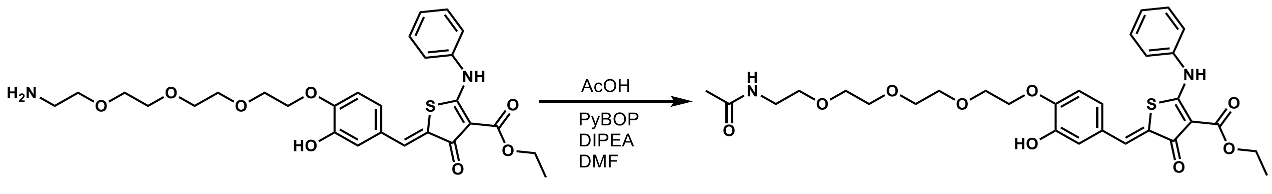


**Scheme S7. Synthesis of Ac-Recruiter.**

**Synthesis of Ac-Recruiter. Ac-Recruiter** was synthesized as described in previously.[16] In brief, AcOH (0.2 mg, 0.004 mmol) was dissolved in 0.05 mL of DMF followed by the addition of PyBOP (3.73 mg, 0.007 mmol) and DIPEA (2.5 µL, 0.014 mmol). The reaction was stirred at room temperature for 10 min followed by the addition of **Recruiter-NH₂** (3 mg, 0.005 mmol). The reaction was stirred at room temperature for 16 h and then diluted by adding 1 mL of MeOH containing 0.1% (v/v) TFA. The product **Ac-Recruiter** (84% yield) was purified by HPLC as described in the **General Synthetic Methods**. $^{1}$H NMR (400 MHz, DMSO) δ 11.24 (s, 1H), 9.43 (s, 1H), 7.87 (s, 1H), 7.63 – 7.39 (m, 6H), 7.08 – 6.93 (m, 3H), 4.28 (q, *J* = 7.1 Hz, 2H), 4.11 (dd, *J* = 5.7, 3.6 Hz, 2H), 3.74 (dd, *J* = 5.8, 3.6 Hz, 2H), 3.58 (m, 2H), 3.49 (m, 2H), 3.37 (t, *J* = 5.9 Hz, 2H), 3.16 (q, *J* = 5.8 Hz, 2H), 1.78 (s, 3H), 1.29 (t, *J* = 7.1 Hz, 3H).

### $^1$H NMR, $^{13}$C NMR, Analytical HPLC, and HRMS Spectra

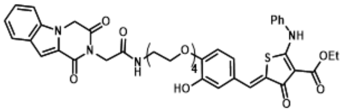


**X1-RiboTAC**

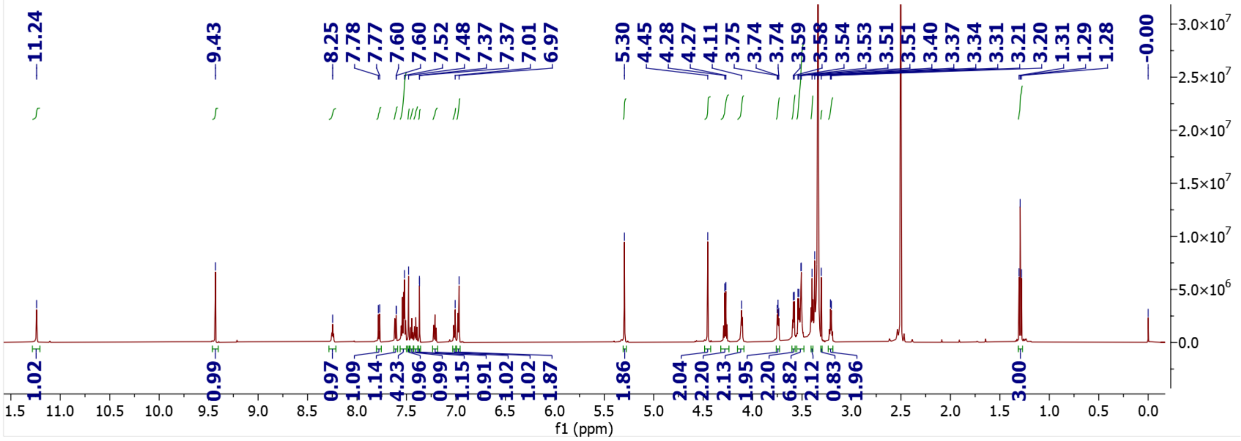


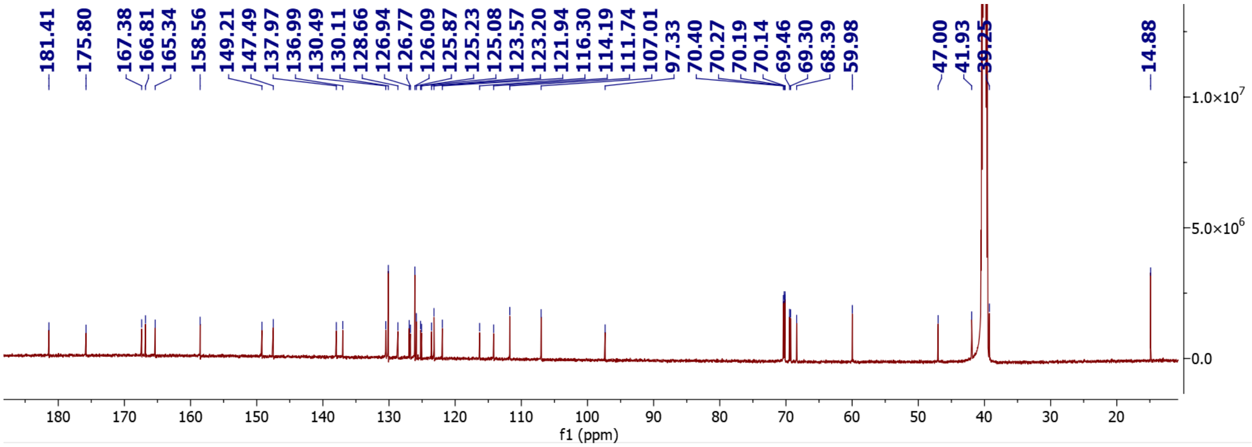


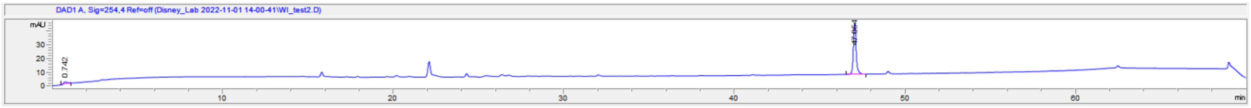


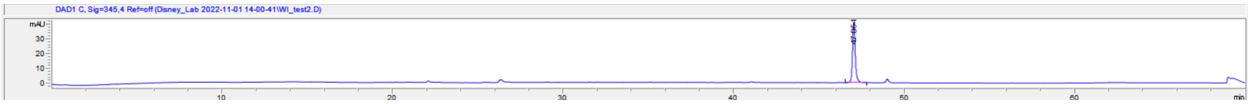


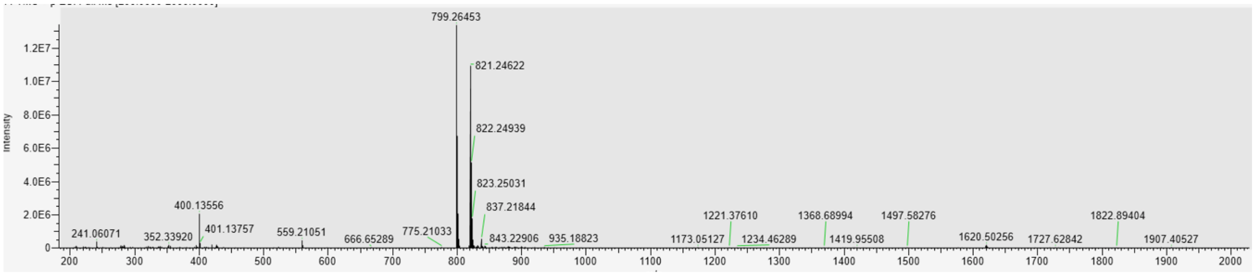

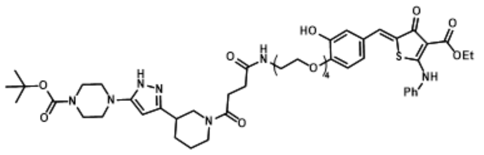
HO
O
O
OEt
NH
Ph
S
H
N
O
4
O
N
N
HN
N
N
O
O
O

**X2-RiboTAC**

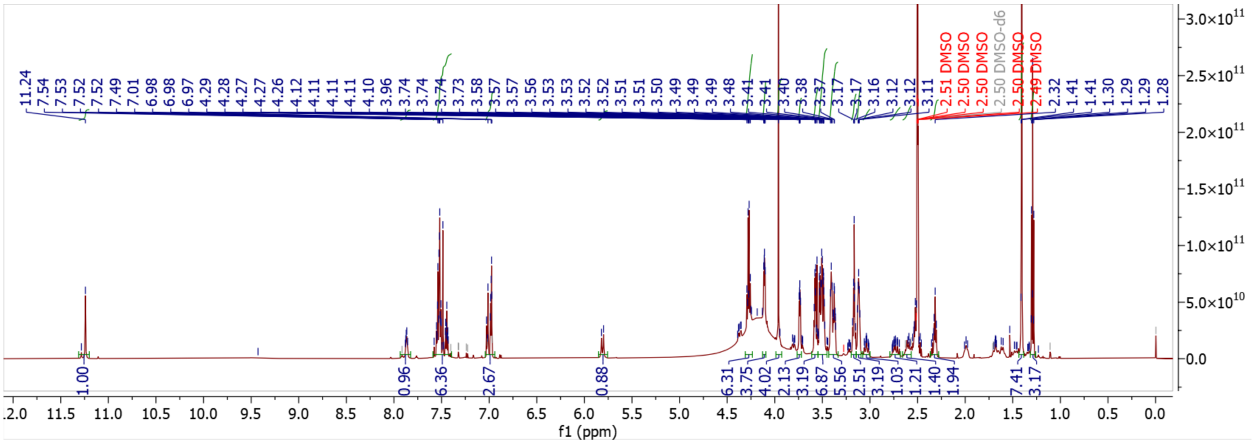
11.24
7.54
7.53
7.52
7.52
7.49
7.01
6.98
6.98
6.97
4.29
4.28
4.27
4.26
4.12
4.11
4.11
4.11
4.10
3.96
3.74
3.74
3.74
3.73
3.58
3.57
3.57
3.56
3.53
3.53
3.52
3.52
3.51
3.51
3.50
3.49
3.49
3.48
3.41
3.41
3.40
3.38
3.37
3.17
3.17
3.16
3.12
3.12
3.11
2.51 DMSO
2.50 DMSO
2.50 DMSO
2.50 DMSO-d6
2.50 DMSO
2.49 DMSO
2.32
1.41
1.41
1.30
1.29
1.29
1.28
1.00
0.96
6.36
2.67
0.88
6.31
3.75
4.02
2.13
3.19
6.87
5.56
2.51
3.19
1.03
1.21
1.40
1.94
7.41
3.17
f1 (ppm)

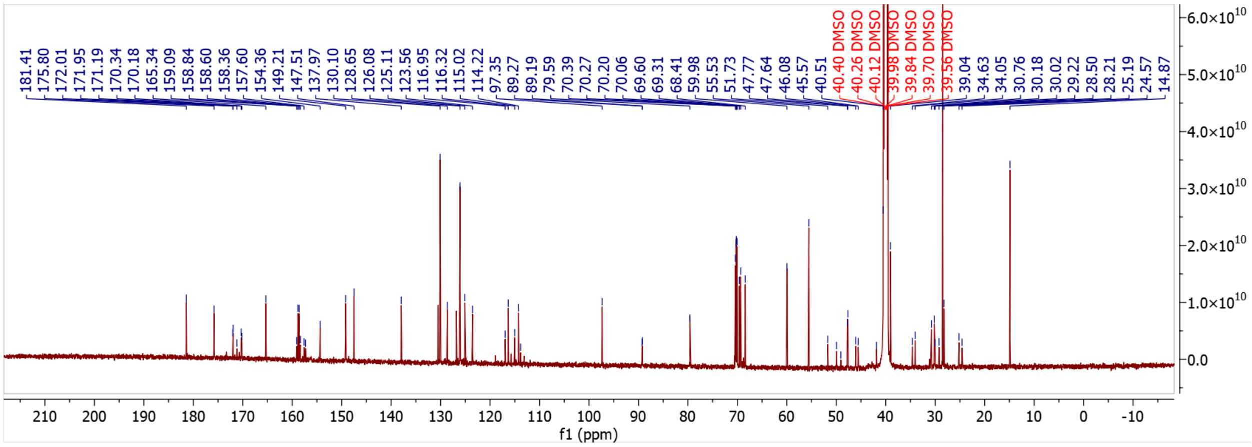
181.41
175.80
172.01
171.95
171.19
170.34
170.18
165.34
159.09
158.84
158.60
158.36
157.60
154.36
149.21
147.51
137.97
130.10
128.65
126.08
125.11
123.56
116.95
116.32
115.02
114.22
97.35
89.27
89.19
79.59
70.39
70.27
70.20
70.06
69.60
69.31
68.41
59.98
55.53
51.73
47.77
47.64
46.08
45.57
40.51
40.40 DMSO
40.26 DMSO
40.12 DMSO
39.98 DMSO
39.84 DMSO
39.70 DMSO
39.56 DMSO
39.04
34.63
34.05
30.76
30.18
30.02
29.22
28.50
28.21
25.19
24.57
14.87
f1 (ppm)

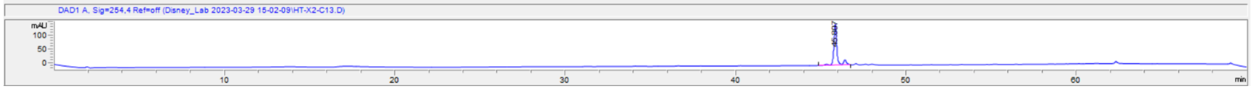
DAD1 A, Sig=254,4 Ref=off (Disney_Lab 2023-03-29 15-02-09\HT-X2-C13.D)
mAU
min

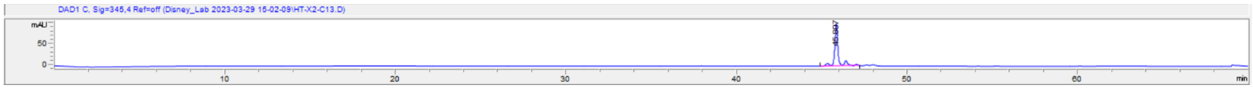
DAD1 C, Sig=345,4 Ref=off (Disney_Lab 2023-03-29 15-02-09\HT-X2-C13.D)
mAU
min

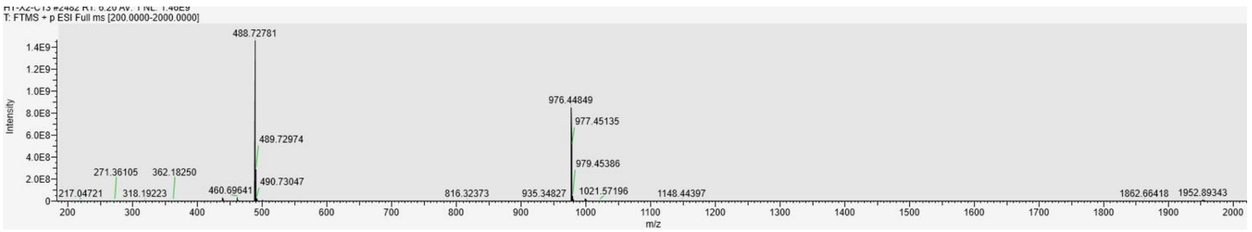
T: FTMS + p ESI Full ms [200.0000-2000.0000]
488.72781
489.72974
490.73047
976.44849
977.45135
979.45386
271.36105
362.18250
217.04721
318.19223
460.69641
816.32373
935.34827
1021.57196
1148.44397
1862.66418
1952.89343
Intensity
m/z

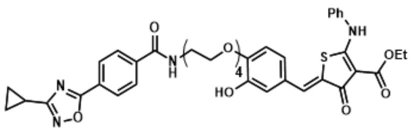

Ph
NH
S
OEt
O
O
N
H
4
HO
N
N-O


X3-RiboTAC

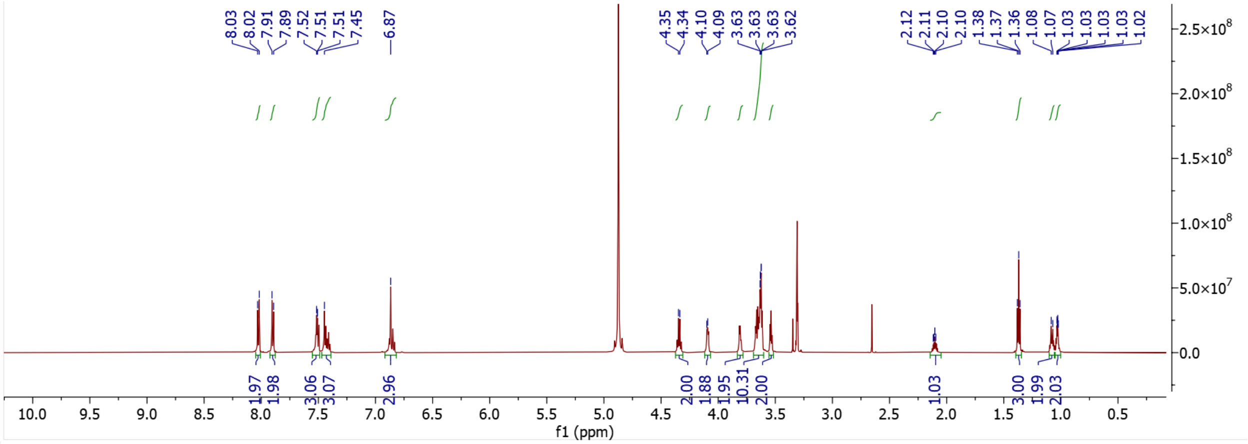

8.03
8.02
7.91
7.89
7.52
7.51
7.51
7.45
6.87
4.35
4.34
4.10
4.09
3.63
3.63
3.63
3.62
2.12
2.11
2.10
2.10
1.38
1.37
1.36
1.08
1.07
1.03
1.03
1.03
1.03
1.02
1.97
1.98
3.06
3.07
2.96
2.00
1.88
1.95
10.31
2.00
1.03
3.00
1.99
2.03
f1 (ppm)


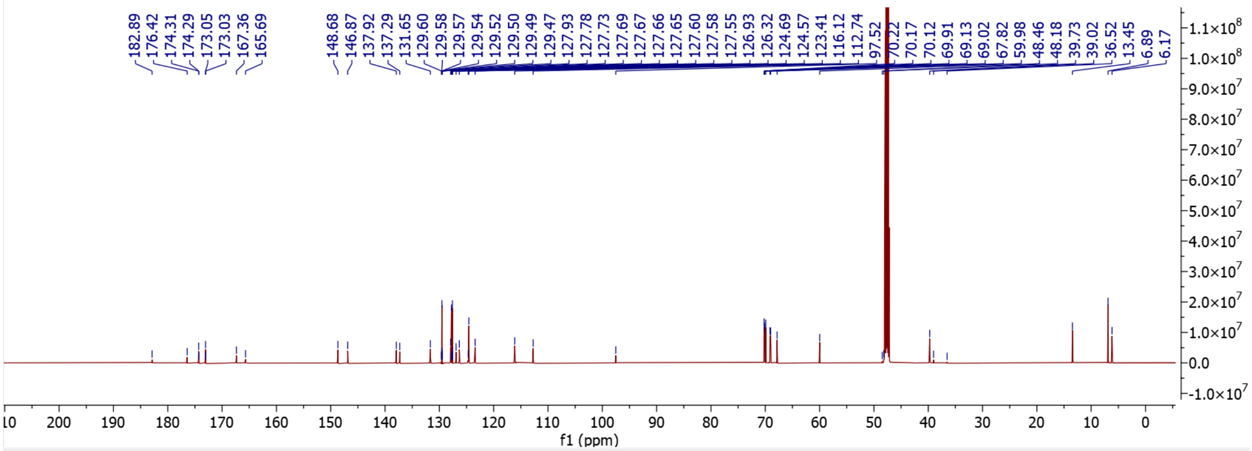

182.89
176.42
174.31
174.29
173.05
173.03
167.36
165.69
148.68
146.87
137.92
137.29
131.65
129.60
129.58
129.57
129.54
129.52
129.50
129.49
129.47
127.93
127.78
127.73
127.69
127.67
127.66
127.65
127.60
127.58
127.55
126.93
126.32
124.69
124.57
123.41
116.12
112.74
97.52
70.22
70.17
70.12
69.91
69.13
69.02
67.82
59.98
48.46
48.18
39.73
39.02
36.52
13.45
6.89
6.17
f1 (ppm)


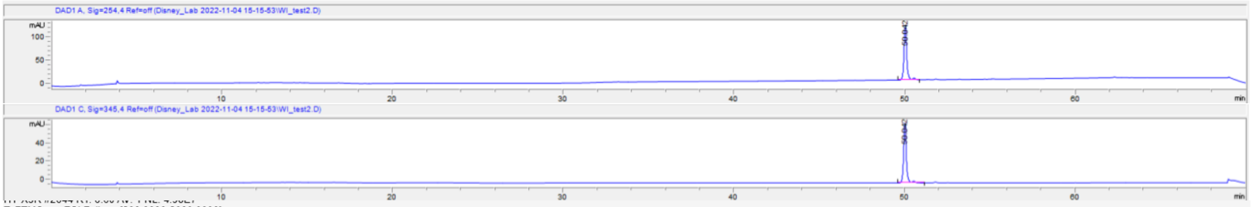

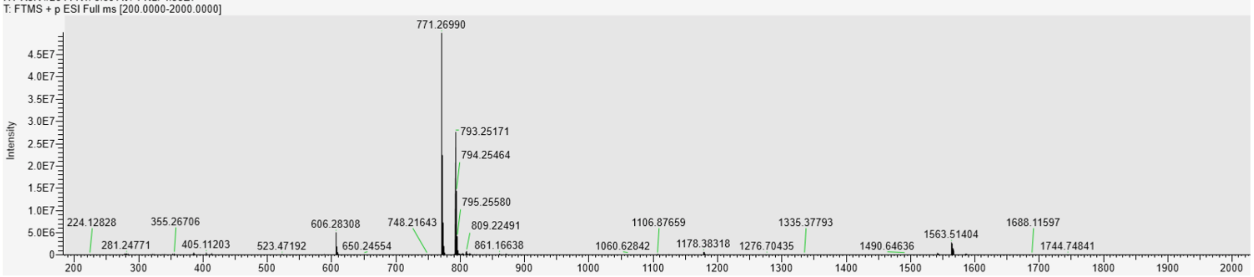

T: FTMS + p ESI Full ms [200.0000-2000.0000]
771.26990
793.25171
794.25464
795.25580
809.22491
861.16638
748.21643
606.28308
650.24554
523.47192
405.11203
355.26706
281.24771
224.12828
1106.87659
1060.62842
1178.38318
1276.70435
1335.37793
1490.64636
1563.51404
1688.11597
1744.74841
Intensity

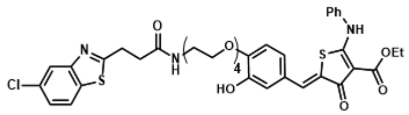
Ph
NH
O
N
S
OEt
Cl
N
H
O
4
HO
O
S
O

**X4-RiboTAC**

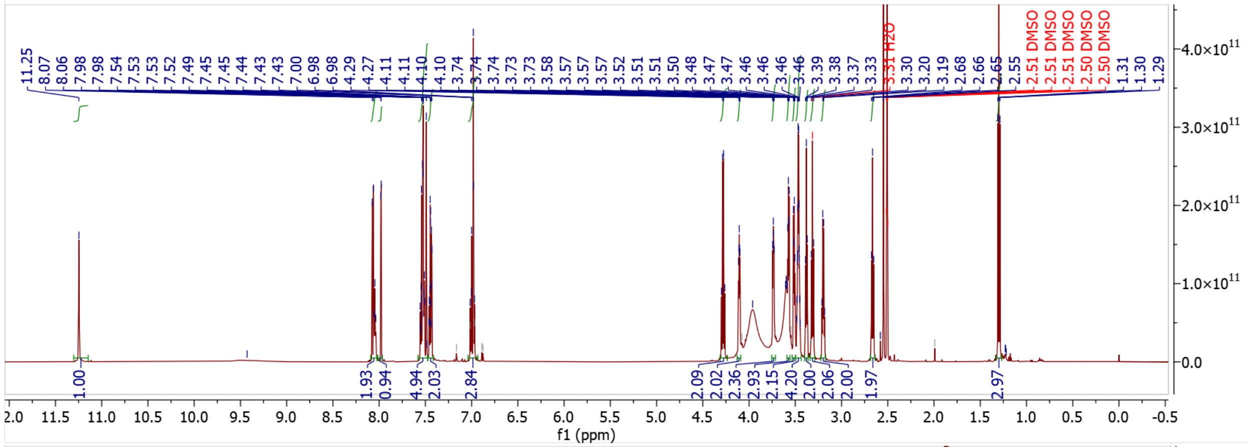
11.25
8.07
8.06
7.98
7.98
7.54
7.53
7.52
7.49
7.45
7.45
7.44
7.43
7.43
7.00
6.98
6.98
4.29
4.27
4.11
4.11
4.10
4.10
3.74
3.74
3.74
3.73
3.73
3.58
3.57
3.57
3.57
3.52
3.51
3.51
3.50
3.48
3.47
3.47
3.46
3.46
3.46
3.46
3.39
3.38
3.37
3.33
3.31 H2O
3.30
3.20
3.19
2.68
2.66
2.65
2.55
2.51 DMSO
2.51 DMSO
2.51 DMSO
2.50 DMSO
2.50 DMSO
1.31
1.30
1.29
1.00
1.93
0.94
4.94
2.03
2.84
2.09
2.02
2.36
2.93
2.15
4.20
2.00
2.06
2.00
1.97
2.97
f1 (ppm)

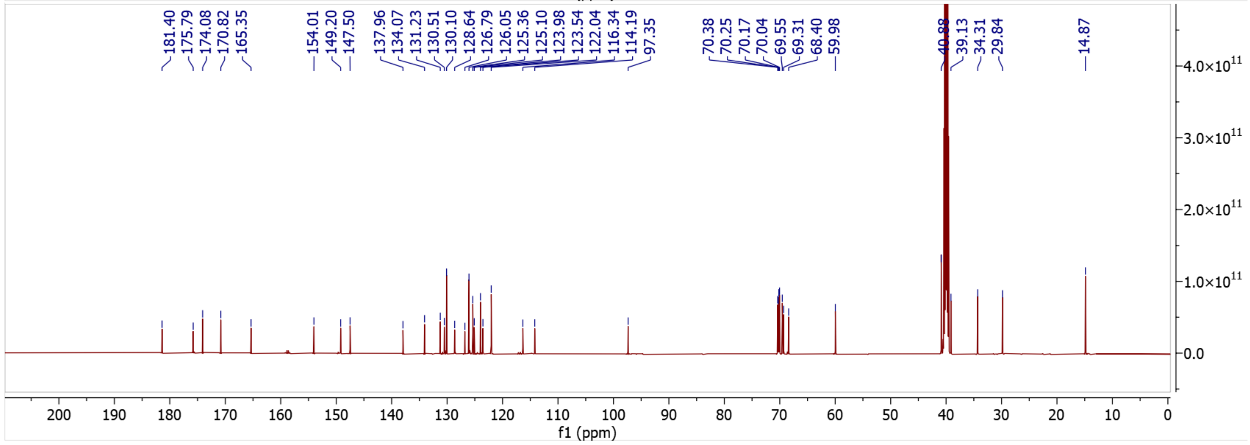
181.40
175.79
174.08
170.82
165.35
154.01
149.20
147.50
137.96
134.07
131.23
130.51
130.10
128.64
126.79
126.05
125.36
125.10
123.98
123.54
122.04
116.34
114.19
97.35
70.38
70.25
70.17
70.04
69.55
69.31
68.40
59.98
39.13
34.31
29.84
14.87
f1 (ppm)

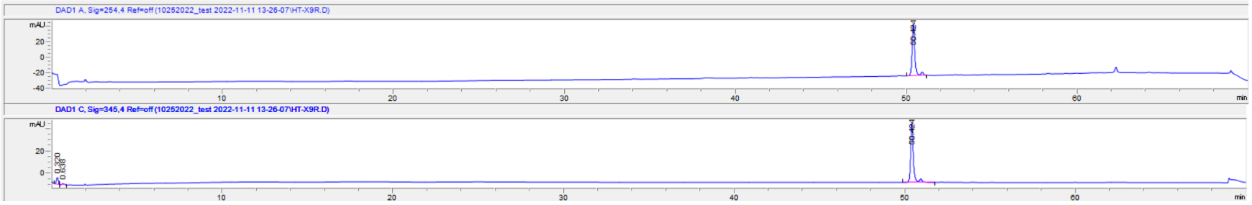
DAD1 A, Sig=254,4 Ref=off (10252022_test 2022-11-11 13-26-07\HT-X9R.D)
DAD1 C, Sig=345,4 Ref=off (10252022_test 2022-11-11 13-26-07\HT-X9R.D)
mAU
min

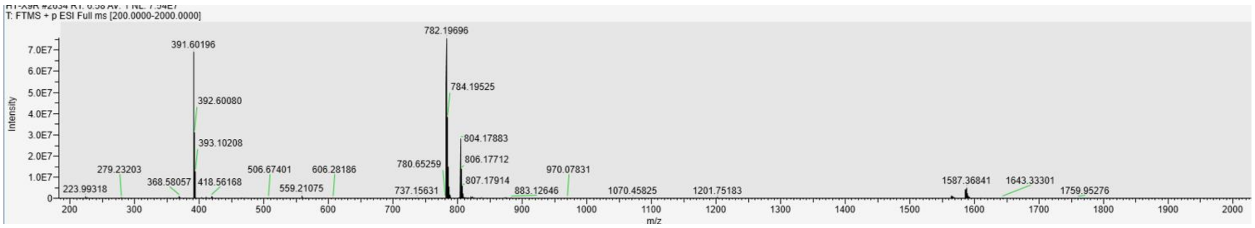
T: FTMS + p ESI Full ms [200.0000-2000.0000]
782.19696
391.60196
784.19525
392.60080
393.10208
804.17883
806.17712
807.17914
279.23203
368.58057
418.56168
506.67401
606.28186
559.21075
780.65259
737.15631
970.07831
883.12646
223.99318
1070.45825
1201.75183
1587.36841
1643.33301
1759.95276
Intensity
m/z

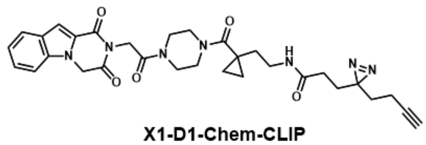

**X1-D1-Chem-CLIP**

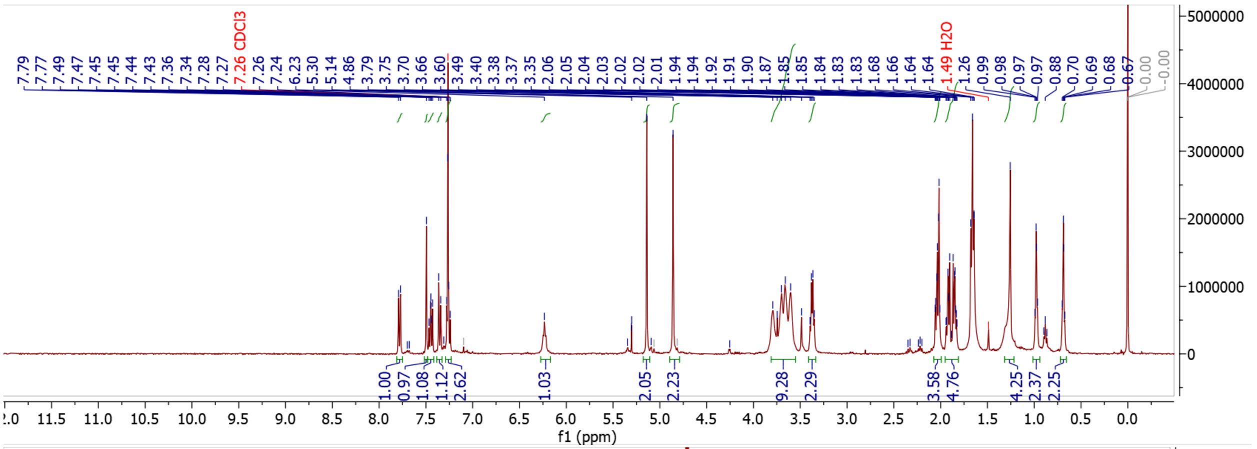

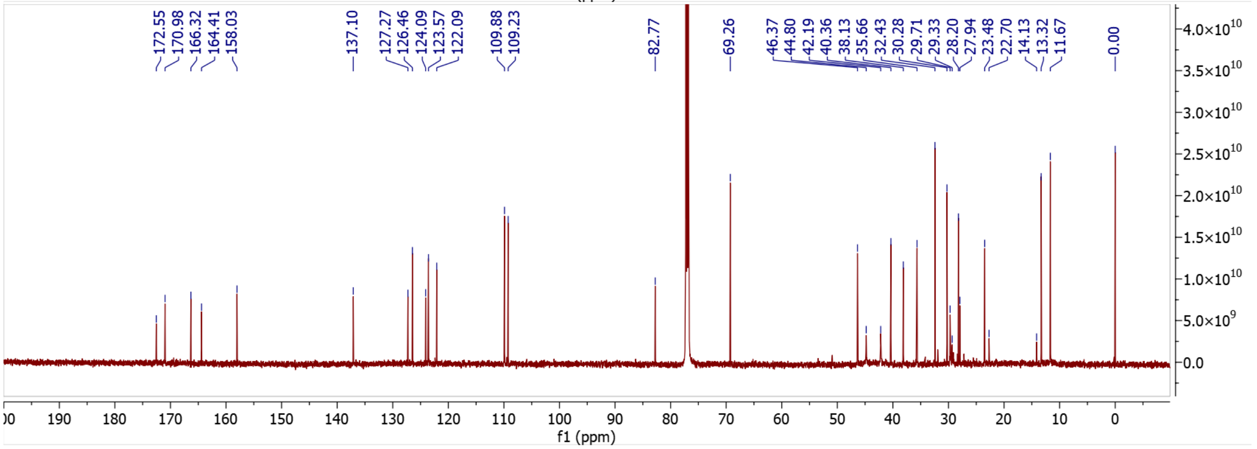

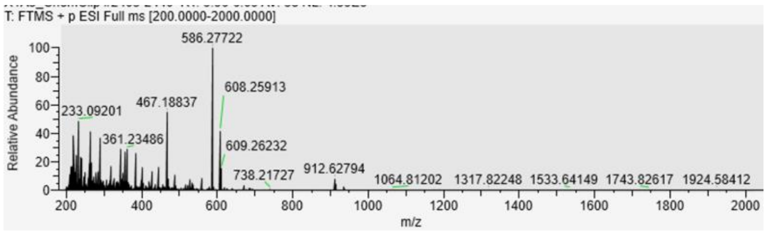

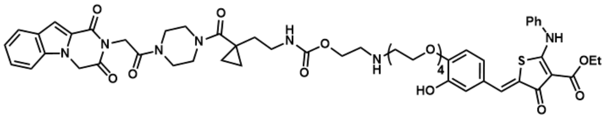


**X1-D1-RiboTAC**

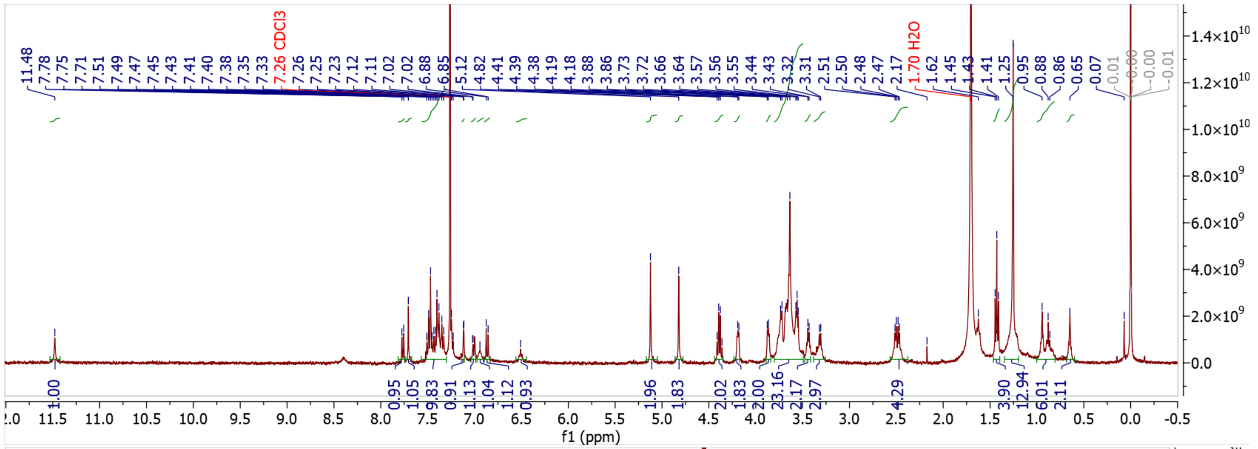


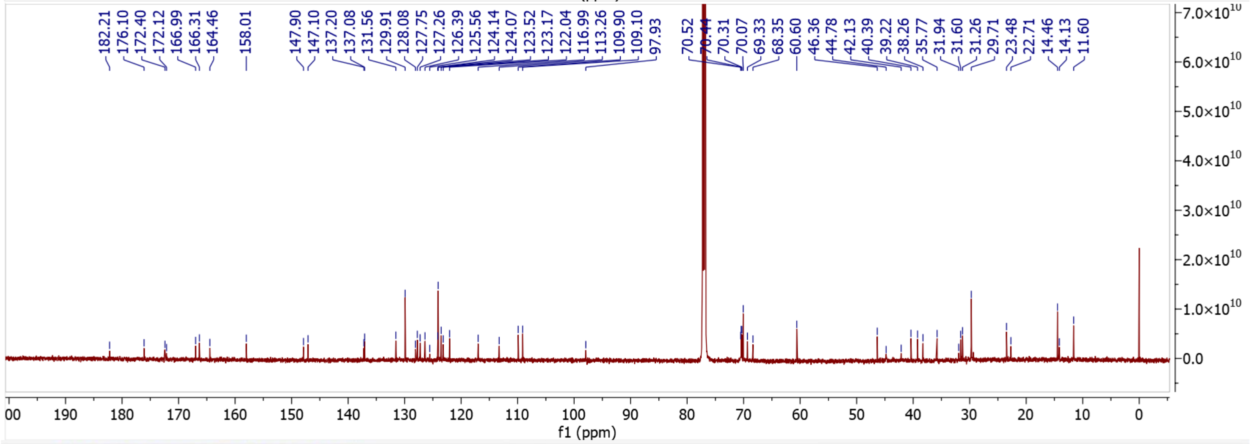


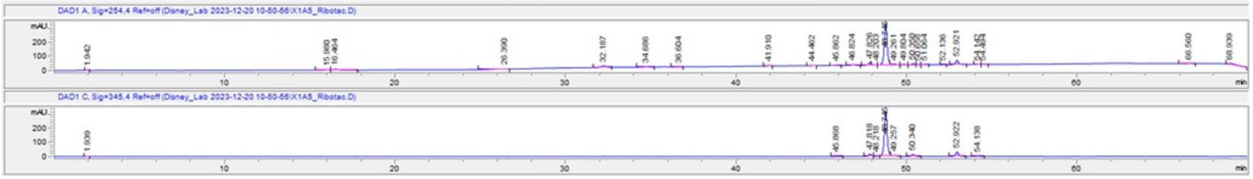


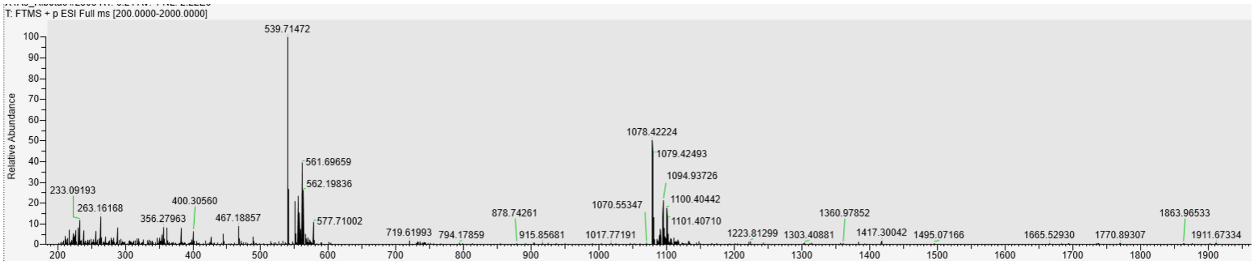

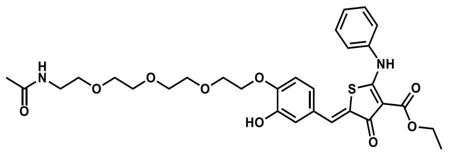

**Ac-Recruiter**

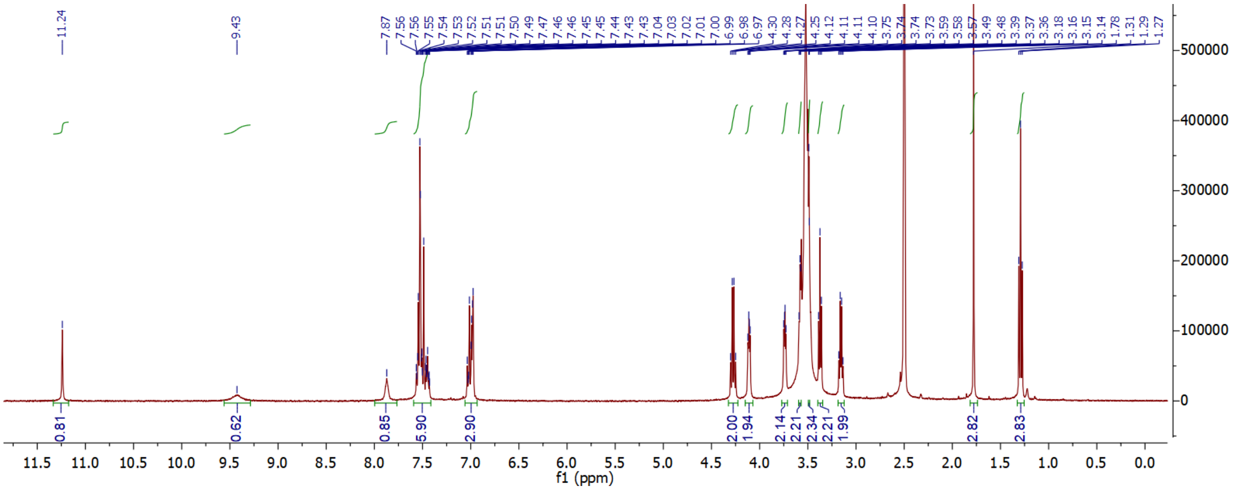

(7) Bouhamdani, N.; Comeau, D.; Turcotte, S. A Compendium of Information on the

Lysosome. *Front Cell Dev Biol* **2021**, *9*, 798262. DOI: 10.3389/fcell.2021.798262.

(13) McKnight, P. E.; Najab, J. Mann-Whitney U Test. In *The Corsini Encyclopedia of Psychology*, pp 1-1.

(14) Szulc, N. A.; Mackiewicz, Z.; Bujnicki, J. M.; Stefaniak, F. fingeRNAt—A novel tool for high-throughput analysis of nucleic acid-ligand interactions. *PLoS Comput. Biol* **2022**, *18* (6), e1009783.

(15) Costales, M. G.; Aikawa, H.; Li, Y.; Childs-Disney, J. L.; Abegg, D.; Hoch, D. G.; Pradeep Velagapudi, S.; Nakai, Y.; Khan, T.; Wang, K. W.; et al. Small-molecule targeted recruitment of a nuclease to cleave an oncogenic RNA in a mouse model of metastatic cancer. *Proc. Natl. Acad. Sci. U.S.A.* **2020**, *117* (5), 2406-2411. DOI: 10.1073/pnas.1914286117.

(16) Tong, Y.; Lee, Y.; Liu, X.; Childs-Disney, J. L.; Suresh, B. M.; Benhamou, R. I.; Yang, C.; Li, W.; Costales, M. G.; Haniff, H. S.; et al. Programming inactive RNA-binding small molecules into bioactive degraders. *Nature* **2023**, *618* (7963), 169-179. DOI: 10.1038/s41586-023-06091-8.
